# The sensory shark: high-quality morphological, genomic and transcriptomic data for the small-spotted catshark *Scyliorhinus canicula* reveal the molecular bases of sensory organ evolution in jawed vertebrates

**DOI:** 10.1101/2024.05.23.595469

**Authors:** H. Mayeur, J. Leyhr, J. Mulley, N. Leurs, L. Michel, K. Sharma, R. Lagadec, J.-M. Aury, O.G. Osborne, P. Mulhair, J. Poulain, S. Mangenot, D. Mead, M. Smith, C. Corton, K. Oliver, J. Skelton, E. Betteridge, J. Dolucan, O. Dudchenko, A.D. Omer, D. Weisz, E.L. Aiden, S. McCarthy, Y. Sims, J. Torrance, A. Tracey, K. Howe, T Baril, A. Hayward, C. Martinand-Mari, S. Sanchez, T. Haitina, K. Martin, S.I. Korsching, S. Mazan, M. Debiais-Thibaud

## Abstract

Cartilaginous fishes (chimaeras and elasmobranchs -sharks, skates and rays) hold a key phylogenetic position to explore the origin and diversifications of jawed vertebrates. Here, we report and integrate reference genomic, transcriptomic and morphological data in the small-spotted catshark *Scyliorhinus canicula* to shed light on the evolution of sensory organs. We first characterise general aspects of the catshark genome, confirming the high conservation of genome organisation across cartilaginous fishes, and investigate population genomic signatures. Taking advantage of a dense sampling of transcriptomic data, we also identify gene signatures for all major organs, including chondrichthyan specializations, and evaluate expression diversifications between paralogs within major gene families involved in sensory functions. Finally, we combine these data with 3D synchrotron imaging and *in situ* gene expression analyses to explore chondrichthyan-specific traits and more general evolutionary trends of sensory systems. This approach brings to light, among others, novel markers of the ampullae of Lorenzini electro-sensory cells, a duplication hotspot for crystallin genes conserved in jawed vertebrates, and a new metazoan clade of the Transient-receptor potential (TRP) family. These resources and results, obtained in an experimentally tractable chondrichthyan model, open new avenues to integrate multiomics analyses for the study of elasmobranchs and jawed vertebrates.

## 1. INTRODUCTION

Cartilaginous fishes, which comprise elasmobranchs (i.e. sharks, skates and rays) and holocephalans (chimaeras) (Janvier 1996), are a group of major interest for vertebrate evolutionary biologists. As sister group to bony fishes, which contain all major vertebrate model organisms (e.g. mouse, chicken, *Xenopus*, zebrafish), they occupy an important phylogenetic position for elucidating early evolutionary events at the gnathostome crown node (Kuraku 2021; Tan et al. 2021; Yamaguchi et al. 2021; Zhang et al. 2022; Marlétaz et al. 2023). This position makes them important models for determining genomic bases of phenotypic evolution, especially the origin of ancestral features of jawed vertebrates and subsequent taxon-specific diversification. Such approaches require the availability of high-quality genomic data from across cartilaginous and bony fish diversity, coupled to a commensurate knowledge of the morphological evolution of traits of interest. In this respect, the more than 1000 species of chondrichthyans remain massively under-represented. However, the past few years have seen a growing effort to produce genomic data for this group resulting in a well-assembled genome for a holocephalan, the elephant shark *Callorhinchus milii*, and several chromosome-level assembled genomes for elasmobranch fishes including batoids (smalltooth sawfish *Pristis pectinate*, thorny skate *Amblyraja radiata,* little skate *Leucoraja erinacea*) and selachians (one Carcharhiniform, the small-spotted catshark *Scyliorhinus canicula* (this study); one lamniform, the great white shark *Carcharodon carcharias*; two orectolobiforms, the epaulette shark *Hemiscyllium ocellatum* and the white spotted bamboo shark *Chiloscyllium plagiosum*; one squaliform, the spiny dogfish *Squalus acanthias*), as listed in Table 1. Additional genomes have been sequenced with a lower quality of assembly, and their annotations are also available (Weber et al. 2020; Stanhope et al. 2023; Zhou et al. 2023) notably through the Squalomix project (Hara et al. 2018; Nishimura et al. 2022). Even fewer cartilaginous fishes have been the focus of integrative studies associating genomic or transcriptomic data with different organisational levels from organism to organ, to cell, or successive developmental stages (see Gillis et al. 2022). Reasons for this lack of integrative data include the difficulty of accessing biological material (e.g. protection status for certain species, such as the smalltooth sawfish), biological features (e.g. size and reproduction, such as for the great white shark and whale shark), and more generally the lack of previous physiological/anatomical studies.

**Table 1:** summary of chromosome-level assembly features available in cartilaginous fishes (green: sharks; blue: skates and rays) with focus on the small-spotted catshark *Scyliorhinus canicula* (highlighted in yellow), and in the partially assembled genome of the elephant shark *Callorhinchus milii*. N: number of chromosomes in the haploid genome. Maximum chromosome size as described in GenBank. Reference publications [1] (Read et al. 2017), [2] (Zhang et al. 2020), [3] (Sendell-Price et al. 2023), [4] (Wagner et al. 2023), [5] (Marra et al. 2019), [6] this work, [7] (Yoo et al. 2022; Marlétaz et al. 2023), [8] (Venkatesh et al. 2007; Venkatesh et al. 2014).

| Order | Assembly (NCBI) | Species name |  | N | Genome size | N >40Mb (%) | Maximum chrom. size |
| --- | --- | --- | --- | --- | --- | --- | --- |
| Orectolobiforms | sRhiTyp1.1 | *Rhincodon typus* | [1] | 50 | 2.9Gb | 28 (56%) | 178Mb |
|  | ASM401019v1 | *Chiloscyllium plagiosum* | [2] | 51 | 3.6Gb | 35 (68%) | 156Mb |
|  | sHemOce1.pat.X.c<br>ur | *Hemiscyllium ocellatum* | [3] | 52 + XY | 4.0Gb | 39 (73%) | 167Mb |
| Squaliforms | ASM3039002v1 | *Squalus acanthias* | [4] | 29 | 3.7Gb | 25 (86%) | 266Mb |
| Lamniforms | sCarCar2.pri | *Carcharodon carcharias* | [5] | 40 + XY | 4.3Gb | 25 (60%) | 274Mb |
| Carcharhiniforms | sScyCan1.1 | ***Scyliorhinus canicula*** | **[6]** | **31** | **4.2Gb** | **21 (68%)** | **313Mb** |
| Rajiforms | Leri_hhj_1 | *Leucoraja erinacea* | [7] | 50 | 2.2Gb | 19 (38%) | 170Mb |
|  | sAmbRad1.1.pri | *Amblyraja radiata* |  | 49 | 2.6Gb | 24 (48%) | 193Mb |
| Rhinopristiforms | sPriPec2.1.pri | *Pristis pectinata* |  | 46 + X | 2.3Gb | 20 (42%) | 150Mb |
| Holocephalan | IMCB_Cmil_1.0 | *Callorhynchus milii* | [8] | na | 0.9Mb | na | na |

This work aims at filling this gap, focusing on sensory systems and using the small-spotted catshark *Scyliorhinus canicula*, an emerging model organism, as the reference species. Understanding how sensory systems diversify during evolution is crucial to apprehend the multiple strategies employed by animals to process environmental information and it typically requires a multi-level approach, from gene to cell and organ. The rise of paired sensory organs, including our eyes, nose, and ears, was a key innovation of early vertebrates, emerging alongside a complex, centralised brain and the prototypical ‘new head’ of vertebrates (Gans and Northcutt 1983). Some of these systems were later lost in tetrapods, specifically: (i) the lateral line -a mechanosensory organ in the form of neuromasts located just below the epidermis or in a canal running into the dermis-, and (ii) the electrosensory ampullary organs, also formed of canals, but filled with a specialised secreted hydrogel which has a high conductivity and, as a result, weak electric fields can be detected by sensory cells in the deepest part of the tube (the ampullae). Cartilaginous fishes display the complete set of paired ectodermal sensory organs (eyes, olfactory organs, ears, electrosensory organs, and lateral line; Collin, 2012). However, cartilaginous and bony fish groups diverged over about 420 My ago and cartilaginous fishes diversified into a myriad of forms with a great range of habitats and life history traits (Ebert et al. 2021). This independent evolution is expected to have generated significant variation in their sensory systems, as observed in anatomical aspects of the electrosensory system (reviewed in Collin, 2012 and see Bottaro, 2022; Newton et al., 2019) and olfactory system (Schluessel et al. 2008; Yopak et al. 2015), as well as in sensory cell types such as photoreceptors (Hart et al. 2020; Yamaguchi et al. 2021).

We have focused on the small-spotted catshark *Scyliorhinus canicula* to provide a reference, laying the groundwork for the analysis of this diversity both within cartilaginous fish and at the origin of jawed vertebrates (Coolen et al. 2008). With easy access across developmental stages, a small body size, non-endangered status, high abundance in the North-East Atlantic and easy maintenance in aquaria, this species has become a target elasmobranch model for research in European laboratories for more than 20 years (Mazan et al. 2000; Tanaka et al. 2002; Coolen et al. 2007; O’Neill et al. 2007; Sakamoto et al. 2009; Oulion et al. 2010; Debiais-Thibaud et al. 2011a; Rodríguez-Moldes et al. 2011; Mulley et al. 2014; Tulenko et al. 2016; Redmond et al. 2018; Mayeur et al. 2021). However, until recently, a major limitation was the absence of a chromosome-level genome assembly.

Here, we describe the assembled and thoroughly annotated genome of the small-spotted catshark, a wide series of transcriptomic data from adult tissues and embryonic stages, and cutting-edge 3D morphological data, now allowing integrative studies in this species, from the level of individual sequences to the whole organism. We exemplify how these resources can be combined to better understand the molecular bases of evolution, focusing on paired sensory organs. From the RNAseq data, we identify the most highly expressed genes in each sensory organ. We then consider candidate gene families known to play major roles in sensory organs: opsins and crystallins in the retina and lens respectively; the four main vertebrate olfactory receptor families in the nose; and the transient-receptor potential (TRP) family, essential for diverse sensory functions. For each family we describe the catshark gene repertoire, examine its evolution in relation to other vertebrates, and analyse tissue-specific expression patterns. Sites of expression are interpreted in the light of new 3D anatomical data and histology, facilitating a deeper understanding of the development and organisation of sensory organs in the small-spotted catshark. These integrative data highlight ancestral gnathostome features of the electrosensory system and evolutionary trends in the olfactory and visual system of cartilaginous fishes.

## 2. MATERIALS AND METHODS

### 2.1 Sampling and sequencing

#### RNA-seq sampling and sequencing

RNAs were directly extracted from tissues flash-frozen in liquid nitrogen at the time of their dissection, following the Sigma TRI-reagent® extraction protocol: one whole embryo was used for each standard embryonic stage 12, 22, 24, 26, 30 and 31 (Ballard et al., 1993), and a list of twenty-five different adult tissues were sampled from three individuals (one female, two males), all of these being of Mediterranean origin, fished off the coast of Banyuls-sur-mer, Fr. RNA libraries were sequenced on Illumina NovaSeq 6000 (150bp, paired end). Raw data were deposited in NCBI (Bioproject: PRJEB35946; BioSample accessions: 21694013 to 21694062; see Supplementary Table 1).

#### Mediterranean genome sequencing & annotation

Tissues used for DNA extraction were sampled from the male1 individual in the sample list (Supplementary Table 1). The assembly sScyCan1.1 is based on 63x PacBio data (CLR sequencing), 43x 10X Genomics Chromium data, and BioNano data generated at the Wellcome Sanger Institute, and 17x Hi-C data generated at the Baylor College of Medicine. The assembly process included the following sequence of steps: initial PacBio assembly generation with Falcon-unzip (Chin et al. 2016); retained haplotig identification with purge_dups; 10X based scaffolding with scaff10x; BioNano hybrid-scaffolding; Hi-C-guided scaffolding by the Aiden lab as part of a collaboration with the DNA Zoo Consortium (www.dnazoo.org) using 3D-DNA and Juicebox Assembly Tools (Durand et al. 2016; Dudchenko et al. 2017; Dudchenko et al. 2018); Arrow polishing; and two rounds of FreeBayes polishing(Garrison and Marth 2012). Finally, the assembly was analysed and manually improved using gEVAL (Chow et al. 2016; Howe et al. 2021), with BlobToolKit (Challis et al. 2020) used to detect possible sequence contamination. Chromosome-scale scaffolds confirmed by the Hi-C data were named in order of size. The interactive contact maps for the Hi-C guided genome assembly step can be found at https://www.dnazoo.org/assemblies/scyliorhinus_canicula.

#### Atlantic genome sequencing & assembly

For Illumina libraries sequenced at Genoscope, 250 ng of genomic DNA extracted from an adult individual fished in Roscoff (France; sex unknown) were sonicated using the S2 Covaris instrument (Covaris, Inc., USA). Sheared DNA was used for Illumina library preparation by a semi-automatized protocol. Briefly, end repair, A tailing and Illumina compatible adaptors (BiooScientific) ligation were performed using the SPRIWorks Library Preparation System and SPRI TE instrument (Beckmann Coulter), according to the manufacturer protocol. A 300-600 bp size selection was applied in order to recover the most of fragments. DNA fragments were amplified by 12 cycles PCR using Platinum Pfx Taq Polymerase Kit (Life Technologies) and Illumina adapter-specific primers. Librarie was purified with 0.8x AMPure XP beads (Beckmann Coulter), and size-selected on 1.5% agarose gel around 600 bp. After library profile analysis by Agilent 2100 Bioanalyzer (Agilent Technologies, USA) and qPCR quantification (MxPro, Agilent Technologies, USA), library was sequenced using 101 base-length read chemistry in a paired-end flow cell on the Illumina sequencer (Illumina, USA).

The MP libraries were prepared using the Nextera Mate Pair Sample Preparation Kit (Illumina, San Diego, CA). Briefly, genomic DNA (4 µg) was simultaneously enzymatically fragmented and tagged with a biotinylated adaptor. Tagmented fragments were size selected (3-5; 5-8 and 8-11 Kb) through regular gel electrophoresis, and circularized overnight with a ligase. Linear, non-circularized fragments were digested and circularized DNA was fragmented to 300-1000-bp size range using Covaris E210. Biotinylated DNA was immobilized on streptavidin beads, end-repaired, then 3’-adenylated, and Illumina adapters were added. DNA fragments were PCR-amplified using Illumina adapter-specific primers and then purified. Finally, libraries were quantified by qPCR and libraries profiles were evaluated using an Agilent 2100 bioanalyzer (Agilent Technologies, USA). Each library was sequenced using 100 base-length read chemistry on a paired-end flow cell on the Illumina HiSeq2000 (Illumina, USA).

The reads were submitted to an initial cleaning step to trim sequences and remove low-quality or very short reads, and then corrected using Musket (Liu et al. 2013) with a 17 kmer size. 180 pb paired-ends reads were joined using fastq-join (https://code.google.com/p/ea-utils/wiki/FastqJoin) with default parameters. Assembly and scaffolding of paired end sequences was conducted using CLC (http://www.clcbio.com/products/clc-assembly-cell/) with a 61 pb kmer size and only solving bubbles shorter than 500bp. Mate pair sequences were introduced in additional scaffolding and gap closing steps, respectively using SSPACE (Boetzer et al. 2011) and Gapcloser (Luo et al. 2012).

### 2.2 Genome structure and synteny

#### Species tree inference

In order to compare evolution of genome structure across chondrichthyans, we obtained thirteen genome assemblies from this group labelled as chromosome level assemblies from NCBI. Three representative outgroup species with chromosome level assemblies were also included in the analysis (Supplementary Table 2). Species tree reconstruction of all 16 species was carried out using a data set of 1,005 genes annotated with BUSCO v5.1.2 (Manni et al. 2021) using the Vertebrata gene set. BUSCO genes present in each of the species in our dataset were extracted, aligned, and trimmed using the busco2phylo pipeline (github.com/lstevens17/busco2phylo-nf), which employed MAFFT v7.4 (Katoh 2005) and trimAl v1.4 (Capella-Gutiérrez et al. 2009) for gene alignment and trimming. All trimmed alignments were concatenated into a supermatrix using PhyKIT (Steenwyk et al. 2021), and species tree inference was carried out with IQ-TREE v2.0 (Minh et al. 2020), applying the LG model with a gamma distribution with 4 categories. Species trees were plotted, alongside the inferred genome size and chromosome number per species (inferred from the genome assemblies) using Toytree (Eaton 2020).

#### GC variation across genomes

GC content was calculated at three levels; (i) genome-wide, measured using the complete genome assembly, (ii) coding regions, measured using the nucleotide versions of the trimmed BUSCO gene alignments, and (iii) GC of the third codon position, also inferred using the BUSCO gene alignments. These three values were plotted against the species tree in Supplementary Figure 1 using ggtree (Yu et al. 2017). For each species, GC content of each chromosome above 1Mb in size was plotted against chromosome size, to inspect for patterns of correlation between GC content and chromosome size in Chondrichthyes Supplementary Figure 2.

#### Genome macrosynteny

Macrosynteny of the focal species *Scyliorhinus canicula* was compared to two other species in Chondrichthyes (*Carcharodon carcharias* and *Pristis pectinata*) using the MCscan pipeline in the jcvi software: github.com/tanghaibao/jcvi (Tang et al. 2008). Briefly, this pipeline uses BLAST to carry out a pairwise synteny search between species using the coding sequences and gene locations. Conserved synteny blocks containing 30 genes or more were retained and visualised, with *Scyliorhinus canicula* set as the focal reference species.

### 2.3 Repeat annotation

The sequences of transposable elements (TEs) in the small-spotted catshark genome were annotated with the Earl Grey TE annotation pipeline (version 1.2, github.com/TobyBaril/EarlGrey) as described previously (Baril et al. 2021; Baril and Hayward 2022). In brief, the pipeline was used firstly to identify known TEs from the vertebrate subset of Dfam (release 3.4) and RepBase (release 20,181,026) (Jurka et al. 2005; Hubley et al. 2016). Next, *de novo* TEs were identified and the boundaries of consensus sequences were extended using an automated “BLAST, Extract, Extend” process (Platt et al. 2016). Any redundant sequences were removed from the consensus library before the genome assembly was annotated using a final combined library, which consisted of the known and *de novo* TE libraries. TE annotations were subsequently processed to remove any overlaps and also to defragment broken annotations before the final TE quantification was performed.

### 2.4 Hox cluster analysis

Hox protein sequences from a range of cartilaginous fish (Supplementary Table 3) and humans were downloaded from NCBI, and used as queries in tBLASTn searches of the *Scyliorhinus canicula* gene model dataset (Mayeur et al. 2021) and the genome assembly. We also used chondrichthyan and human protein sequences of the HoxC flanking genes *tspan31*, *ddx23*, *rnd1*, *adcy6*, *smug1*, and *copz1* (Yamaguchi, Uno, et al. 2023) to identify the syntenic region in the *S. canicula* assembly.

### 2.5 Mediterranean and Atlantic population differentiation

To estimate the population differentiation between Mediterranean and Atlantic catshark populations, we used RNA-seq data from three NCBI bioprojects: PRJEB36280 (Mediterranean), PRJNA255185 (North Wales, UK) and PRJNA504730 (English channel, Brittany, France). We used the Nextflow (v.23.10.0; Di Tommaso et al. 2017) f-core rnavar (v.1.1.0; Ewels et al. 2020) pipeline for read mapping and SNP calling, which is based on the GATK RNA-seq variant calling best-practice workflow (github.com/gatk-workflows/gatk4-rnaseq-germline-snps-indels). We used the “--star_twopass” option to run STAR two-pass read mapping (Dobin et al. 2013), and ran read sets of different lengths separately so that optimal STAR indices could be produced for each read length (additionally, the following non-default options were used: --remove_duplicates, --skip_baserecalibration, --skip_variantannotation, -- generate_gvcf). Variant calls from all individuals were then combined and jointly genotyped using the COMBINEGVCFS and GENOTYPEGVCF programmes from the GATK software suite (v.4.2.6.1; McKenna et al. 2010). To produce unbiased estimates of population genetic statistics, we called genotypes at both variant and invariant sites. All sites were filtered to a maximum missing data percentage of 15% (i.e. two individuals missing) and minimum read depth of 5 using VCFtools. Variant and invariant sites were then separated and variant sites were additionally filtered to include only SNPs with minor allele frequency > 0.1 and minimum genotype quality of 30 with VCFtools, before the sites were recombined with the concat function from BCFtools (Li 2011). Fixation index (Weir and Cockerham’s *F_ST_*), absolute divergence (*d_XY_*), and nucleotide diversity for each population (π) were calculated in non-overlapping 1Mb windows with pixy (Korunes and Samuk 2021). Additionally, per-site Weir and Cockerham’s *F_ST_* was calculated using VCFtools. Gene density and GC content were calculated in non-overlapping 1Mb windows using BEDTools (Quinlan and Hall 2010). Outlier SNPs potentially under divergent selection between Atlantic and Mediterranean populations were identified using pcadapt (Luu et al. 2017) with *K* = 2, LD clumping (size = 500, thresh = 0.1) and the “componentwise” method. Genome-wide statistics were visualised with qqman (Turner 2018). Principal Component Analysis (PCA) was performed using PLINK 2.0 (v.2.00a3LM; Chang et al. 2015), after thinning SNPs to one per 100 Kb to reduce the impact of genetic linkage. Population structure was estimated using ADMIXTURE (v.1.3.0; Alexander et al. 2009) using the same thinned set of SNPs as the PCA analysis and 5-fold cross validation. The code used in this section is available at github.com/ogosborne/Scyliorhinus_canicula_popgen.

### 2.6 Reference gene model

The reference gene model ‘ncbi-utrs’ previously published in Mayeur et al. 2021 was used for RNA-seq mapping. Its construction aimed at maximising sequence predictions in 3’UTRs, and it was conducted as follows using custom R and PERL scripts (Supplementary text). First, an "isoform-collapsed" version of gene models was constructed by selecting the most supported isoform of each gene, as defined in NCBI Annotation Release 100 of the *Scyliorhinus canicula* genome (sScyCan1.1 version). All NCBI predicted transcripts were then mapped back by BLASTn (identity=100% and e-value<1E^-06^) to this ‘isoform-collapsed’ reference in order to identify for each gene the isoform with the longest 3’UTR. When detected, the additional 3’UTR sequence was appended to the sequence of the ‘isoform-collapsed’ reference. A second round of 3’UTR expansion was conducted from the resulting gene model set by including divergent isoforms identified with lower BLASTn supports (identity<100% and e-value<1E^-06^). A final iteration of 3’UTR expansion was carried out using a transcriptome assembled from available public NCBI SRA reads as well as locally obtained RNA-seq data as a query (listed in Supplementary Table 4). The transcriptome assembly used in this final step was conducted using DRAP (Cabau et al. 2017) and isoform-collapsed using Corset and SuperTranscripts (Davidson and Oshlack 2014; Davidson et al. 2017). In this final step, the 3’UTR appending step was carried out using only BLASTn hits with 100% identity with the subject, e-value<1E^-06^.

### 2.7 Transcriptome mapping and gene expression ranking

The paired end RNA-seq reads from adult and embryonic *S. canicula* tissue samples were mapped onto the reference gene model ncbi-utrs using BWA-MEM (Li 2013). After sorting and converting the resulting alignments to BAM files using SAMtools, counts for each library were extracted. Count data was then normalised as Transcripts Per Kilobase Millions (TPMs) and used as such for quantification of expression (Supplementary Dataset 1). If a tissue had several replicates (max 3 replicates), TPM values were averaged, leading to a final number of 31 tissues compared. For each gene, Z-scores were then calculated to evaluate expression over-represented in a tissue, as Z = (x − μ) ∕ σ where for any sample: x is the observed TPM value in a given tissue, µ is the mean TPM value of all tissues, σ is the standard deviation of all tissues. We generated three Z-score tables: one encompassing all tissues & stages (Supplementary Dataset 2), another calculated only with TPM values in embryonic stages (Supplementary Dataset 3) and one calculated only with TPM values in adult tissues (Supplementary Dataset 4). For the analysis of gene expression dynamics during the developmental window spanning stages 22 to 31, we used Moran’s index as measure of temporal autocorrelation. Moran’s indexes and their statistical support were calculated for each gene with the R package spdep and its function moran.test following a data formatting step with ade4. To identify genes with high level and high specificity of expression in a given organ, we calculated a score integrating expression level and tissue specificity as follows: score = ln(TPM+1)*Z. For each tissue, we could then rank the genes of interest and focused on the 50 genes with the highest score.

### 2.8 Phylogenetic reconstructions

Phylogenetic analyses of gene families were conducted by retrieving protein sequences for all ohnologs (orthologs and paralogs) from a set of species including actinopterygians (*Erpetoichthys calabaricus*, *Lepisosteus oculatus*, *Danio rerio*, *Oryzias latipes*), amniotes (*Homo sapiens* or *Mus musculus*, *Gallus gallus* or *Anolis carolinensis*), an amphibian (*Xenopus tropicalis*), a lungfish (*Protopterus annectens*) and chondrichthyans (*Scyliorhinus canicula*, *Carcharodon carcharias*, *Amblyraja radiata*, *Callorhinchus milii*), as well as cyclostomes (*Petromyzon marinus*, *Eptatretus burgeri*) when available. Homologous sequences from the amphioxus *Branchiostoma floridae* were also included as an outgroup reference. This search was conducted using Genomicus, controlled by synteny data when available (Muffato et al. 2010; Louis et al. 2012; Louis et al. 2015) and TBLASTN (e-value<10^-10^) searches against Genbank or relevant predicted cDNA databases, controlled by reciprocal BLAST. The protein sequences were aligned using SeaView v4.6.4 (Gouy et al. 2010), MUSCLE v3.7 (Robert C Edgar 2004; R. C. Edgar 2004), and MAFFT v7 (Katoh et al. 2019). Gene tree inferences were performed using maximum likelihood with IQ-Tree (Minh et al. 2020) and ModelFinder to find the model of best fit (Kalyaanamoorthy et al. 2017), branch supports were given as percentages of 1000 SH-aLRT branch tests and 1000 UltrafastBootstrap tests (Minh et al. 2013).

### 2.9 3D reconstruction of synchrotron data

Iodine staining and synchrotron scanning of the head of a *S. canicula* hatchling (8cm total length) were performed as previously described (Leyhr et al. 2023). Briefly, the head was fixed in 4% paraformaldehyde, rinsed in 1x phosphate-buffered saline, and gradually dehydrated in 25%, 50%, 75%, and 100% ethanol. Staining with iodine was performed overnight in a 1% iodine solution in 100% ethanol according to the previously published protocol for I_2_E staining (Metscher 2009). After staining, the head was transferred to 96% ethanol before being scanned at 3µm isotropic voxel size using propagation phase-contrast synchrotron radiation microtomography (PPC-SRμCT) (Walsh et al. 2021) at beamline BM05 of the European Synchrotron Radiation Facility – Extremely Brilliant Source (ESRF-EBS) in France. Reconstructed jpeg2000 image stacks were imported into VGStudio MAX version 3.5.1 (Volume Graphics, Germany) for manual segmentation and rendering. Organ volumes and surface areas were calculated using the Porosity/Inclusion Analysis module. Only the left side of the head was manually segmented, and bilateral organ volumes are reported as double the left values, assuming bilateral symmetry.

### 2.10 Histology and *in situ* hybridisation

The head of embryos at stage 31 (late stage of organogenesis) and stage 34 (preceding hatching) and of a recently hatched juvenile (6 weeks post-hatching) were sampled from individuals previously euthanised, fixed in paraformaldehyde 4% in phosphate buffered saline 1x (PBS), and preserved in 100% ethanol (after graded dehydration in ethanol/PBS) for other studies. These samples were embedded in paraffin and cut at 10µm thickness. *In situ* hybridisations with RNA probes were performed as described previously (Lagadec et al. 2015) followed by nuclear counterstaining with Nuclear Fast Red (Sigma) and mounting in Eukitt. DNA matrice were synthesized by Twist Bioscience with probe sequences as given in Supplementary Table 5. For crystallin genes we designed two probes (against XM_038788722.1 and XM_038788824.1), both supposed to hybridise all of the recently duplicated crystallin gene copies as the nucleotide sequences differ only by about 5%. More distant copies (XM_038788959.1, XM_038790385.1, XM_038788706.1) differ from the probe sequence of about 15%. These two probes gave equal expression patterns. The probes for *v2rl1* and *v2rl4* were not considered to cross react (about 65% distance in the alignment between the probe and the potentially cross-reacting gene sequence). Similar dissimilarity was observed for *moxd2.1* and *moxd2.2* (60% distance one way, 75% the other way). Histological staining with Hematoxylin-Eosin-Saffron was made by an automat Leica autostainer X with standard protocol, at the RHEM platform in Montpellier on adjacent sections of the same individual (10µm-thick). All slides were scanned with a Nanozoomer S210 (Hamamatsu).

## 3. RESULTS

### 3.1 Genomic and transcriptomic resources

#### 3.1.1 Genomic resources

The chromosome-level genomic assembly was obtained from one male small-spotted catshark collected in the Mediterranean Sea (western Gulf of Lion, off Port-Vendres, France). The sequencing strategy relied on a combination of PacBio, 10x Genomics Chromium, Bionano and Hi-C, that resulted in a chromosome-level genome reconstruction including 31 chromosomes (covering 97,7% of the total genome) and 614 unassembled contigs, for a total of 4.22 Gb (Fig. 1A-B). These data are available in NCBI as sScyCan1.1 (RefSeq GCF_902713615.1), and summary statistics for the assembly are presented in Supplementary Table 6, with BlobToolKit (Challis et al. 2020) GC-coverage and cumulative sequence plots provided in Supplementary Figure 3. Twenty-one of these chromosomes are longer than 40 Mb (ranging between 71 and 313 Mb) and should therefore be considered macrochromosomes (Marlétaz et al. 2023), with the remaining ten microchromosomes (which includes the X chromosome) ranging in size between 13 Mb and 30 Mb (Supplementary Table 7).

**Figure 1.**
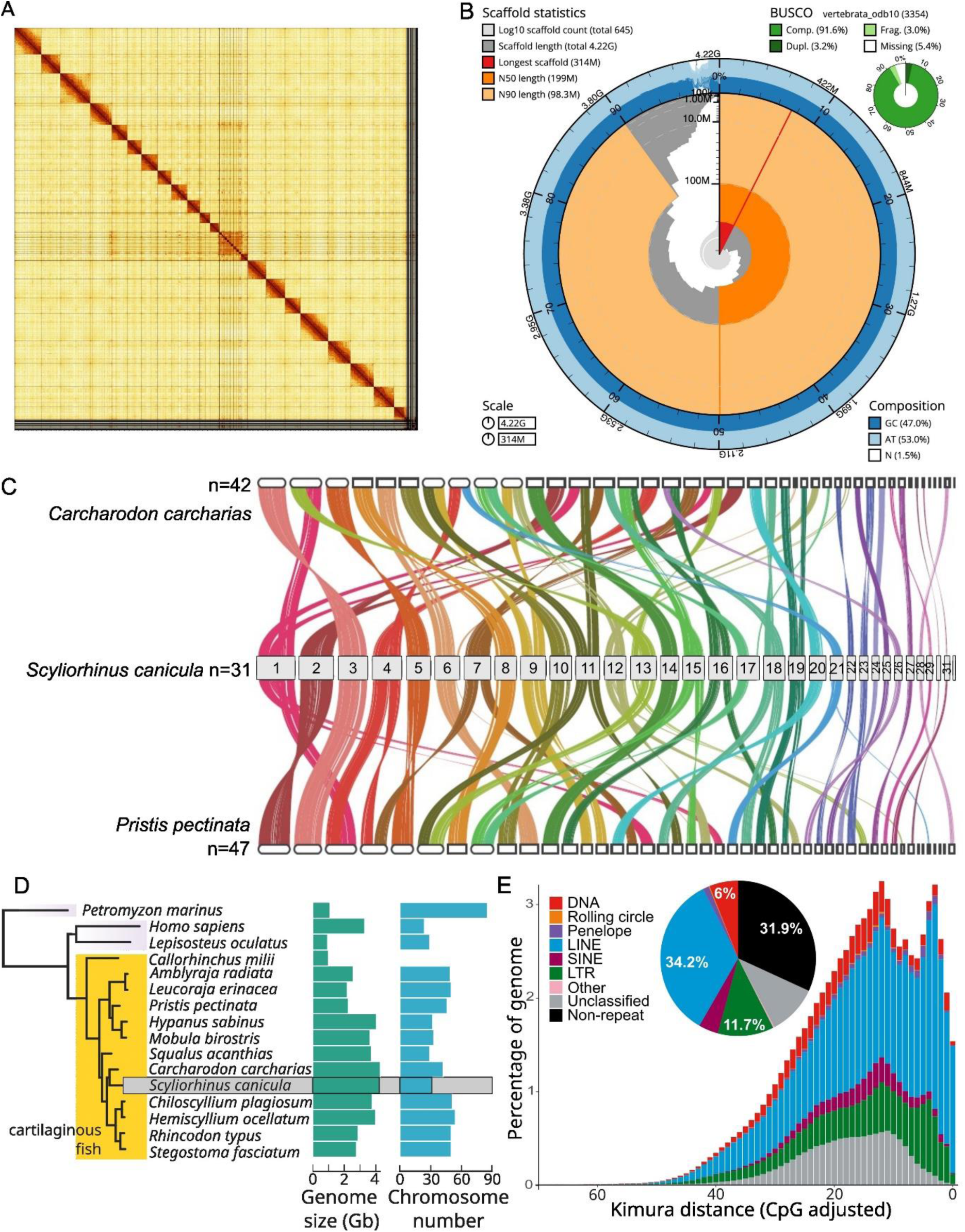
Genome assembly of *Scyliorhinus canicula*: (A) Hi-C contact map of the assembly, visualised using HiGlass. The absence of off-diagonal signals illustrates the correct assignment of sequence to chromosome-level scaffolds, bar the remaining unassigned contigs (bottom right). (B) The BlobToolKit Snailplot shows N50 metrics and BUSCO gene completeness. The main plot is divided into 1,000 size-ordered bins around the circumference with each bin representing 0.1% of the 4.2 Gb assembly. The distribution of scaffold lengths is shown in dark grey with the plot radius scaled to the longest scaffold present in the assembly (314 Mb, shown in red). Orange and pale-orange arcs show the N50 and N90 scaffold lengths (199 Mb and 98 Mb), respectively. The pale grey spiral shows the cumulative scaffold count on a log scale with white scale lines showing successive orders of magnitude. The blue and pale-blue area around the outside of the plot shows the distribution of GC, AT and N percentages in the same bins as the inner plot. A summary of complete, fragmented, duplicated and missing BUSCO genes in the vertebrata_odb10 set is shown in the top right. (C) Macrosynteny shared between three elasmobranch species. Synteny blocks based on alignment using all coding genes in three species, with 15,540 placed in shared synteny blocks between *Scyliorhinus canicula* and another shark, the great white shark *Carcharodon carcharias*, and 14,832 between *Scyliorhinus canicula* and a batoid, the small-tooth sawfish *Pristis pectinata*. Colours represent conserved synteny blocks and are ordered based on the focal species *Scyliorhinus canicula*. (D) Genome evolution across cartilaginous fishes: species phylogeny (left) consisting of 16 species including 3 outgroup species (see Material and methods). Bar plots show genome size distribution per species, and chromosome count per species (both estimated from genomes assemblies). (E) Pie-chart: proportion of the main TE classes in the assembled genome; graph: repeat landscape plot showing the proportion of repeats at different genetic distances (%) to their respective RepeatModeler consensus sequence, genetic distance is calculated under a Kimura 2 parameter model with correction for CpG site hypermutability: lower genetic distances suggest shorter time of divergence.

The genome of the small-spotted catshark has therefore over 60% macrochromosomes, as observed in most other shark genomes and in contrast to batoids species (Table 1). Macrosynteny is generally well conserved across elasmobranch fishes (Fig. 1C). However, the 31 chromosomes identified here represent a smaller karyotype than that reported for several other chondrichthyan species (Fig. 1D). Large synteny blocks carried by one chromosome in *S. canicula* are often distributed on 2 to 3 chromosomes in *C. carcharias* and *P. pectinata*, suggesting a number of recent chromosome fusions in the catshark or parallel fissions in the two other species (e.g. catshark chromosomes 1 or 4, Fig. 1C).

Analysis of GC content (Supplementary Figures 1 and 2) suggests a relatively stable GC percentage across chondrichthyan chromosomes, with a AT-bias in short (∼1Mb) unplaced scaffolds in *S. canicula*. Repeat sequences are highly abundant in the Mediterranean small-spotted catshark genome, with transposable element sequences accounting for more than two thirds of total genomic content (68.14%, Supplementary Table 8, Fig. 1E). LINEs are the most abundant transposable element sequence type by far (34.2%, Supplementary Table 8) and account for the vast majority of recently and currently active elements, as indicated by low sequence distances of annotated elements to their respective consensus sequences (Figure 1E). In addition, many LTR transposable elements (11.7%, Supplementary Table 8) and DNA transposable elements (6%, Supplementary Table 8) sequences are also present in the small-spotted catshark genome (Supplementary Table 8, Fig. 1E). However, both LTR TEs and DNA TEs appear to be decreasing in activity over more recent timescales (Fig. 1E).

A previously sequenced genome was obtained from an Atlantic individual (sex unknown) through Illumina sequencing of genomic DNA libraries (average fragment lengths ranging between 180bp and 8Kb) and resulted in a 4.31Gb draft assembly, consisting of 3,502,619 scaffolds (>200bp; maximum size 206 Kb; N50=6,665 bp). Both assemblies could be aligned over more than 90% of their length (Fig. 2B), with a 98.3% nucleotide identity. Unaligned sequences (about 9% and less than 1% of the Atlantic and Mediterranean genomes, respectively) essentially consist of repetitive sequences and no coding sequences were identified in these data.

**Figure 2.**
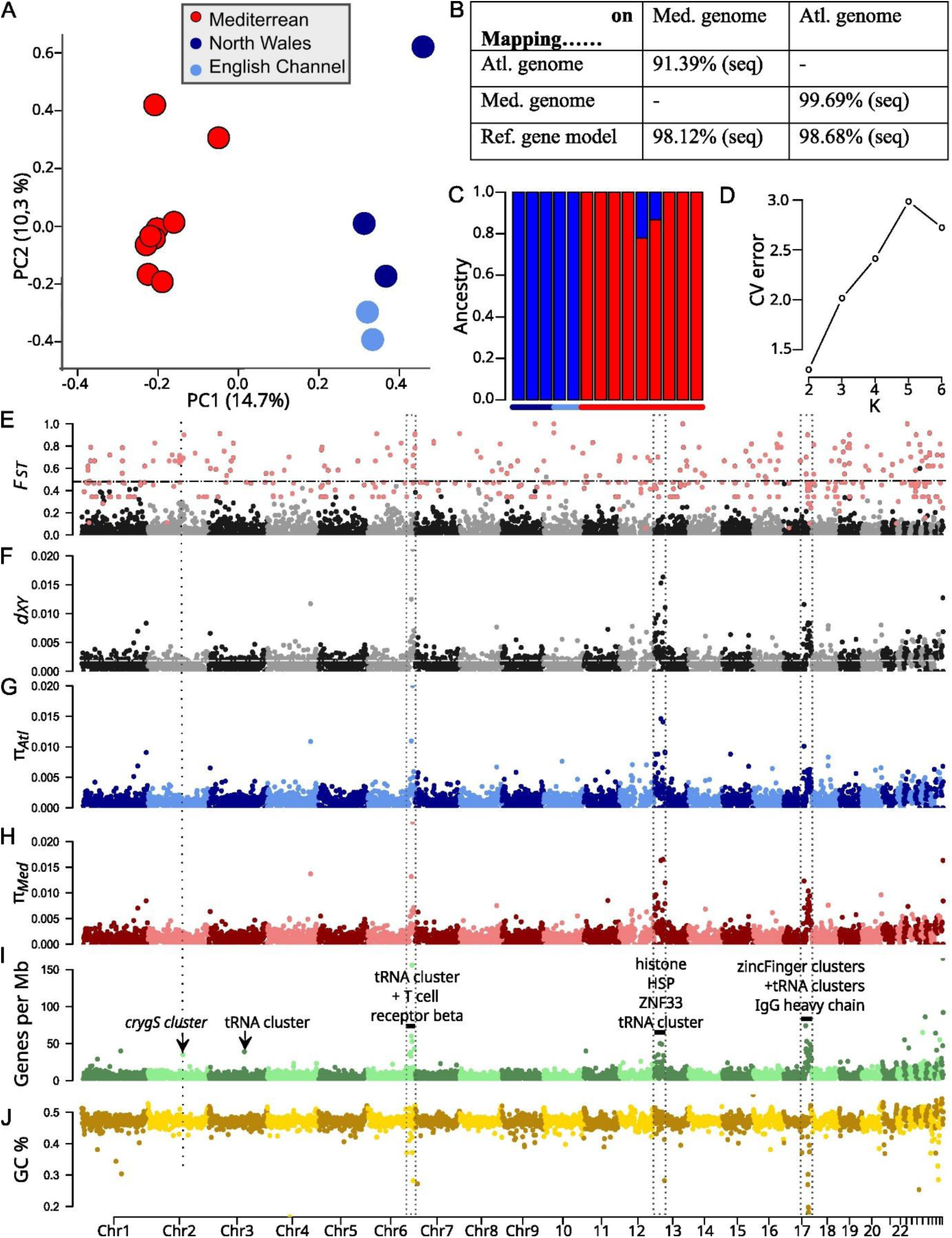
Atlantic and Mediterranean populations of the small-spotted catshark (A) Principal Component Analysis based on 12,005 SNPs separating Atlantic and Mediterranean samples. (B) Proportion of reciprocal coverage between the Atlantic and the Mediterranean small-spotted catshark genomes, and between genomes and the reference gene model used in this study (percentage of the sequences from the source (first column) that can be mapped on the target (first line). (C) ADMIXTURE analysis at *K* = 2. Bars show estimated ancestry proportion of each individual, and coloured lines at the top and bottom of the plot indicate the source population (same legend as in A). (D) Cross-validation error (CV error; *K* = 2 - 6; b) indicates that *K* = 2 was the best fit to the data. (E-J) Genome-wide patterns of differentiation (E: Weir and Cockerhams *FST*, based on 172,078 SNPs, dotted line indicates the mean value for all SNPs in the genome) absolute divergence (F: *dXY*, dotted line indicates the absolute value for the whole genome), nucleotide diversity in Atlantic (G: π*Atl.*) and Mediterranean (H: π*Med.*) populations, gene density (I: annotated loci per Mb) and GC content (J). Mean values for non-overlapping 1 Mb windows are shown as points in alternating colours for each chromosome. Red points in the *FST* plot show site-level *FST* values for variants which were significant outliers in the PCAdapt analysis.

Using RNAseq data acquired from individuals of known origin, we estimated the genomic diversity in a Mediterranean and two Atlantic (North Wales and English Channel) catshark populations (n=9 individuals, n=3 and n=2 respectively) using a SNP-based population differentiation analysis. The mean F_ST_ value across all variants is 0.044 (Supplementary Table 7) and the Mediterranean and Atlantic samples are better described with distinct ancestry (K=2) with minimal admixture (Fig. 2A, C-D) supporting differentiation between Mediterranean and Atlantic populations, in accordance with previous studies (Gubili et al. 2014; Kousteni et al. 2015; Ramírez-Amaro et al. 2018; Manuzzi et al. 2019; Melis et al. 2023). Significant outliers in the F_ST_ distribution were scattered across all chromosomes (Fig. 2E). Three genomic long-range regions (on chromosomes 6, 13 and 17) display high inter-population and intra-population nucleotide diversity values (boxed zones in Fig. 2E-J). Gene density also was high in these regions, and scanning through the annotated loci identified gene clusters with apparently a large number of recent duplicates (e.g. histone coding genes on chromosome 13, zinc-finger binding transcription factor coding genes on chromosomes 13 and 17). Our use of a RNAseq dataset obtained from gonads and early embryonic stages necessarily restrains the interpretation of the F_ST_ analysis. Several statistics of this genomic landscape show significant and strong correlation (Supplementary Figure 4) supporting the idea of high F_ST_ and nucleotide diversity values to come with regions of high gene density and GC content. As recombination rate is known to correlate with gene density and GC content, this genomic landscape may result from several forces, including balancing or divergent selection but also regional variation in recombination and mutation rates.

#### 3.1.2 Transcriptomic resources

Transcriptomic resources from a reference set of adult tissues and embryonic stages were generated by Illumina sequencing of cDNAs and used to annotate the Mediterranean chromosome-level genome, resulting in the prediction of 24,473 gene models. Alignment of these gene models against the Atlantic genome highlighted a 98.22% sequence identity (Fig. 2B), indicating substantial sequence variation in transcribed sequences between both origins. In order to improve the representativity of this gene model reference and extend the length of reference transcripts notably in 3’UTRs, we generated a second reference, referred to hereafter as *ncbi-utrs*, using a clustering approach and including NCBI collapsed isoforms plus transcriptomic data obtained by the CEL-Seq2 protocol (Hashimshony et al. 2016), which results in 3’UTR enriched sequence data. This *ncbi-utrs* database contains a total of 30,348 sequences (N50=5,181bp, mean length=2,980bp) and harbours a 95.7% BUSCO score with 3,208 complete BUSCO groups out of 3,354 searched (69 fragmented and 77 missing). In accordance with previous studies (Oulion et al. 2010), we identified 34 Hox genes in this dataset (Supplementary Figure 5, Supplementary Table 9). We found no evidence for the HoxC cluster known to be present in the elephant shark *Callorhinchus milii* (Ravi et al. 2009) and the zebra shark *Stegostoma tigrinum* (Yamaguchi, Uno, et al. 2023). Over 24,473 NCBI sequences, 11,945 (48.8%) were extended with a mean addition of about 1,845 bp per prolonged sequence. This *ncbi-utrs* reference was used for all subsequent analyses. Its mapping onto the Mediterranean and Atlantic genomes suggests excellent coverage for both assemblies, with only 1.88% and 1.32% of the ‘ncbi-utrs’ sequences respectively missing in each one.

We next took advantage of the transcriptomic reference to gain insight into (1) the dynamics of gene expression during organogenesis and (2) expression specificities of a broad range of adult tissues. Concerning organogenesis, we focused on a temporal window encompassing stages 22, 24, 26, 30 and 31, characterised by major morphogenetic changes in the nervous, sensory, cardiovascular, respiratory, excretory and musculoskeletal systems (Fig. 3A). This covers a long period of embryonic development: one month between stage 22 and stage 30, and another 20 days for stage 31 itself (Ballard et al., 1993). For each of these stages and each *ncbi-utrs* gene model, TPM (Transcripts Per kilobase Millions) and Z-score values were calculated to provide quantified indicators of relative expression levels and expression specificities of genes across this series of embryonic stages (Supplementary Dataset 3). Pearson correlation analysis for all genes across stages 22-31 shows that stages 30-31 are most clearly correlated and set apart from earlier stages, suggesting major developmental transitions between stage 26 and 30 (Fig. 3B). To further identify genes showing similarities in their expression dynamic between stages 22 and 31, we conducted a hierarchical clustering approach. This analysis was restricted to a set of 10,651 genes, selected for the coherence of their expression changes between successive stages (expression autocorrelation score across stages>0.1: Supplementary Dataset 5). In line with the Pearson correlation analysis, it led to the identification of two major gene clusters, referred to as clusters 1 and 2 (Fig. 3C; Supplementary Dataset 5). These two clusters respectively contain genes with higher expressions at stages 22-24 (cluster 1) and stages 30-31 (cluster 2) (Fig. 3C). Cluster 2 is further subdivided into four groups: clusters 2.1 and 2.2 contain genes whose expression increases at stage 26, with peaks at either stage 30 (cluster 2.1) or stage 31 (cluster 2.2), while both other clusters contain genes with a later increase of expression at stages 30 with highest expression at stage 30 (cluster 2.3) or stage 31 (cluster 2.4). Gene ontology (GO)-term analysis reveals very different functional annotations between each of these five clusters (Fig. 3D; Supplementary Dataset 6). Genes in cluster 1 (expression repressed at stages 30-31) are frequently associated with cell division (DNA replication, cell cycle, chromosome segregation and their regulation), pattern specification and some aspects of neuronal differentiation (neural fate specification, spinal cord motor neurons). As an example, several Hox genes and related long non-coding RNAs belonged to cluster 1 (Supplementary Figure 5), with peaks of expression either early (for ‘anterior’ Hox genes) or late (more ‘posterior’ Hox genes) in the stages 22-26 period.

**Figure 3:**
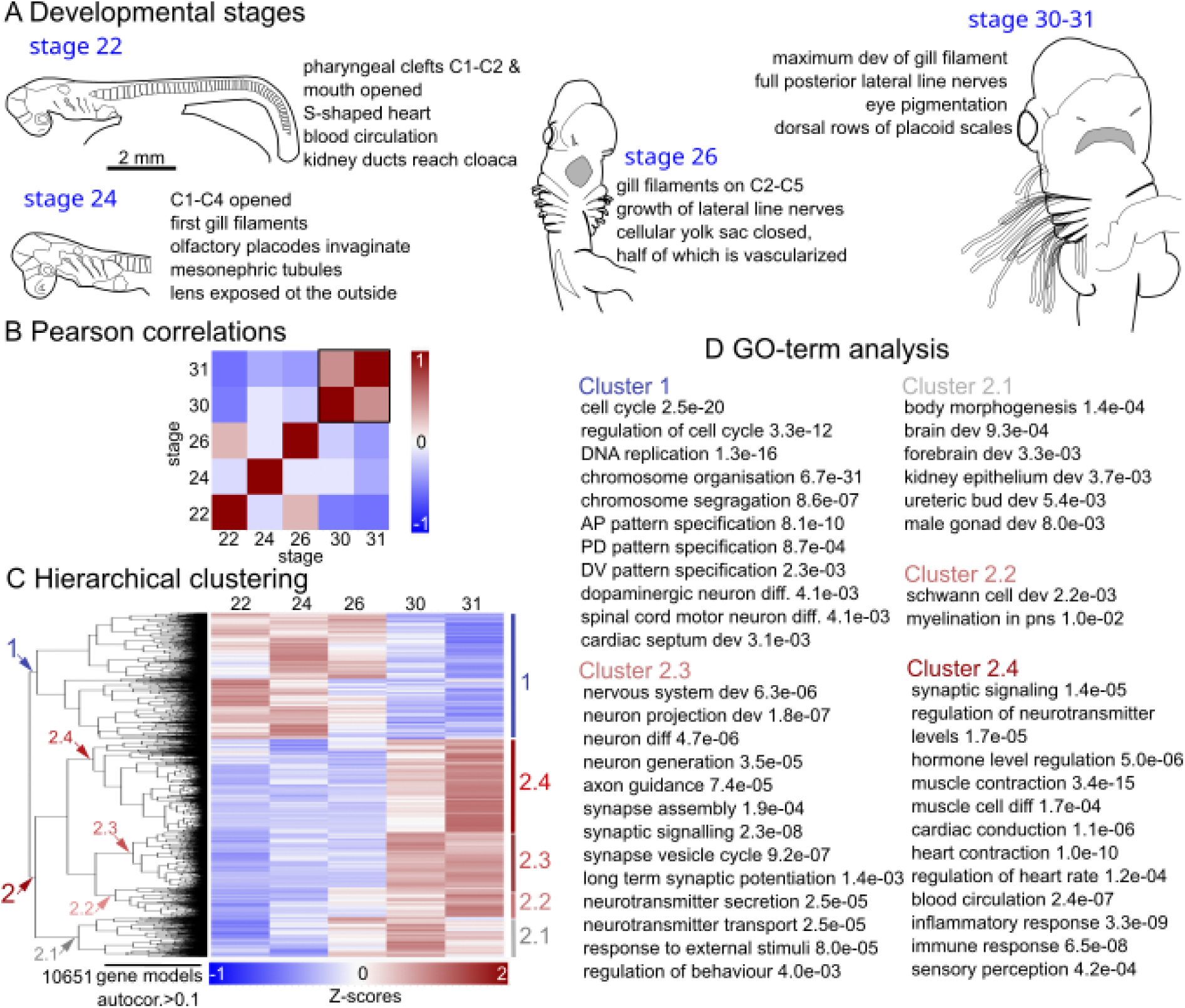
Transcriptomic comparison between stage 22, 24, 26, 30 and 31 embryos. (**A**) Broad morphological characteristics of the stages analysed (after Ballard et al., 1993). Lateral (stages 22 and 24) or ventral views (stages 26, 30-31) of a catshark embryo (stage 22) or its embryonic cephalic region (stages 24-31) are schematised for each stage. (**B**) Pairwise Pearson correlations between stages across all genes with sum of TPM>50. A black box shows the correlation observed between stages 30 and 31. (**C**) Gene groupings obtained by hierarchical clustering of Z-score transformed expression of the 10,651 genes with regionalised expression (autocorrelation>0.1). The two major clusters are respectively numbered 1 (in blue) and 2 (in red), the latter is further subdivided into four groups, referred to as 2.1, 2.2, 2.3 and 2.4. The corresponding nodes are shown by arrowheads in the dendrogram on the left, and the clusters are delineated by vertical bars on the right. (**D**) Selection of enriched GO-terms in clusters 1, 2.1, 2.2, 2.3 and 2.4, with p-values indicated. Detailed results for the GO-term analysis are shown in Supplementary Dataset 6. C1-C6 refer to pharyngeal clefts, from anterior to posterior. AP: anterior-posterior; dev: development; diff: differentiation; DV: dorsal-ventral; PD: proximal-distal; pns: peripheral nervous system.

GO-terms related to different aspects of organogenesis and myelination prevail in clusters 2.1 and 2.2 respectively (genes showing an increase of expression by stage 26). Genes in cluster 2.3 (expression established at stage 30) show a large number of highly significant enrichments in GO-terms related to the generation and differentiation of neurons, the formation of axonal and dendritic projections, the assembly and function of synapses, and the homeostasis of neurotransmitters, suggesting that the establishment of the basic architecture of the catshark neural circuitry is a major ongoing process at stage 30. GO-terms related to behavioural, cognitive, or sensory processes as well as to the formation of the circulatory system are also significantly enriched at this stage. A significant association with GO-terms related to synaptic and neurotransmitter signalling, as well as sensory perception is maintained for genes in cluster 2.4 (expression peaking at stage 31). Additional GO-terms enriched in this cluster include heart function, the differentiation of smooth and striated muscles, the differentiation of different lymphoid cell types, and hormonal and immune responses, suggesting that these processes become prevailing between stages 30 and 31 (Fig. 3D and Supplementary Dataset 6). Taken together, these data highlight relationships between specific gene expression dynamics and the initiation of biological or developmental processes in the developmental window analysed. The functional annotation thus provides a basis to search for underlying developmental regulators.

Concerning the analysis of gene specificities in adult tissues, we focused on organs of the nervous, digestive, excretory, cardiovascular and reproductive systems, as well as on a selection of sensory organs and skeletal structures (Fig. 4A). Z-scores and TPMs values were calculated for each gene and tissue included in this sampling (Supplementary Dataset 7). Pearson correlations for each pair of tissues and across selected genes (Z-score>1 and TPM>5 for at least one tissue) highlight positively correlated combinations of tissues, such as neural crest and paraxial mesoderm-derived skeletal components (Meckel’s cartilage, vertebrae and chondrocranium), central nervous system organs (brain, eye, spinal cord), striated muscles (hypaxial muscle, heart), mineralised tissues (dental lamina, skin denticles), tissues involved in the maintenance of osmotic homeostasis (rectal gland and kidney), and hematopoietic organs or cells (spleen, blood) (Supplementary Figure 6). Some unexpected correlations were retrieved between ovary, pancreas, stomach and liver. Because these organs all displayed a high proportion of genes with lower levels of expression, we interpret these correlations as an artefact due to shared low expression levels for a large set of genes. Another correlation was observed between gills, dental lamina, and skin denticles that may come from all three tissues being of epidermal origin, in addition to common processes of mineralised structure development shared by skin denticles and teeth (Debiais-Thibaud et al. 2011b; Debiais-Thibaud et al. 2019). For all tissues analysed, we retrieved genes possessing both high Z-scores and high TPM values, albeit with very different distributions along these two parameters depending on the tissue (Fig. 4B-E; Supplementary Figure 6 and Supplementary Dataset 7). Among those with the highest Z-scores, we identified many known markers of homologous organs or cell populations in osteichthyans (Fig. 4B-E; see also Supplementary Figure 6). For organs unique to chondrichthyans, such as the ampullae of Lorenzini, the rectal gland (a salt secreting osmoregulatory organ), or skin denticles (dermal scales), we establish lists of candidate gene signatures, which provide a starting point to examine their molecular functions (Supplementary Figure 6 and Supplementary Dataset 7). As expected, GO-term enrichment analysis for each one of these tissues led to the identification of terms relevant for known functions of these tissues (Fig. 4; Supplementary Dataset 8). In summary, despite some possible paralog misidentifications, unavoidable in automatic annotations, these transcriptomic data yield coherent molecular blueprints of selected organs, and provide a chondrichthyan reference to explore ancient organ molecular signatures across gnathostomes as well as chondrichthyan specificities.

**Figure 4:**
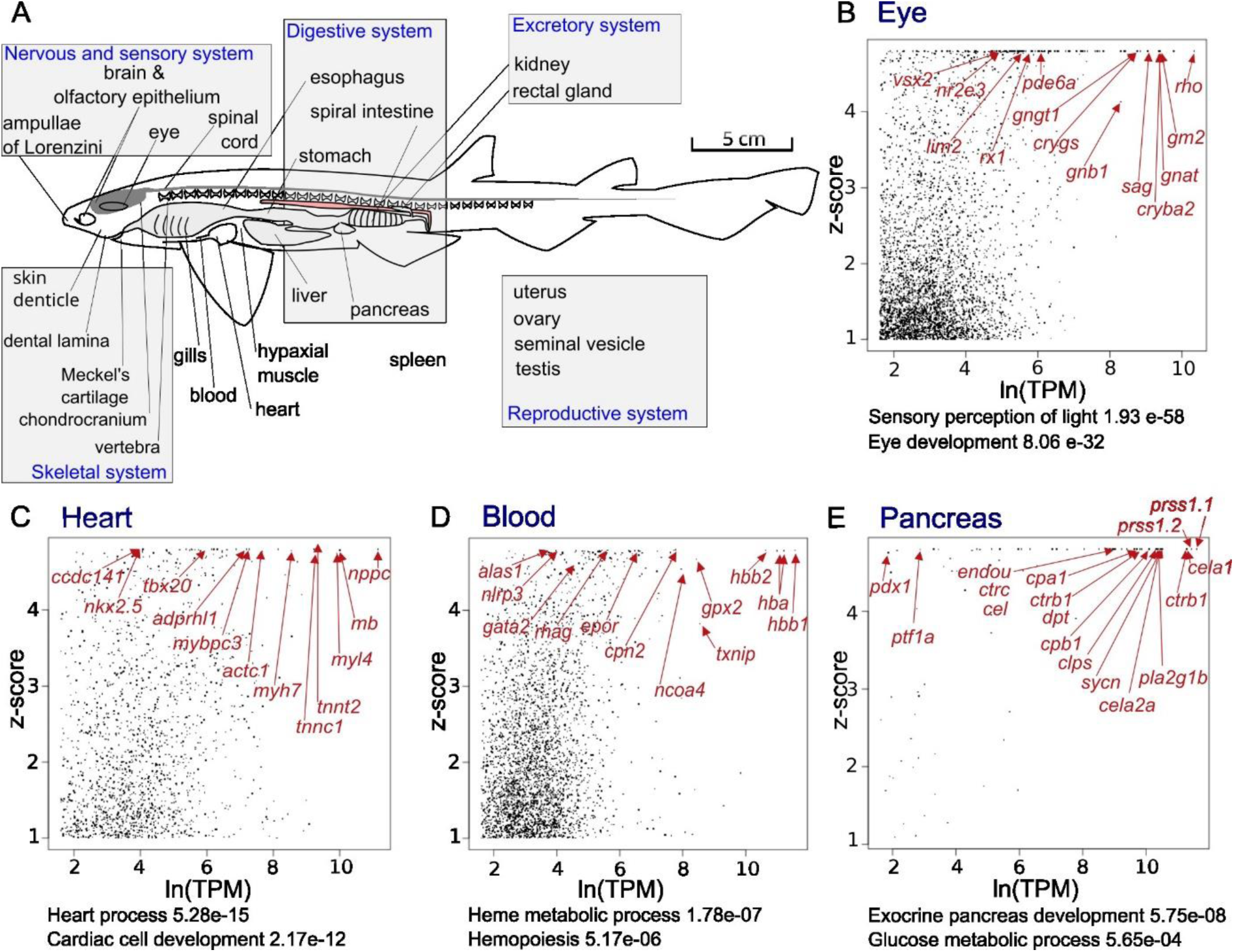
Transcriptomic analysis of small-spotted catshark adult tissues. (**A**) Scheme showing the adult tissues included in the transcriptomic analysis. (**B**), (**C**), (**D**), and (**E**) graphs showing Z-score and TPM values for each gene in the eye (**B**), heart (**C**), blood (**D**), and pancreas (**E**). Each dot represents one gene and the graph is restricted to genes with a Z-score>1 and TPM>5 in the tissue considered. GO-terms relevant to known functions of these tissues are indicated below the graphs. Red arrows point to a selection of genes, indicated in red, which exhibit high Z-scores in the catshark, consistent with selective expressions either in the eye (**B**), heart (**C**), blood (**D**), or pancreas (**E**) as documented in osteichthyans.

### 3.2 Characterisation of sensory organs and cells

#### 3.2.1 The anatomy of sensory organs in three dimensions

To provide a comprehensive morphological reference of sensory organs and their 3D organisation, we analysed the head of a late embryo (pre-hatching stage, 8 cm total length) by combining diffusible iodine-based contrast enhancement with propagation phase-contrast synchrotron radiation micro-computed tomography (DICE-PPC-SRµCT (Leyhr et al. 2023)). The reconstructed scan images were manually segmented to produce 3D renderings of the soft tissues (Fig. 5A-F). The volume of the whole head was measured to approximately 283 mm^3^. The eyes, nasal cavities, lateral line, and ampullae of Lorenzini (including their canals) altogether comprise 14% of the head total volume in this late embryonic stage (Fig. 5A-C), measuring 22.8 mm^3^, 8.6 mm^3^, 0.9 mm^3^, and 6.0 mm^3^, respectively. The sensory surface area of the ampullae of Lorenzini (approximated by excluding the canal surface, Fig. 5F) was 58.2 mm^2^, retinas were 46.6 mm^2^ (Fig. 5C) and olfactory epithelia (surface of the olfactory rosettes, Fig. 5E) were 146.3 mm^2^. The ampullae of Lorenzini therefore occupy a volume comparable to the olfactory organs, and have a sensory surface area comparable to that of the visual system. The 3D visualisation of the 219 left ampullae of Lorenzini and their canals (Fig. 5A-C) recovers the superficial ophthalmic (SO), buccal (BUC), and mandibular (MAN) clusters (following Norris, 1929; Rivera-Vicente et al., 2011). We can refine this clustering to 17 subclusters defined by distinct canal orientations and/or surface pore locations within the SO (10) and BUC (7) clusters (Fig. 5B-C). Through these subclusters, sensory ampullae within both the SO and BUC clusters are connected to surface pores on the anterior, dorsal, and lateral/ventrolateral surfaces of the head, with the greatest concentration of pores on the anteroventral surface of the snout (Fig. 5A-C, Supplementary Figure 7). The 3D identification of the canal, the ampullae and their sensory alveoli, with correspondence to histological observations is similar to data obtained in adult individuals despite the difference in size (i.e. 150-200 µm alveolar diameter at pre-hatching and juvenile stage (Fig. 6) *versus* 750-800 µm alveolar diameter in adults (Crooks and Waring 2013)). The number of alveoli was correlated with the size of the ampullae in our sample, suggesting the number of alveoli observed in a pre-hatching might not be representative of the adult condition.

**Figure 5:**
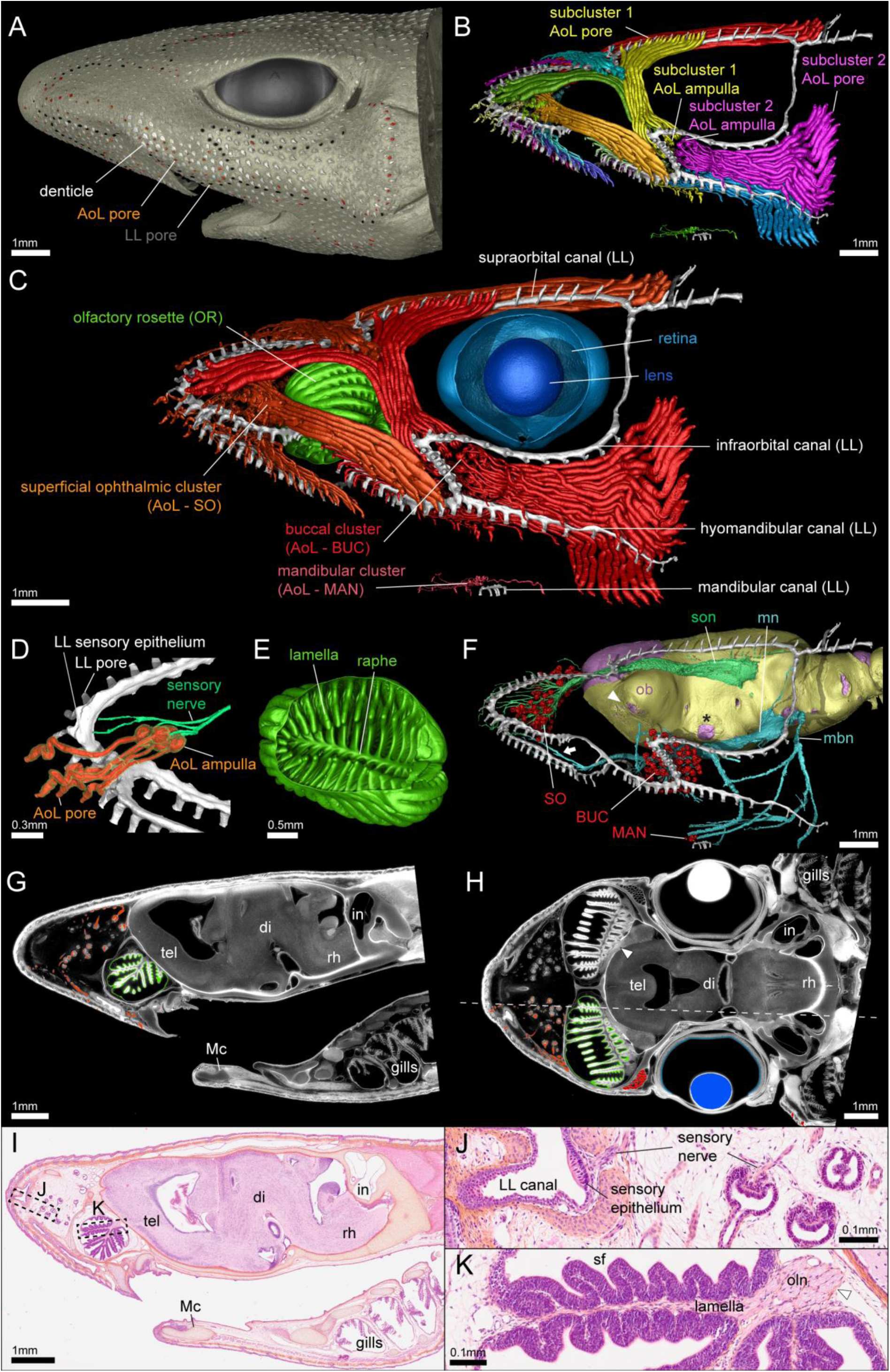
3D organisation of head sensory organs in a pre-hatchling *Scyliorhinus canicula*. (**A**) Lateral view of whole-head surface rendering, with dermal denticles in white and the lateral line (LL) skin surface pores in black. Surface pores for the ampullae of Lorenzini (AoL) canals are displayed in shades of red. (**B**) Rendering of subclusters of AoL as determined by canal orientations, shown in a range of different colours (lateral line in white). (**C**) Lateral 3D rendering of sensory organs. Shades of red = AoL with canals coloured for three clusters as defined by the location of the ampullae: orange = superficial ophthalmic (SO); red = buccal (BUC); old rose = mandibular (MAN); white = lateral line (LL); green = olfactory rosette (OR); blue = retina; dark blue = lens. (**D**) Medial rendering of the anterior snout (anterior to the left), isolating a single neural projection of the superficial ophthalmic nerve (light green) to both the lateral line (white) and several AoL with canals and sensory ampullae (orange = internal cast, transparent yellow = epithelium). (**E**) Anteroventral rending of the isolated left OR. (**F**) Lateral rendering displaying the interaction between sensory organs and the nervous system. Lilac = brain, yellow = internal fibrous surface of chondrocranium, light green = superficial ophthalmic nerve (son), light blue = maxillary (mn) and mandibular (mbn) nerves, white = lateral line, red = ampullae of the AoL system. White arrowhead indicates olfactory nerve foramen leading to the olfactory bulb (ob), black asterisk indicates optic nerve foramen. (**G, H**) Virtual parasagittal and horizontal histological sections with left sensory organs coloured as in (**C**). The dotted line in (**H**) indicates the position of the sagittal section displayed in (**G**). (**I**) Parasagittal histological section in plane corresponding to (**G**). tel = telencephalon, di = diencephalon, rh = rhombencephalon, in = inner ear, Mc = Meckel’s cartilage. (**J**) Histological detail in the lateral line and AoL. (**K**) Histological detail in the olfactory epithelium showing secondary folding (sf) of the sensory epithelium on each lamella and axon projections of the olfactory neurons (oln) emerging from the sensory epithelium.

**Figure 6:**
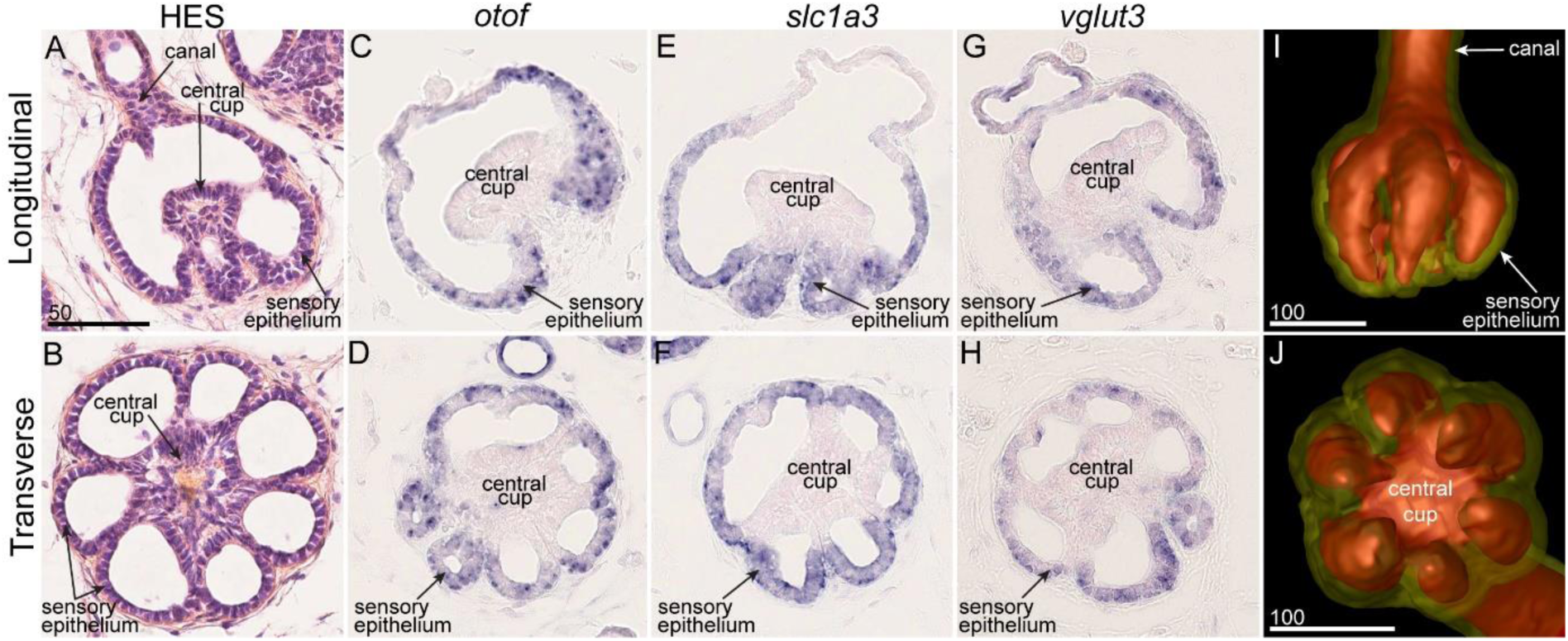
Histological (HES) staining of the sensory ampullae of Lorenzini in a juvenile catshark in their longitudinal (**A**) and transverse (**B**) sections. Gene expression patterns for *otof* (**C, D**), *slc1a3* (**E, F**) and *vglut3* (**G, H**) in similar orientations. (**I, J**) 3D renderings of a sensory ampulla (red = hydrogel-filled cavity, transparent yellow = epithelium) in similar orientations. Scales in microns.

The number of observed olfactory lamellae (∼33 per rosette, Fig. 5E) is consistent with the number in adult individuals (Ferrando et al. 2017) which is low relative to most other chondrichthyans (Ferrando et al. 2017; Dymek et al. 2021; Clark et al. 2022). Similar to adult olfactory organ, each olfactory rosette is partially separated by the central raphe into two compartments: one located dorsoanterior, the other ventroposterior. A large number of nerve bundles from individual lamellae run either lateral or medial to the olfactory rosette, cross the chondrocranium border through fibrous fenestrae, and merge into two short olfactory nerves (a medial and a lateral component), which reach the olfactory bulb medially and laterally in the anterior braincase (arrowhead, Fig. 5F, H, K, Supplementary Figure 8 and 9, Supplementary Dataset 9 and 10). The catshark olfactory bulb is bipartite, with two separate zones for olfactory nerve layer and glomerular layer, but a common granule cell layer (Supplementary Figure 9, Supplementary Dataset 9 and 10). The virtual sections confirm that medial nerve obtains input from medial lamellae, whereas the lateral nerve collects the output from the more laterally situated lamellae of the olfactory epithelium, each from both halves of the olfactory organ (dorsoanterior and ventroposterior, Supplementary Dataset 9 and 10).

Three major branches of the anterior lateral line nerve can be delineated: the superficial ophthalmic, maxillary (or buccal), and mandibular (following Boord 1977). The maxillary and mandibular nerves extend out of the chondrocranium in a single branch before dividing (light blue, Fig. 5F), while the superficial ophthalmic nerve emerges anterodorsally (light green, Fig. 5F). By visualising the anterior lateral line nerves and the sensory ampullae without their canals (Fig. 5F), it is apparent that the SO, BUC, and MAN clusters are innervated by the superficial ophthalmic, maxillary, and mandibular nerves respectively, with no visible overlap or exceptions (similar to observations in *Squalus acanthias* by Norris, 1929). One branch of the maxillary nerve extends anteriorly (arrow, Fig. 5F) but appears to interact exclusively with the anterior lateral line, not the electrosensory ampullae. Other anterior lateral line nerve bundles innervate both the lateral line and ampullae of Lorenzini sensory bulbs (Fig. 5D). Virtual histological sections (Fig. 5G, H) provide comparable morphological detail to traditional stained histological sections (Fig. 5I-K). Overall, the description of sensory organs in a pre-hatching stage resembles the adult situation despite the overall smaller size of organs, suggesting that sensory organs mostly display their final functional organisation prior to hatching.

#### 3.2.2 The sensory epithelium of the ampullae of Lorenzini

Several candidate genes for the ampullae of Lorenzini have been identified by differential expression in RNA-seq data in another shark, *Scyliorhinus retifer* (Bellono et al. 2017) and in a bony fish, the paddlefish *Polyodon spathula* (Modrell et al. 2017). Similar to Bellono et al. 2017, we found that the *parvalbumin alpha* (*pvalb*) gene is one of the most highly expressed transcripts with high specificity to the ampullae of Lorenzini (Supplementary Dataset 11). In addition, we uncovered specific expression of several ion channels previously associated to electroreception in elasmobranch and/or paddlefish electrosensory cells (Supplementary Dataset 11): the L-type voltage-gated calcium channel Cav1.3 (*cacna1d*), the calcium-gated potassium channel BK (*kcnma1*), and two voltage-gated potassium channel subunits (*kcnab3*; *kcn3*). Two other ion channels involved in signal transduction in these sensory cells are also found to be expressed in the ampullae of Lorenzini tissue, albeit without strong specificity (Z-score<1: *cacnb2* and *atp1b2*). We found specific expression of several transcription factors, including *atoh1*, *pou4f3* and *six2* (Modrell et al. 2017; Minařík et al. 2024), in the ampullae of Lorenzini (Supplementary Dataset 11). Finally, two genes known to code for presynaptic proteins of electrosensory cells in bony fish (Modrell et al. 2017), Otoferlin (*otof*) and Vglut3 (*slc17a8*), had very high specificity and levels of expression (Supplementary Dataset 11). Another highly expressed and specific transcript coding for a glutamate transporter, *slc1a3*, was also identified (Supplementary Dataset 11). We confirmed that expression of these last three genes was specifically located in cells of the sensory epithelium of the ampullae of Lorenzini, excluding the central cup (see **Fig. 6**).

#### 3.2.3 The olfactory epithelium: supporting cells versus olfactory receptors

Olfactory sensory neurons (OSNs) detect odorant molecules and pheromones by means of G protein-coupled receptors (GPCRs). Extensive research has uncovered the genetic variability underlying adaptive evolution of these olfactory receptor families in OSNs, from their origin and evolution in vertebrates to more recent events of gene family expansion or contraction (Saraiva and Korsching 2007; Grus and Zhang 2009; Bear et al. 2016; Dieris et al. 2021; Policarpo et al. 2022).

Examination of the 50 most abundantly and specifically expressed genes from the ‘brain & olfactory epithelium’ library (Supplementary Dataset 11) did not identify genes encoding GPCRs, but highlights two genes known to possess an olfactory role in bony vertebrates, *trpc2* and *s100z*, and two duplicates of *moxD2* that we named *moxd2.1* and *moxd2.2*. The vertebrate *Moxd2* gene encodes a monooxygenase named “Monooxygenase DBH Like 2”, of unknown but possible olfactory function (Hahn et al. 2007). Our genome screening and phylogenetic reconstruction recovered one *moxd2* gene in the great white shark genome and none in other available cartilaginous fish genomes, but the two duplicates in the small-spotted catshark have a time of divergence older than that of the last common ancestor of these two shark species (Supplementary Figure 10). Several duplicates were also found in *Petromyzon marinus*, *Protopterus annectens*, and *Erpetoichthys calabaricus*, but all duplication events appeared specific to each one of these three branches (Supplementary Figure 10). Both *moxd2* copies were found located in a single chromosomal region in *Protopterus annectens* and *Scyliorhinus canicula* (Chr8.part0 and Chr16, respectively), separated by other genes. The two genes found between the catshark *moxd2* duplicates (namely *emg1* (XM_038773333) and *peptidase inhibitor 16-like* (XM_038773822)) have orthologs on the Chr8.part0 of *Protopterus annectens* but the former is located more than 300Mb upstream and the latter 1,5Gb downstream of the *moxd2* genes. Little similarity between these loci in these two species and the phylogenetic relations of these sequences (Supplementary Figure 10) support a scenario of parallel tandem duplication events in these two lineages. Both catshark *moxd2* paralogs showed identical expression patterns in the most superficial cell layer of the olfactory epithelium, known to consist of supporting cells (Fig. 7A, B and Supplementary Figure 11 for moxd2.2 expression; Ferrando et al. 2010).

**Figure 7:**
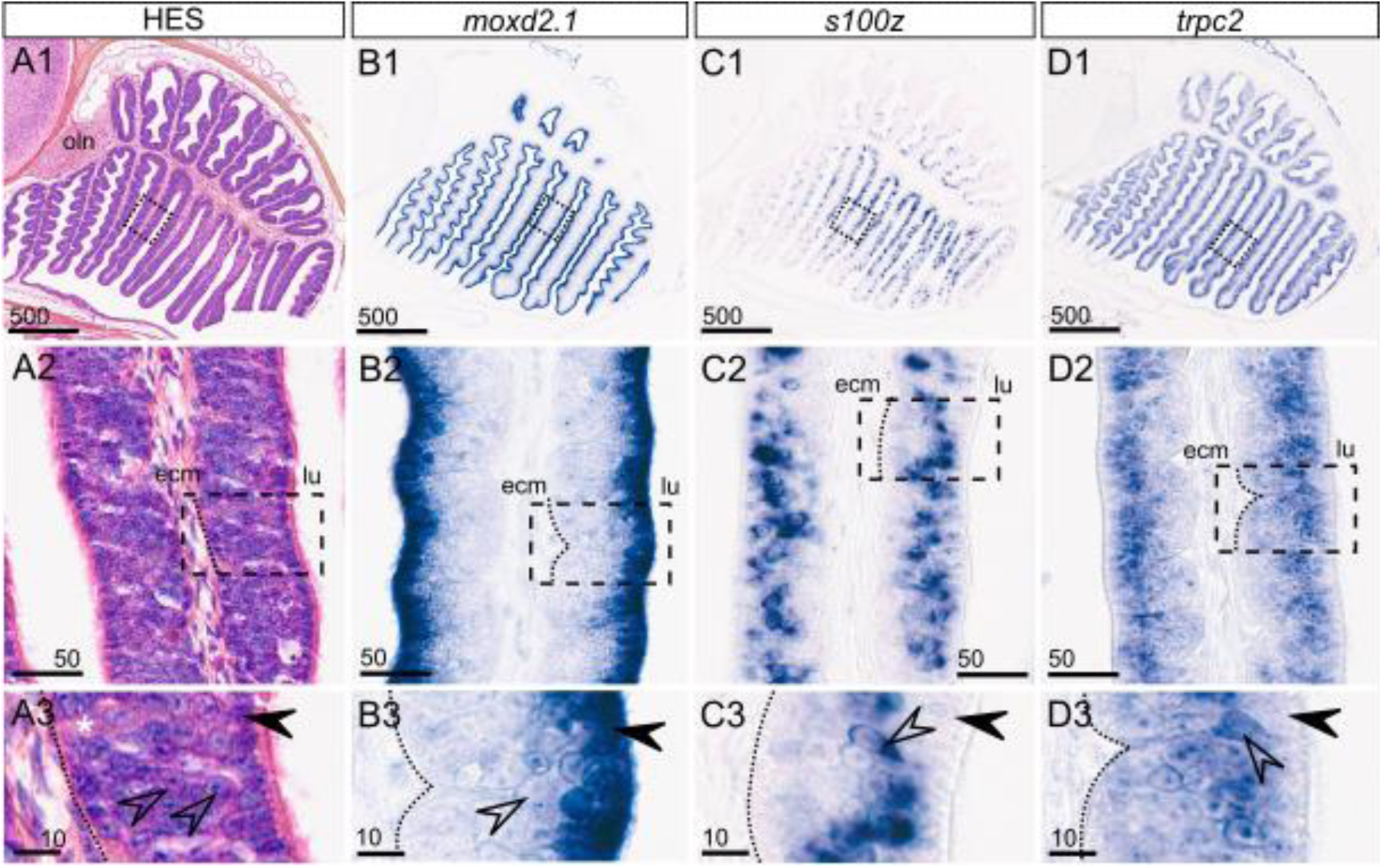
Histological (HES) staining of the olfactory epithelium in a juvenile catshark (**A**) with higher magnification (**A2, A3**) in transverse section. Gene expression patterns for *moxd2.1* (**B**), *s100z* (**C**) and *trpc2* (**D**) in parallel sections. ecm: extracellular matrix of the basal conjunctive tissue; lu: lumen of the olfactory rosette; oln: olfactory nerve; dotted line: basal lamina between the sensory epithelium and the underlying conjunctive tissue; asterisk: putative ionocyte (large clear cell); open arrowhead: putative sensory cell bodies; black arrowhead: putative supporting cells. Scales in microns.

Phylogenetic analysis identified a single *s100z* gene in each tested vertebrate species including the catshark (Supplementary Figure 12), and genomic data support conserved synteny between human and catshark for the genes neighbouring *s100z*. The *s100z* gene encodes a protein that belongs to the S100 family of calcium-binding proteins and is known to be expressed in a large population of microvillous OSNs in zebrafish (Oka et al. 2012). Pseudogenisation of *s100z* in mammals was found in species with a reduced or absent vomeronasal organ, which houses the microvillous OSNs (Hecker et al. 2019). Here we show that *s100z* expression is located in an intermediate layer of the catshark olfactory epithelium (Fig. 7C), known to host the sensory cell bodies (Ferrando et al. 2017; Syed et al. 2023). A similar expression pattern was observed for the *trpc2* gene (Fig. 7D), which is known to be expressed in microvillous OSNs in the catshark (Syed et al. 2023). The *s100z* cell population seems less contiguous compared to *trpc2*-expressing cells and may delineate a neuronal subpopulation.

### 3.3 Rise and fall of paralogs in sensory gene families in cartilaginous fish

We examined the evolutionary dynamics, gene-wise and expression-wise, of several candidate gene families known to play major roles in the respective sensory organs: the four main olfactory receptor families for the olfactory epithelium, opsins and crystallins for the eye, and the transient-receptor potential (TRP) ion channel family essential for diverse sensory functions. For each family, we established the catshark gene repertoire, showed its evolution in relation to other vertebrates using systematic phylogenetic analyses for unambiguous resolution of orthology relationships, and analysed tissue-specific expression patterns for selected family members.

#### 3.3.1 Small repertoire sizes of olfactory receptor gene families

The olfactory sense of vertebrates employs four major olfactory receptor families for detection of odours. They are the odorant receptors (OR), the trace amine-associated receptors (TAAR), the olfactory receptors class A-related (ORA) also called vomeronasal receptor type 1 (V1R), and the vomeronasal receptors type 2 (V2R, in bony fish also called olfactory receptor class C, OlfC). These receptor families were first identified in mammalian species before being found later in bony fish, and all four have also been detected in cartilaginous and jawless fishes (Nei et al. 2008; Sharma et al. 2019; Dieris et al. 2021; Kowatschew and Korsching 2021; Syed et al. 2023). The ORs are the largest olfactory receptor family of bony vertebrates (e.g. 1948 genes in the elephant *Loxodonta africana* (Niimura 2014)), but the catshark has only 2 canonical OR and 6 OR-like genes (Syed et al. 2023). The ORA family shows large gene expansions in tetrapods (here named V1Rs, Young et al. 2010), the TAAR family is massively expanded in teleost fish (Korsching 2020), similar to the TAAR-like (TARL) family in lampreys (Dieris et al. 2021), but these families consist of only 3, 3, and 2 genes respectively in the catshark (Syed et al. 2023). The restricted diversity of orthologs in these gene families in cartilaginous fish was shown to be linked to the loss of function of these receptors in the adult olfactory epithelium (Syed et al. 2023). Here we examined the potential alternate sites of expression of these receptors.

No specificity of expression towards the ‘brain & olfactory epithelium’ tissue was detected for either of the two canonical OR genes, *or1* and *or2*, nor the six OR-like genes (*or3* to *or9,* excluding *or7* that is not a predicted gene in the current version of the genome, but see Syed et al. 2023), supporting the loss of olfactory function in the catshark. For *or3* we have confirmed the absence of expression in the olfactory epithelium by *in situ* hybridisation (**Fig. 8A**). However, six of the eight OR genes showed highly specific expression in other tissues in the transcriptomic data (Supplementary Figure 13). Among these, *or3* and *or5* had their highest levels of expression in the ampullae of Lorenzini. In line with this, *or3* expression was detected in the cells lining the canals but not in the sensory zones of the ampullae (**Fig. 8B, C**). Transcripts of *or2* are broadly identified across tissues but over-represented in the testis (Supplementary Figure 13). Both *or1* and *or4* are expressed with high specificity in the spleen, and *or9* in stomach (Supplementary Figure 13). The specific albeit low expression of *or9* in the stomach may be related to an expression restriction to a small subpopulation of cells (Supplementary Figure 13), conceivably involved in sensing metabolic status (Kitamura et al. 2014). The two remaining genes, *or6* and *or8*, show overall weak and rather unspecific expression, and may exert their function in other tissues or developmental stages than those examined here (Supplementary Figure 13).

**Figure 8:**
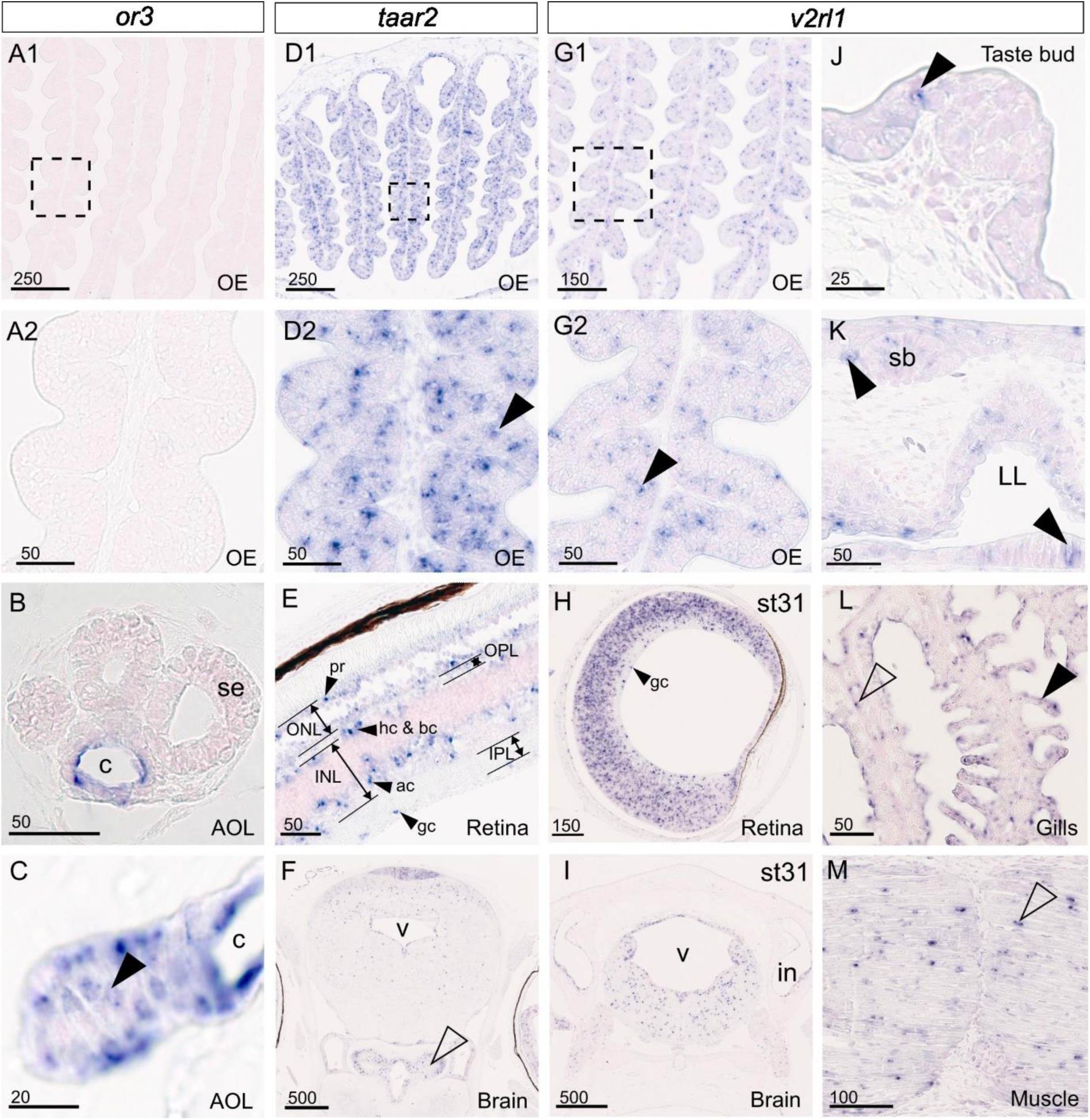
Gene expression patterns for *or3* (**A-C**), *taar2* (**D-F**) and *v2rl1* (**G-M**) in transverse sections of a juvenile catshark (except in **H, I**: stage 31 embryo) in the olfactory epithelium (OE; **A, D, G**) or non-olfactory sites: ampullae of Lorenzini (AoL; **B, C**), c: canal of an AoL, se: sensory epithelium of an AoL; differentiated (**E**) and undifferentiated retina (**H**), ac: amacrine cells; gc: ganglion cells, hc & bc: horizontal and bipolar cells; INL: inner nuclear layer, IPL: inner plexiform layer, ONL: outer nuclear layer, OPL: outer plexiform layer, pr: photoreceptor cell layer; posterior brain (**F, I**), arrowhead: posterior hypothalamus, in: inner ear, v: ventricle of the mesencephalon; taste bud (**J**); epiderm and lateral line (LL; **K**), sb: scale bud; gills (**L**) and muscle tissue (**M**). Black arrowheads point to scattered epithelial cells positive for gene expression; open arrowheads point to endothelial (**L**) or muscle (**M**) cells. Scales in microns.

Similarly, none of the genes of the ORA, TAAR or TARL show a high expression specificity for the ‘brain & olfactory epithelium’ in our transcriptomic data (Supplementary Figure 13), but some exhibit strikingly specific expression in other tissues, e.g. *taar2* in the eye and *tarl4* in the spiral intestine. *In situ* hybridisation with a *taar2* probe showed dense labelling in several brain regions and layered expression in the eye at both borders of the inner and the outer nuclear layer (layers that include photoreceptors, horizontal and bipolar cells, amacrine cells and ganglion cells, **Fig. 8E, F**). We also detected sparse labelled cells in the juvenile olfactory epithelium, suggesting a conserved function in olfaction, restricted to the juvenile stage of development (**Fig. 8D**) as *taar2* expression was not detected in adult olfactory tissues (Syed et al. 2023).

V2R is the main olfactory receptor family in the catshark (Sharma et al. 2019; Syed et al. 2023), and here we examined the 32 canonical *v2r* genes annotated in the gene model as well as three *v2r-like* genes. Twenty-one *v2r* genes show expression specificity in the ‘brain & olfactory epithelium’ sample (Supplementary Figure 14), consistent with their known expression in adult OSNs (Syed et al. 2023). Six genes show enriched expression in ‘brain & olfactory epithelium’ but their maximal Z-score is in another tissue (*v2r2*, *v2r5*, *v2r9*, *v2r11*, *v2r29*, *v2r33*) and four genes are not enriched in the ‘brain & olfactory epithelium’ sample (*v2r4*, *v2r10*, *v2r12*, *v2r28*). These data suggest that the large majority of *v2r* genes indeed have expression restricted to the olfactory sensory neurons, while others are additionally or exclusively expressed in other organs suggesting additional functions. For instance, a group of three closely related genes, *v2r9*, *v2r10,* and *v2r11* (Syed et al. 2023), also have enriched expression in the rectal gland (Supplementary Figure 14). This suggests that the *v2r9*, *v2r10*, *v2r11* clade may have undergone neofunctionalisation in the rectal gland, possibly in ionic homeostasis (Burger and Hess 1960). Along the same line, three phylogenetically distant genes (Syed et al. 2023), *v2r2*, *v2r5* and *v2r12* are highly enriched in the testis, suggesting three independent neofunctionalisation events. Two closely related genes, *v2r28* and *v2r29*, have enriched expression in some embryonic stages (22 to 26) and ‘brain & olfactory epithelium’ tissue (Supplementary Figure 14), although both with low levels of expression. Stages 22 to 26 correspond to early organogenesis and indeed the olfactory placode, visible already at stage 20, matures into a pit during these stages (Ballard et al. 1993), suggesting these genes may have acquired a function in the early development of the sensory olfactory placode. Several pairs of paralogous genes show very different expression specificities, suggesting that expression regulation is highly dynamic in this family (e.g. *v2r9*/*v2r10*, and *v2r33*/*v2r35a*). As a consequence, despite the v2r gene family showing the typical expansion seen for olfactory receptor gene families, their function is not restricted to the olfactory system and may involve such different sites as testis and rectal gland.

Finally, in the v2r-like gene family, only *v2rl4* has higher level of expression in the ‘brain & olfactory epithelium’ tissue, but also displays enriched expression in gills (Supplementary Figure 13). The other two genes, *v2rl1* and *v2rl3*, have their higher expression level first in gills, with *v2rl1* also more highly expressed in the oesophagus, skin denticles, and dental lamina (Supplementary Figure 13). Because the dental lamina, gills and anterior oesophagus are places where taste buds are found, we wished to better localise gene expression in relation to taste buds. *In situ* hybridisation for *v2rl1* and *v2rl4* showed virtually identical sites of expression, but no specific expression in taste buds, rather in cells scattered throughout the epithelium (**Fig. 8J** and Supplementary Figure 15). These genes also display expression in several brain regions (**Fig. 8I** and Supplementary Figure 15). Expression is also detected in the eye, intervertebral tissue, some chondrocytes, some muscle and gills, illustrating a wide array of expression sites during embryonic development and in the juvenile stage (**Fig. 8** and Supplementary Figure 15). Unexpectedly, *v2rl1* and *v2rl4* expression was also detected in sparse cells in the olfactory epithelium that may be OSNs (**Fig. 8G** and Supplementary Figure 15). Taken together, our study points to several independent heterotopic shifts of expression in the *v2r* and *v2r-like* families, with noteworthy diverse expression patterns of the *v2r-like* gene family, which suggests a broad range of functions in this clade of receptors.

#### 3.3.2 Contracted visual and non-visual opsin gene families

Visual opsins are light-activated receptors that are conserved in all animal groups. Together with their sister groups of non-visual opsins and non-canonical opsins, they belong to a wider group of GPCRs that are involved in signal transduction in animal cells (Koyanagi et al. 2021). Vertebrate opsins are classically categorised into several opsin subfamilies (Terakita 2005) and many lineage-specific duplications and losses were identified over the course of vertebrate diversification (Rennison et al. 2012).

In the small-spotted catshark genome, we identified two genes for visual opsins (Supplementary Figure 16): the one encoding Rhodopsin (*rho*) with extremely high and exclusive expression in the eye (Supplementary Figure 17), and the second visual opsin gene *rh2*, which has very low levels of expression in all sampled tissues (Supplementary Figure 17). *In situ* hybridisation of *rh2* showed expression in the poorly differentiated retina of a stage 31 embryo (**Fig. 9C**) but not in a juvenile (**Fig. 9B**), supporting a restricted transient expression in early retina progenitors. Rhodopsin therefore appears as the single visual opsin involved in light-sensing in the adult small-spotted catshark. The other three known gnathostome visual opsin genes (*sws1*, *sws2* and *lws/mws*, expressed in vertebrate cone cells) are absent in the catshark: *sws* genes are lost in all cartilaginous fishes while *lws/mws* loss is more recent as it is still found in some shark species (Yamaguchi et al., 2021). In addition to the isolated presence of the Rhodopsin visual pigment in the adult catshark retina, only three G-protein coding genes were recovered in our top-50 Eye list (Supplementary Dataset 11): *gnat* (G-protein, alpha transducing activity polypeptide 1), *gngt1* (G-protein, gamma transducing activity polypeptide 1) and *gnb1* (G-protein G(I)/G(S)/G(T) subunit beta-1) that are considered specific to signal transduction in rod cells only (Peng et al. 1992), congruent with the previous finding that the small-spotted catshark adult eye only has rod cells (Bozzanao et al. 2001) with the single rhodopsin pigment.

**Figure 9:**
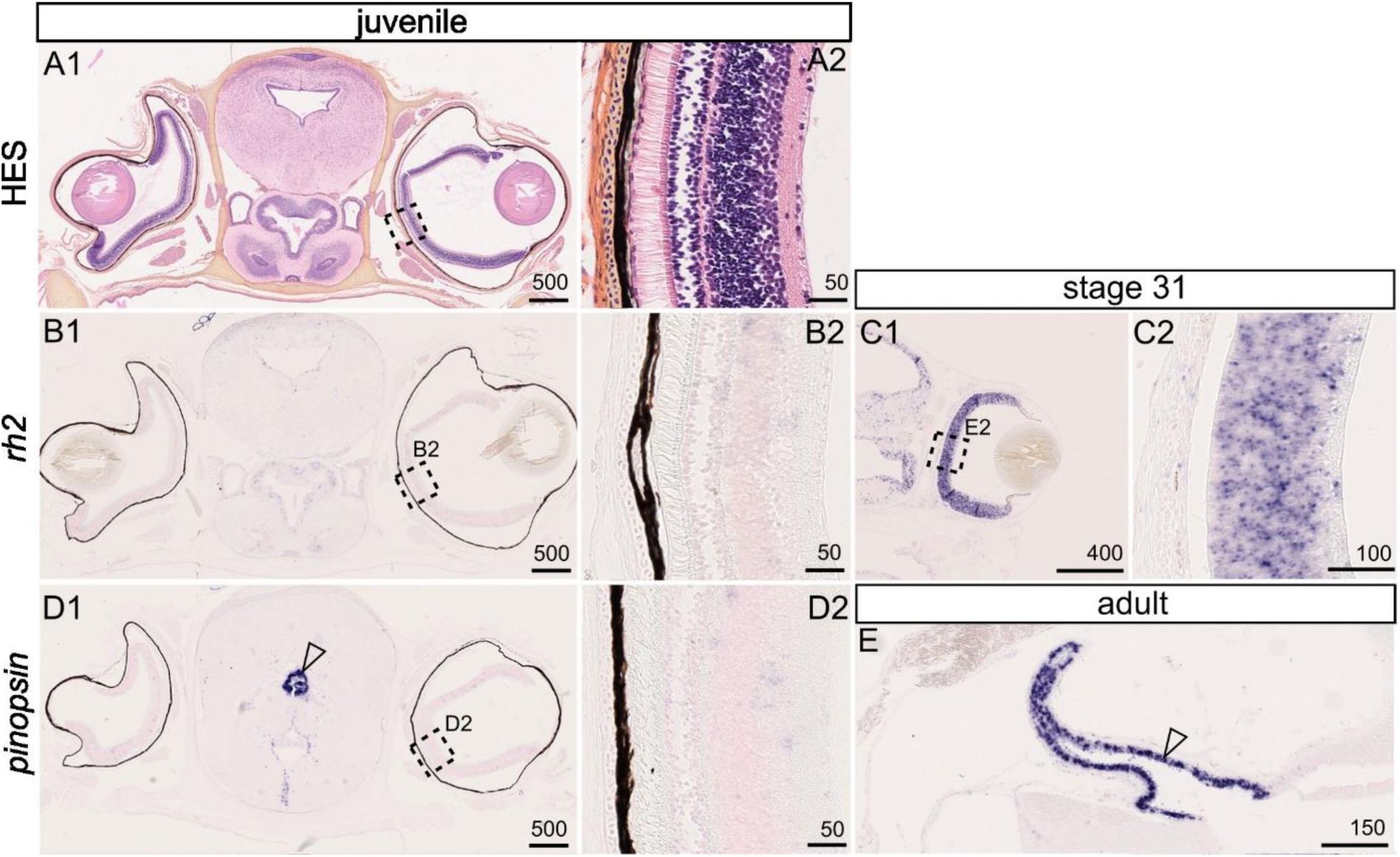
(**A)** Histological (HES) staining of a transverse section in the head of a juvenile catshark: general view (**A1**) with inset (**A2**) on the retina histology. Gene expression patterns for *rh2* (**B, C**) and *pinopsin* (**D, E**; arrowhead points to expression in the pineal stalk): in a juvenile (**B, D**); in the developing retina of a stage 31 embryo (**C**); in an adult catshark (**E**). Scales in microns.

Jawed vertebrates also have three canonical non-visual opsins: Pinopsin, VA-opsin and Parietopsin. The small-spotted catshark has all but the Parietopsin gene (Supplementary Figure 16), similar to other cartilaginous fishes (Yamaguchi et al. 2021). In the catshark, the gene coding for VA-opsin is only found expressed at very low levels but specifically in the eye, while the gene coding for Pinopsin is expressed quite ubiquitously at low levels with its zone of highest expression in the ampullae of Lorenzini (TPM=8.5; Z-score=2.88; Supplementary Figure 17). However, *in situ* hybridisations failed to detect expression of *pinopsin* either in the ampullae of Lorenzini or in the retina of a juvenile individual (**Fig. 9D2**). Instead, a highly specific expression was detected in the pineal organ of both a juvenile and an adult specimen (**Fig. 9D1, E**).

Non-canonical opsins include several subfamilies (Terakita 2005). In the Melanopsin subfamily, jawed vertebrates have up to two genes (*Opn4* and *Opn4x*). Only *opn4* is found in the small-spotted catshark (Supplementary Figure 18), similar to other elasmobranchs (Yamaguchi et al. 2021). The catshark *opn4* shows strong and specific expression in the eye (Supplementary Figure 17), supporting a conserved function in vision (Díaz et al. 2017). Within the Encephalopsin/TMT-opsin subfamily (Kato et al. 2016), the catshark genome includes the Encephalopsin (*opn3*) gene and a TMT1 gene (*tmt1*, Supplementary Figure 19), similar to other elasmobranchs (Yamaguchi et al. 2021). Encephalopsin expression is highest in the testis, then in embryonic stages and the ‘brain & olfactory epithelium’ samples (Supplementary Figure 17). Encephalopsin is named after its expression that was first described in brain tissues (Blackshaw and Snyder 1999), and extensive brain expression is known at embryonic stages (Davies et al. 2021). A function of opsins in sperm thermotaxis was demonstrated in the mouse, where Encephalopsin was the most highly expressed of all opsins in spermatozoa (Pérez-Cerezales et al. 2015). Both these expression patterns appear conserved in the catshark, suggesting sensing capabilities by opsins in catshark sperm cells. On the other hand, the catshark *tmt1* (Multiple-tissue opsin) is primarily expressed in the eye but at low levels, and its second highest site of expression is the spinal cord (Supplementary Figure 17). In the Neuropsin subfamily (Opn5/6), up to seven paralogs were identified in jawed vertebrates (Yamashita 2020) but only one was found intact in the catshark genome (Supplementary Figure 20), which is less than most other elasmobranchs (Yamaguchi et al. 2021). Expression of this gene, *opn5L1b*, is found in skin denticles, testis, eye, and spinal cord, although with low levels of expression (Supplementary Figure 17). From synteny comparisons with other elasmobranch *opn5m* genes, a pseudogenised sequence of *opn5m* was identified suggesting a recent loss of this opsin in the catshark lineage (Supplementary Figure 20).

Two other Opsin subfamilies are involved in the regulation of photoreception in vertebrates: the Peropsin (Rrh) subfamily involved in regulation of all-trans-retinol uptake from photoreceptors (Cook et al. 2017), and the Photoisomerase (RGR, or Opn6) subfamily involved in the modulation of retinaldehyde levels in retinal cells (Díaz et al. 2017). We found one gene for each subfamily: the catshark *rrh* (Supplementary Figure 21) is expressed with specificity in the eye while the catshark *rgr1* expression was highest in the eye and testis (Supplementary Figure 17). A remnant sequence of a pseudogenised *rgr2* is identifiable in the catshark locus where *rgr2* is found in other elasmobranchs (Supplementary Figure 22).

Overall, these data show a strong conservation of all aspects of photoreception in the small-spotted catshark despite a restricted gene repertoire (10 opsins out of the 26 paralogs identified in jawed vertebrates (see Yamaguchi et al., 2021), **Table 2**).

**Table 2:**
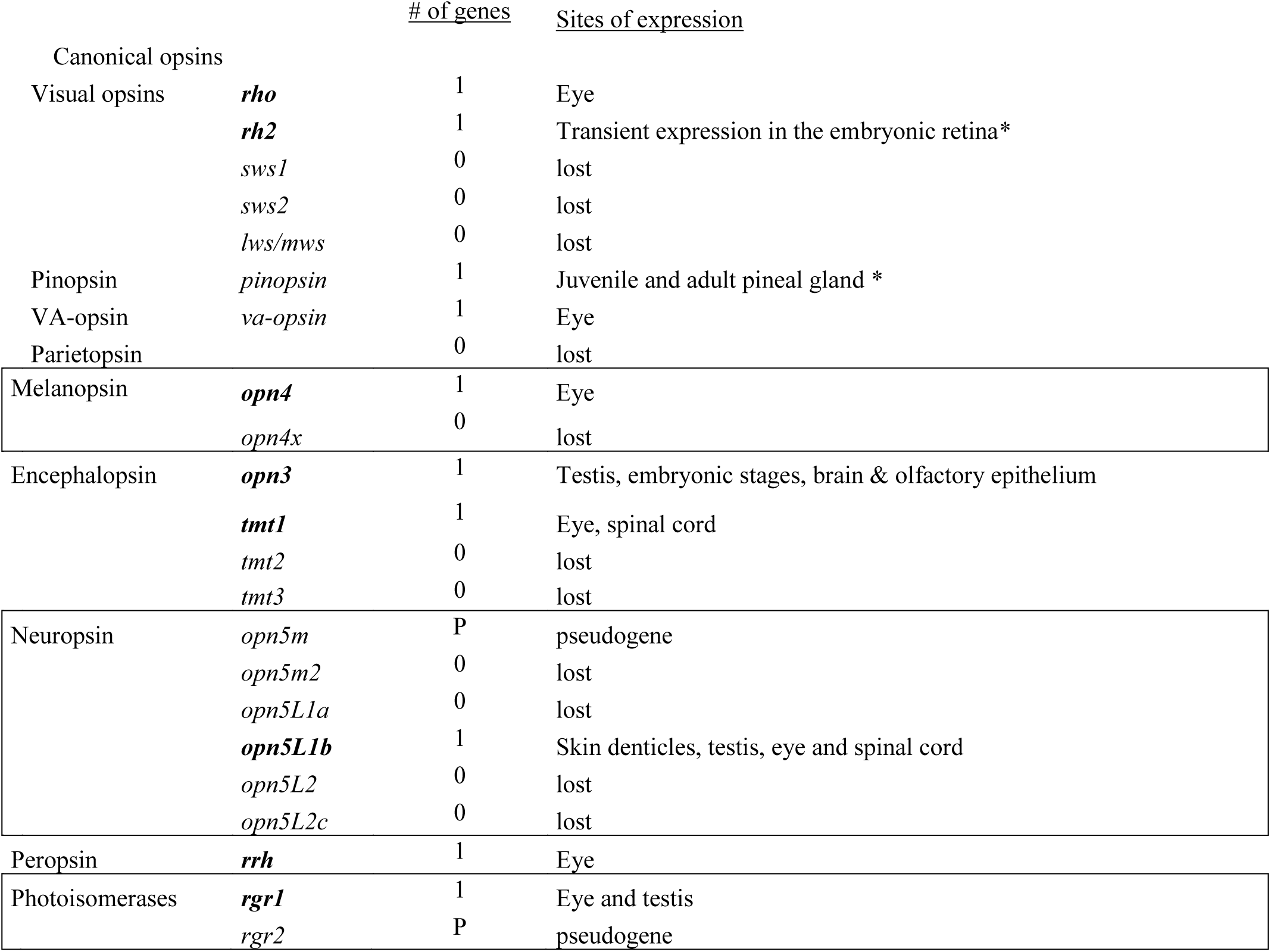
identified opsin gene complement (**bold**) in the small-spotted catshark with confirmed gene losses (grey). Sites of expression are defined as sites of over-represented expression in the RNA-seq data (Z-score>1), or confirmed sites of expression by *in situ* hybridisation (*).

#### 3.3.3 Amplified crystallin gene family

Vertebrate crystallins evolved with the vertebrate camera eye. Alpha-crystallins evolved from heat-shock proteins while beta- and gamma-crystallins evolved from calcium-binding proteins that gave rise to an ancestral chordate beta-/gamma-crystallin gene (Kappé et al. 2010; Slingsby et al. 2013; Cvekl et al. 2017). These proteins are highly expressed in lens fibre cells to ensure the transparency of, and a refraction gradient in, the lens of the vertebrate eye. Their genes were shown to have undergone rapid evolution in sequence and gene number in several vertebrate lineages in relation to ecological diversification (Slingsby et al. 2013; Malik et al. 2021) and are also linked to eye pathologies (Panda et al. 2016).

Out of the top 50 genes in our eye transcriptome (Supplementary Dataset 11), sixteen were annotated as crystallin coding genes. Of these, two were alpha-crystallins (*cryaa* and *cryab*) and five were beta-crystallins (*cryba1*, *cryba2*, *crybb1*, *crybb2* and *crybb3*), making this gene repertoire similar to other vertebrates (Slingsby et al. 2013) with probable high functional conservation (Ghahghaei et al. 2009). The remaining nine crystallin genes were categorised by the automatic annotation as gamma-crystallins. We searched the catshark genome for additional similar genes and identified a total of forty-one gamma-crystallin genes, showing a much higher diversity than described in mammals (Slingsby et al. 2013). To further understand the evolution of this clustered gene family, we generated a phylogenetic reconstruction of their relationships that identified well-supported vertebrate clades for one gamma-S and one gamma-N crystallin gene, both used as outgroups (Supplementary Figure 23 and Weadick & Chang, 2009). The catshark gamma-S gene (*crygs*) was located 2 Mb away from a dense cluster of all 40 other genes located along a 1Mb portion of chromosome 2 that also includes an *eya-4-like* gene (**Fig. 10B**). An additional non-coding RNA with sequence similarity to the surrounding gamma-crystallin genes was identified on the same cluster, suggesting recent pseudogenisation. Our phylogeny gives evidence for multiple series of duplication at different time scales. In particular, it shows a large chondrichthyan-specific clade and a large bony vertebrate-specific clade suggesting much of gene duplication has happened after the divergence of these two lineages. (Supplementary Figure 23). For both taxa, there are several copies in a given species that group together, showing recently generated duplicates in all lineages (the four human genes coding for gamma-crystallins A, B, C, and D group together, as a sister clade to thirty copies in the amphibian *Xenopus tropicalis* (Supplementary Figure 23)). For the cartilaginous fish clade of gamma-crystallins, we generated a nucleotide-sequence phylogenetic reconstruction for better resolution in the more recent nodes (Supplementary Figure 24): we uncovered a clade of 26 duplicates in the elephant shark *Callorhinchus milii*; a clade of 20 duplicates in the skate *Amblyraja radiata*; a clade of 16 duplicates in the great white shark and a clade of 30 duplicates in the catshark *Scyliorhinus canicula* (light blue boxes in **Fig. 10B**). These 30 copies constitute a monophyletic group and therefore originated from duplication events that occurred after the divergence from the great white shark lineage, less than 180 My ago (following TimeTree of Life, Kumar et al., 2022). Other duplicates, also located on the same 40 gene cluster, were closer to great white shark sequences so duplicated earlier in their common lineage (deep blue boxes in **Fig. 10B**). Species-related clades of gamma-crystallins in bony fishes suggest this “duplication hotspot” is a shared characteristic of jawed vertebrates. We compared the expression patterns of all copies in our transcriptomic data: two copies generated by the most ancient duplication events show a non-specific pattern of expression, but all others had expression specific to the adult eye and/or embryonic stages 30 and 31 (**Fig. 10C**). Notably, most copies generated by more ancient duplications had an adult eye-biased expression, while most of the more recent copies had expression biased towards the embryonic stages, sometimes in addition to strong expression also in the adult eye (**Fig. 10C**). We confirmed the location of expression of these recent copies in the embryonic and juvenile lens cells by *in situ* hybridisation (**Fig. 10D-F**). Tests for positive selection in the clade of these recently duplicated gamma-crystallin genes in the catshark were non-significant. Taken together, these data suggest that ancestral copies were expressed mostly in the adult eye, but that duplications, followed by partitioning of the ancestral expression pattern led to specialised sites of expression in the embryonic eye without positive selection for new protein functions.

**Figure 10.**
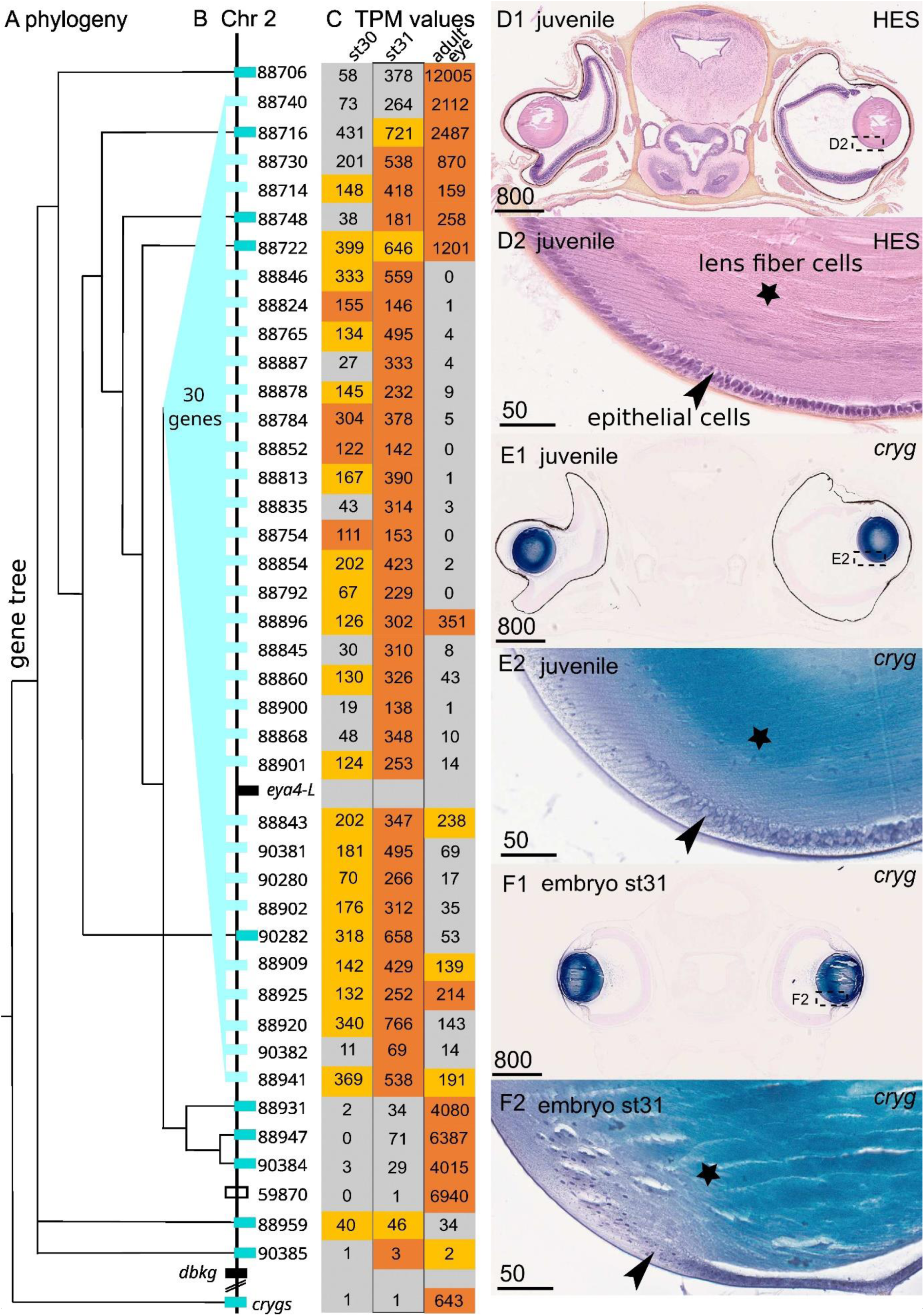
simplified representation of the phylogenetic relationships between the small-spotted catshark gamma-crystallins, with the single gamma-S crystallin gene (*crygs*) as the outgroup (**A**) mapped on the gene organisation along chromosome 2 (**B**); gene names appear as the last 5 digits of their genomic ID XM_0387xxxxx. TPM values in three selected samples (embryo stage 30, embryo stage 31 and adult eye) for each gene copy (**C**) with colour code indicated the associated Z-score (vermillon: Z-score>4; orange: 4>Z-score>1). Histological staining of a transverse section through the eye of a juvenile catshark (**D1** general view, **D2** close-up on lens cells) and *in situ* hybridisation with a gamma-crystallin (*cryg1*) RNA probe in the corresponding sample (**E**) and in a stage 31 embryo (**F**). The lens epithelial cells are labelled with an arrowhead, and the elongated lens fibre cells are marked with a star: both cell types are positive for *cryg* RNA *in situ* hybridisation. Scales in microns.

#### 3.3.4 A new vertebrate Transient Receptor Potential channels clade

The superfamily of Transient Receptor Potential (TRP) channels are classified in vertebrates as eight families belonging to two groups: group 1 with Trpa (Ankyrin family), Trpc (Canonical family), Trpm (Melastatin family), Trpn (nomp-C family), Trpv (Vanilloid family) and a more recently described Trpvl (Vanilloid-Like family); group 2 with Trpp (Polycystic family, that also includes Pkd1 and related paralogs, and Pkd2 and related paralogs) and Trpml (Mucolipin family) (Venkatachalam and Montell 2007; Peng et al. 2015; Himmel and Cox 2020). TRPs are structurally defined as ion channel subunits that form pores when organised in tetramers. All are permeable to cations but some are selective to calcium ions (Venkatachalam and Montell 2007). TRPs are considered polyvalent actors of sensation because of the wide diversity of physical and chemical stimuli that may activate their opening: in mammals they were shown to function in vision, taste, and chemesthetic sensation, olfaction, hearing, touch, nociception, and thermo- and osmo-sensation (reviewed in Venkatachalam & Montell, 2007). They function in the plasma membrane of excitable cells (e.g. depolarising sensory cells) but also in calcium signalling in non-excitable cells by modulating calcium movement through the plasma membrane or through intracellular membranes enclosing calcium reservoirs (reviewed in Gees et al., 2010).

Our extensive survey in the small-spotted catshark genome identified most of the expected TRP genes, but we also uncovered a previously unknown vertebrate clade of TRP. This TRP class was until now only identified in protostomes and non-bilaterian species and named Trpvl because in previous analyses it grouped as sister clade to the Trpv group (Peng et al. 2015; Himmel and Cox 2020). However, we recovered a sister relationship with Trpa with a strong support (**Fig. 11A**, Supplementary Figure 25): for this reason, we suggest to change the name of the class to Trpw. In the Trpw clade, one metazoan group is recovered (including sequences from protostomes and cnidarians) where an ortholog herein named Trpw1 is also identified in *Xenopus tropicalis*, *Protopterus annectens*, *Danio rerio* and *Lepisosteus oculatus*, in addition to their chondrichthyan orthologs. Three additional catshark copies group with one *Callorhinchus milii* and three *Petromyzon marinus* orthologs, although low support in the clade and lack of information from other gnathostome clades does not allow further identification of the duplication history within this clade. The catshark paralogs in this clade were herein named *trpw2.1*, *trpw2.2* and *trpw2.3* (**Fig. 11A**, Supplementary Figure 25). Their predicted proteins include the expected 6 transmembrane motifs that are involved in the ion pore structure, and all have 4 repeats of an ankyrin domain which make them structurally more similar to the Trpc family (Fig. 10B, Huffer et al. 2020). In the RNA-seq data, the newly identified *trpw1* gene exhibited its highest expression in the ‘brain & olfactory epithelium’ (Supplementary Figure 26). In situ hybridisation on sections of a juvenile showed scarce punctuated signal in the telencephalon (**Fig. 11D**) and a highly specific expression in the intermediate cell layer of the olfactory epithelium, known to mostly consist of the sensory neuron cell bodies (**Fig. 11E**).

**Figure 11:**
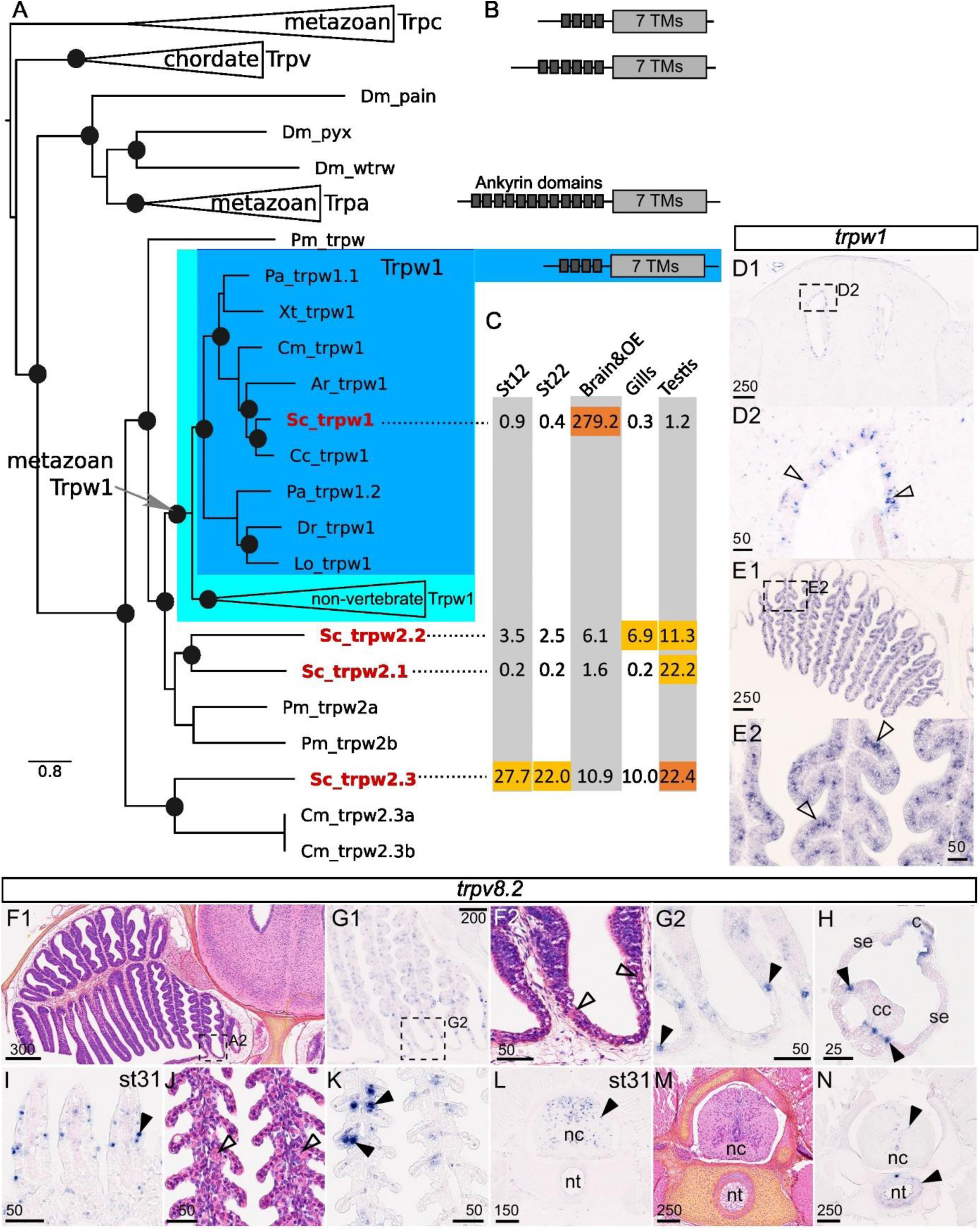
(**A**) phylogenetic relationships between the Trpc, Trpv and Trpa groups and the identification of a new metazoan Trpw clade: Trpc is taken as an outgroup and robust sister-relationship is recovered between the metazoan clades of Trpa and Trpw, excluding the chordate Trpv clade. Branch support is indicated by a black dot if the posterior probability value is >0.88. Species names: Ar: *Amblyraja radiata*; Cc: *Carcharodon carcharias*; Cm: *Callorhinchus milii*; Dm: *Drosophila melanogaster*; Dr: *Danio rerio*; Lo: *Lepisosteus oculatus*; Pa: *Protopterus annectens*; Pm: *Petromyzon marinus*; Sc: *Scyliorhinus canicula*; Xt: *Xenopus tropicalis*. (**B**) scheme of the ankyrin and trans-membrane (TM) protein domains. (**C**) TPM values for a selection of tissues (red underline for Z-score>4, light orange for 4>Z-score>1). (**D**) expression of *trpw1* in the telencephalon of a juvenile (arrowheads point to individual cells). (**E**) expression of *trpw1* in the sensory layer (open arrowheads) of the olfactory epithelium of a juvenile. **F, J, M**) Histological (HES) staining of transverse sections of a juvenile catshark through the olfactory epithelium (**F1**: general view with **F2** as a close up on the non-sensory epithelium), the gills (**J**) and anterior vertebra (**M**); putative ionocytes in the olfactory epithelium and gill epithelium are indicated with open arrowhead. Gene expression patterns for *trpv8.2* in the olfactory epithelium (**G1** and close up in **G2**), in an ampulla of Lorenzini (**H**), in the gills (**I**) and the vertebra (**L**) of a stage 31 embryo, and in the developing gills (**K**) and vertebra (**N**) of a juvenile catshark. Isolated cells with gene expression are indicated with a black arrowhead; c: ampulla canal (see Fig. 4); cc: central cup (see Fig. 4); nc: notochord, nt: neural tube; se: ampulla sensory epithelium (see Fig. 4). Scales in microns.

More generally in the group 1 TRPs, we uncovered all Trpc paralogs (Trpc1 to 7, Supplementary Figure 27) as well as the Trpn and Trpa paralogs (Supplementary Figure 28 and Supplementary Figure 29 respectively). We recovered most of the diversity of Trpm genes (gnathostome clades Trpm1 to 7) although Trpm8 sequences appear to be lost in all chondrichthyan genomes Supplementary Figure 30). The Trpv clade has an overall greater diversity of paralogs in chondrichthyans than in other gnathostomes (Morini et al. 2022). However, the Trpv1 and Trpv2 sister paralogs are restricted to sarcopterygian species and only one gene was found orthologous to these duplicates in chondrichthyans and in actinopterygians, herein named *trpv1/2* (Supplementary Figure 30, following Morini et al., 2022). The Trpv5/6 sequences have diversified in parallel through lineage-specific duplications in gnathostomes (Flores-Aldama et al. 2020; Morini et al. 2022), with a total of three chondrichthyan Trpv5/6 copies (herein named *trpv5/6.1*, *trpv5/6.2*, and *trpv5/6.3*; Supplementary Figure 31). In our transcriptomic data (Supplementary Figure 26), the three catshark Trpv5/6 paralogs have their highest expression (Z-score>1) in gills, but also ‘brain & olfactory epithelium’ and seminal vesicle (*trpv5/6.2*) or kidney (*trpv5/6.1*), supporting the hypothesised function in ion transport for excretion (Flores-Aldama et al. 2020). We also identified the two additional gnathostome paralogs that are the sister groups to the Trpv5/6 clade (Trpv7 and Trpv8 following Morini et al., 2022), one of them being duplicated in chondrichthyans (paralogs named *trpv8.1* and *trpv8.2* in the catshark; Supplementary Figure 31). All three genes have their highest expression in the gills, but both *trpv8.1* and *trpv8.2* also have high expression in the ampullae of Lorenzini and ‘brain & olfactory epithelium’ tissues, suggestive of sensory functions (Supplementary Figure 26). As these paralogs were only recently identified (Morini et al. 2022), we chose to characterise the expression pattern of one of them (*trpv8.2*): its expression was pervasive in the gill epithelium (**Fig. 11I, K**), congruent with expression by cells involved in ion transport. It is also expressed in the olfactory epithelium with expression in few cells of the sensory epithelium (thickened zone of the lamellae) but more cells in the non-sensory part of the epithelium (thinner epithelium between two lamellae, **Fig. 11F, G**), similar to the known distribution of ionocytes (Dymek et al. 2021). Expression in the ampullae of Lorenzini also suggested expression in a scarce population of cells that seem to be located in the zone intermediate between the central cup and the sensory epithelium (**Fig. 11H**). Further cellular characterisation will be necessary to identify putative ion transport in this zone. We also detected expression of this *trpv8.2* gene in the neural system (developing and juvenile spinal cord) and in a fibrous zone of the vertebral body (**Fig. 11L, N**).

In the TRP group 2, we recovered all three paralogs of the Trpml (Mucolipin) genes, and identified a recent duplication in the catshark lineage that generated two duplicates of *trpml3* (named *trpml3.1* and *trmpl3.2*; see Supplementary Figure 32). The expression of *trpml3.1*, but not of *trpml3.2*, appeared higher in the ampullae of Lorenzini and the ‘brain & olfactory epithelium’ tissues (Supplementary Figure 26). We also identified catshark orthologs for each three known gnathostome Trpp paralogs (*pkd2*, *pkd2L1* and *pkd2L2*, Supplementary Figure 33 with gene nomenclature following England et al. (2017)), with only *pkd2* showing over-expression in the ampullae of Lorenzini (Supplementary Figure 26). Despite poor current knowledge on the evolution of the Pkd1-related group (England et al. 2017), we uncovered a catshark ortholog for each of the gnathostome paralogs *pkd1*, *pkd1L1*, *pkdrej*, and *pkd1L2* genes (Supplementary Figure 34 and 35) but none showed biased expression toward sensory organs (Supplementary Figure 26). Two additional duplicates were found in several chondrichthyans, which we named *pkd1L4.1* and *pkd1L4.2* because they were found both outside of the known Pkd1L2 and Pkd1L3 clades (Supplementary Figure 35). *pkd1L4.2* had very low levels of expression, while *pkd1L4.1* showed higher expression in a range of tissues, including the eye, ampullae of Lorenzini, and spinal cord (Supplementary Figure 26).

Overall, the proper identification of paralogs in TRP gene families and their associated pattern of expression in RNA-seq data highlight several potential actors of sensory and ion-trafficking functions that could not have been identified from a simple analogy with what is known in classical model organisms.

## 4. DISCUSSION

### 4.1 High quality datasets of diverse nature fuel integrative analyses of morphological and molecular evolution

The catshark has attracted attention as an experimental chondrichthyan model for more than a century (Coolen et al. 2008), but the lack of a high-quality genome has been a major limitation to its development as a model. Here we report the availability of a chromosome-level genome assembly and transcriptomic data across a broad array of adult and embryonic tissues. We integrate these data with morphological descriptions and in situ analyses to gain new insights into the evolution of sensory systems in gnathostomes. This approach opens the field for a plethora of cellular and organ-level analyses in this important model organism.

#### Genome evolution of ‘big genome’ organisms

Although long neglected because of their large genomes, elasmobranchs have recently seen a number of chromosome-level genomes released. These resources now allow comparative genomic analyses within this group and with other jawed vertebrates, helping to identify the genetic bases of morphological or physiological diversifications in the early stages of vertebrate evolution. The catshark genome is one of the largest sequenced in chondrichthyans but has comparatively fewer chromosomes. This was suggested to reflect chromosomal organisation traits of ancestral gnathostomes (Marlétaz et al. 2023), but may also result from secondary fusions during the recent evolution of Carcharhiniforms, a hypothesis that could be further tested by synteny comparisons across a denser sampling of selachian genomes. The transposable element (TE) content in the catshark represents two thirds of the total genome, with a strong representation of LINE retrotransposons (34%). These results are very similar to those reported for other shark genomes, such as the great white shark (Marra et al. 2019), hammerhead shark and mako shark (Stanhope et al. 2023), and also for the little skate genome (Marlétaz et al. 2023) where TEs account for more than half of total genomic content, and where LINEs account for approximately one third of total genomic content. Expansion of LINEs accounts for larger genomes in other lineages of vertebrates (Tan et al. 2021), and this process appears to have shaped elasmobranch genomes for more than 270My (last common ancestor of elasmobranch fishes in Time Tree of Life (Kumar et al. 2022)). Further studies on the evolution of cell size in fossil elasmobranch and stem cartilaginous fishes may help identify the timing of genome expansion, following a strategy previously used in the study of salamander genomes (Laurin et al. 2016). Such large genomes have ecological consequences as they impact cell size, cell division time, and therefore reproductive and developmental features of the species. Sharks and rays have evolved diverse modes of reproduction (from egg-laying to several forms of live-bearing strategies (Katona et al. 2023)) that may be shaped to be compatible with these constraints. The availability of two sets of genomic sequences and RNA-seq data coming from two populations of the small-spotted catshark allowed to describe a population genomic landscape for the first time in this species. The identified outlier regions may contain the genetic bases of diverging morphotypes between Atlantic and Mediterranean populations (as identified in growth patterns or tooth shape (Berio et al. 2022)). Further studies using genomic re-sequencing data and a larger sample size could delineate these with more confidence.

#### Integrating scales of observation: genome, molecules, cells, organs

Genome-based technologies have opened novel perspectives to explore gene regulatory networks, cellular phenotype, organ development, and physiology model organisms that are neither genetically nor surgically amenable. Our integration of three very different methods (synchrotron imaging, histological observations and densely sampled RNA-seq data) exemplifies the potential of integrative approaches combining morphological observations at different resolutions and scales with gene-centred analyses for a comprehensive view of a complete system. The combination of high-resolution 3D reconstruction of organs at nearly cellular resolution using synchrotron imaging and 2D *in situ* hybridisation analyses on histological sections thus gives access to gene expression features with a cellular resolution in the complete organ context. This allowed for instance to pinpoint the cell heterogeneity in the olfactory epithelium, with only a small subset of cells expressing *trpv8.2*. Our bulk RNA-seq data themselves provide an unparalleled depth of molecular characterisation of organs, which opens perspectives on at least two levels of analysis. Firstly, our large diversity of twenty-five different adult tissues leads to unbiased, broad scale identification of organ-specific gene expression signatures. This expression information is an important complement to systematic comparative genomic approaches, aimed at identifying gene gains and losses relevant to morphological and physiological diversifications. The organ signatures thus provide putative gene identifiers of highly specialised cell types to be further characterised using advanced organ-focused approaches such as single-cell RNA-seq or spatial transcriptomics. Such data are crucial to explore the evolution of cell types within organs. Similarly, the dense transcriptomic sampling of a developmental window when major organogenesis processes take place leads to a fine-grained evaluation of the timing of underlying cell differentiation progression and concomitant gene expression dynamics. Secondly, the dense RNA-seq data combined with the availability of a chromosome level genome assembly allows comprehensive pictures of the structural and functional evolution of candidate multigene families of interest. On the one hand, the chromosome-level genome assembly allows unambiguous identifications of gene identities through additional phylogenetic analyses (as automated annotation occasionally leads to debatable ortholog identification), and helps identify gene losses or/and gene pseudogenisation (e.g. opsins) using synteny arguments. On the other hand, transcriptomic characterisations of a large sampling of tissues and embryonic stages provide expression information for each individual gene family member leading to hypotheses of neo- or sub-functionalisation.

### 4.2 Multi-pronged biological datasets in the catshark to understand the evolution of sensory systems in chondrichthyans and in early jawed vertebrates

#### Visual system: Opposing trends in two major gene families

Using these new resources, we identified and characterised a large locus in cartilaginous fish genomes made of a cluster of tens of gamma-crystallin genes. In all aquatic species, many of these genes grouped together as species-specific clades in our phylogenetic reconstruction, making them recent duplicates (in an amphibian, several actinopterygians (also see Lemopoulos and Montoya-Burgos 2022), and chondrichthyans). Considering these gene clusters as the product of many recent duplications that happened independently in all sampled lineages, we argue that our data point to an ancient duplication hotspot in the genomes of jawed vertebrates. Shared by cartilaginous and bony fishes, this locus of continuous duplication of gamma-crystallins may have been active already in early jawed vertebrates, most probably also associated with a high degeneration rate as an ancestral condition leading to the loss of any 1- to-1 orthologous gene between lineages. Consistent with this hypothesis, we identified in the catshark cluster a putative pseudogene: a non-coding RNA still showing extensive expression in the adult eye with a nucleotide sequence comparable to other gamma-crystallin genes. We did not detect positive selection within the most recent clade of gamma-crystallins in the catshark, suggesting that maintenance of large complements of similar genes is driven by selection on the quantity of the produced proteins, rather than by positive selection on sequence variants. However, such a gene cluster with high rates of duplication (and probably pseudogenisation) generates a large number of genes coding for lens proteins that may then be subject to subfunctionalisation and adapt quickly to ecological constraints of vision in deep or shallow waters (Weadick and Chang 2009). Such gene cluster may participate in population divergence, which should be further tested with adequate sequence data (e.g. eye RNAseq data to generate SNP comparison). A reduced necessity for large amounts of these proteins may have led to the loss of most gamma-crystallin genes in terrestrial vertebrates (Plotnikova et al. 2007; Cvekl and Eliscovich 2021).

An opposing trend was observed in the opsin gene family. With this high-quality genome of the small-spotted catshark, we confirmed the secondary loss of many opsin genes in the last common ancestor of elasmobranchs (*sws1*, *sws2*, *opn4x*, *tmt2*, *tmt3* and several clades of the *neuropsin* family), which was previously suggested on the basis of incomplete genomic data (Yamaguchi et al. 2021). In addition, we identified a previously undescribed catshark *rh2* gene with restricted expression in the undifferentiated embryonic retina. The general opsin gene loss identified for the elasmobranch lineage (where *rh2*, one of the *sws* genes, *opn4x*, *tmt*, *VA-opsin* and *pinopsin* were lost) might be compared to the losses associated with the ‘nocturnal bottleneck’ hypothesised in mammals during their early diversification (reviewed in Gerkema et al. 2013). These co-occurring losses of opsin genes (not observed in the elephant shark (Yamaguchi et al. 2021)) should now be more densely mapped onto the cartilaginous fish phylogeny to test if they could have resulted from a common event associated with the deep-sea life history for the elasmobranch common ancestor, or rather from gradual, progressive losses. The reduction of visual opsin genes in elasmobranchs seems to have been balanced by adaptative mutations in rhodopsin but also by structural adaptation of the rod cells (Bozzano et al. 2001; Yamaguchi, Koyanagi, et al. 2023). The catshark transcriptomic dataset further showed a strong specificity towards adult eye expression in some but not all of both visual and non-visual opsin genes, the latter being known to be expressed in the eyes of other gnathostomes (e.g. Upton et al. 2021; Salgado et al. 2022). Expression of opsins in other tissues raises questions on additional functions of these proteins, including at embryonic stages, as previously suggested (Liebert et al. 2022). A curious observation is the co-expression of *rgr1* and *encephalopsin* (Opn3) in the testis. Putative thermal sensation of sperm cells through expression of opsin proteins, as proposed in mammals, should also be investigated in elasmobranch fishes, in particular regarding the processes of sperm cell guidance to, and storage in, the spermatheca of females (Griffiths et al. 2012).

Finally, we identified distinct combinations of Trp gene expression in several sensory organs sampled in the small-spotted catshark, including in the eye (Table 3). These results suggest considerable conservation between cartilaginous fish and the lobe-finned lineage in visual organs, since a similar combination of Trps was reviewed for the mammalian retina, both in the photoreceptors and associated cells involved in transducing the signal (Pickard and Sollars 2011). Further comparisons with actinopterygians will be necessary to infer if this situation is conserved from the gnathostome last common ancestor or rather derived from convergent evolution.

**Table 3:** Summary of the TRP genes with higher expression in sensory organs (bold: Z-score>4; others: Z-score between 1 and 4). Asterisk: genes validated by *in situ* hybridisation; underlined: gene expression incongruent with sensory cell expression.

|  |  | Ampullae<br>of Lorenzini | Eye | ‘brain & olfactory<br>epithelium’ |
| --- | --- | --- | --- | --- |
| Group1 | Trpc |  | *trpc4, 5* | ***trpc2\****, *trpc3*, *trpc5* |
|  | Trpm | *trpm6* | ***trpm1, 3*** | ***trpm2\****, *trpm3* |
|  | Trpv | *trpv8.1*,<br><u>*trpv8.2\**</u> | *trpv9* | *trpv5/6.2*, *trpv3*,<br>*trpv8.1*, <u>*trpv8.2\**</u> |
|  | Trpw |  |  | ***trpw1\**** |
| Group2 | Trpml | *trpml3.1* |  | *trpml3.1* |
|  | Trpp | *pkd2* | *pkd1L4.1* |  |

#### Olfactory system: Neofunctionalisation in three minor and one major olfactory receptor families

We show a morphologically and molecularly well-developed olfactory epithelium already at late embryonic stages, and described it in 3D as very similar between the pre-hatching and adult stages despite the size difference (Schluessel et al. 2010). We found no anatomical evidence for a separate accessory olfactory system such as those observed for both lamprey and lungfish, in this pre-hatching stage (Ren et al. 2009; Nakamuta et al. 2012). If this is representative of the adult condition, then the catshark is similar to ray-finned fish, rather than to lungfish. This might suggest that a single olfactory epithelium is the ancestral state for vertebrates and that both accessory olfactory systems described in the lamprey and lobe-finned fishes (lungfish and tetrapods) have evolved independently. The anatomical arrangement of this olfactory epithelium and its projection through two olfactory nerves to the olfactory bulb is consistent with previous reports at earlier catshark stages (Quintana-Urzainqui et al. 2014), appears similar to a recent report for adult bamboo shark (Camilieri-Asch et al. 2020) and seems to be widespread in the chondrichthyan lineage (Yopak et al. 2015). Since all molecular markers investigated show similar expression in all lamellae, the division in two olfactory nerves does not appear to reflect a functional subdivision of the olfactory epithelium.

Our analysis of highly enriched expression yielded several genes with specific expression patterns in the olfactory epithelium. Trpc2 is a component of the signal transduction cascade in olfactory microvillous neurons of bony vertebrates (Liman et al. 1999; Stowers et al. 2002; Liberles 2014) and has often been used as marker gene for these neurons (e.g. see Sato et al. 2005; Hecker et al. 2019; Syed et al. 2023). The high expression levels of *trpc2* are consistent with its presence in (nearly) all olfactory receptor neurons in the catshark (our results and Syed et al. 2023); the other main population of olfactory sensory neurons, ciliated neurons, are absent in sharks (Ferrando and Gallus 2013). The expression of *trpc2* in the olfactory epithelium suggests the gnathostome-wide conservation of Trpc2 function in chemo-sensation. The catshark gene *trpw1*, a vertebrate Trpw gene clade first described in this work, shows a very similar expression in cells of the neuronal layer, albeit less frequent than *trpc2*-expressing cells, consistent with *trpw1* expression labelling a subpopulation of microvillous neurons. In addition, the calcium-binding protein S100z has been suggested as another marker of microvillous receptor neurons in bony vertebrates (Kraemer et al. 2008; Oka et al. 2012; Hecker et al. 2019), and our results suggest it to be shared by all gnathostomes. Moreover, *s100z*, together with *trpw1,* expression appears to label fewer cells than *trpc2* suggesting that different subpopulations of microvillous neurons may occur in the catshark.

Moxd2 is a monoxygenase related to dopamine beta hydroxylase that originated in vertebrates as a duplication of an older *moxd* gene. The function of Moxd proteins is unknown in any species, but its relation to DBH and the presence of *Moxd2* expression in mouse medial olfactory epithelium (array data in Su et al. 2004) have led other authors to suggest a function in neurotransmitter conversion or modification of incoming odorant molecules (Hahn et al. 2007). We note that high expression levels of *moxd2* (similar to *s100z* levels) are present also in a recent transcriptome of zebrafish olfactory epithelium (gene name moxd1l, Calfún et al. 2016). Our transcriptome data show that enrichment of *moxd2* in the olfactory epithelium appears to be a general gnathostome feature. Moreover, we identify for the first time the cell type expressing *moxd2* genes in the catshark. Both *moxd2* paralogs are abundantly expressed in the supporting (sustentacular) cells of the catshark olfactory epithelium (apical layer), but not in the neuronal nor basal cell layer. It is currently unknown whether the *moxd2* genes of bony vertebrate species share the same expression in supporting cells. However, supporting cells were shown to metabolise odorants by a two-step process involving monoxygenases in phase I (P450 cytochromes, Chen et al. 1992), and several transferases in phase II (olfactory UDP glucuronosyl transferase, Lazard et al. 1991; and sulfotransferase, Miyawaki et al. 1996). Moxd2 may then act in the supporting cells as a phase I enzyme to begin metabolisation and thereby removal of odorants from the catshark olfactory epithelium. However, *moxd2* genes were apparently lost in parallel in a variety of lineages (several chondrichthyans, our study; humans, Hahn et al. 2007; catarrhines and whales, Kim et al. 2014) suggesting functional redundancy with other monoxygenases. Conservation of two *moxd2* paralogs with similar expression pattern in the small-spotted catshark should be better understood by testing potential differences in the function of each protein.

Sharks have a small olfactory receptor repertoire compared to many bony vertebrates, e.g. the main olfactory receptor family of bony vertebrates, the ORs, is contracted in sharks and consists of only two OR genes plus seven OR-like genes (Syed et al. 2023). In bony vertebrates OR genes are expressed by ciliated neurons, so the loss of OR genes and loss of OR expression in the olfactory epithelium is consistent with the putative secondary loss of ciliated neurons in sharks (Ferrando and Gallus 2013). We find that the majority of OR genes are most highly expressed in other organs, suggesting function in the immune system (spleen) or other sensory organs (ampullae of Lorenzini). Similarly, three even smaller families of olfactory receptors, TAARs, TARLs (olfactory in cyclostomes only), and ORAs, are not enriched in olfactory tissue and all of them have higher expression levels in different other tissues. ORAs are supposed to be ancestrally olfactory receptors, as they are partially olfactory in cyclostomes (Kowatschew and Korsching 2022) and olfactory in bony fish; neofunctionalisation outside of the olfactory system in cartilaginous fish therefore involved the loss of the ancestral function. The catshark V2R-like genes appear to have a broad spectrum of functions both in the olfactory epithelium and elsewhere, including other sensory organs such as the eye. In contrast, nearly all V2R genes (considered the main olfactory receptor family of sharks (Syed et al. 2023)) show unique enrichment in the ‘brain & olfactory epithelium’ tissue, consistent with other expression data (Syed et al. 2023). This supports specific function in olfaction only for V2R genes, although additional and sometimes exclusive expression of some V2Rs in non-olfactory tissues suggest some degree of neofunctionalisation (e.g. in ionic homeostasis), even in the only family considered to be dedicated to olfactory function in the catshark.

To summarise, the catshark (and possibly elasmobranchs in general) appear to use primarily a single gene family for the detection of odours (the V2Rs) out of the four canonical vertebrate olfactory receptor gene families. This represents a nearly inverse situation compared to the jawless vertebrate lamprey, whose V2Rs are not expressed in the olfactory epithelium, in contrast to ORs, TAAR-like genes and V1Rs (Kowatschew and Korsching 2022), highlighting the need for a focused examination of the evolution of their function in chondrichthyans. The expression patterns of several genes in the olfactory neuron layer are suggestive of sub-populations in these sensory cells, and axons coming from medial or lateral lamellae project through two separate olfactory nerves: both these observations support a structural complexity that should be uncovered by further functional studies of the shark olfactory system.

#### Electrosensory system: Catshark provides insight into ancestral state of electroreception

We showed that the 3D spatial organisation of the ampullae of Lorenzini is already present in embryonic stages of the small-spotted catshark together with histological differentiation of cells in the sensory, versus supporting, versus canal zones of the organ. Nervous connections between the different clusters of ampullae and the anterior lateral line nerves appear to coalesce with sensory neurons connected to the sensory patches of the lateral line. Previous analyses of genes involved in electrosensory systems in vertebrates have focused on electrophysiology (e.g. in elasmobranch fishes, Bellono et al. 2017) or differential gene expression between the skin and ampullary organ tissue (e.g. in elasmobranch fishes, Bellono et al. 2017; and in bony fish, Modrell et al. 2017). Our analysis of transcriptomic data supports the identification of ribbon-synapse components in electrosensory cells of the catshark, similar to what was proposed for the paddlefish (Modrell et al. 2017), with expression of glutamate transporters, L-type voltage-gated calcium channels (Ca_v_1 channels) and the transmembrane protein Otoferlin (Modrell et al. 2017; Manchanda et al. 2021). In the paddlefish, little transcriptomic data differentiated hair cells from electrosensory cells with the exception of two potassium voltage-gated channels described to be specific to the ampullary organs (Modrell et al. 2017). We detected high and specific expression of the orthologous potassium voltage-gated channel *kcnab3* gene in the catshark ampullae of Lorenzini, suggesting this is an ancestral characteristic for all jawed vertebrates.

Our results also suggest that some TRPs might be expressed in sensory organs but with an ion-transport function instead of a sensory function (e.g. *trpv8.2* in the ampullae of Lorenzini and the olfactory epithelium). The ampullae of Lorenzini display their own combination of expressed TRPs (Table 3), making the members of this gene family excellent candidates for further description of signal transduction in the electrosensory system of vertebrates.

## CONCLUSION

Despite the paramount importance of chondrichthyans in vertebrate evolution, their poor suitability to genetic approaches due to their large size and long generation times have been a major limitation in their development as biological models. However, the availability of an increasing number of chromosome-level genomes in this clade marks its entry in the post-genomic era, with novel perspectives in multiple thematic fields, including not only comparative genomics but also population genetics, physiology, neuroscience, and evolutionary developmental biology. Combinations of epigenomics, single-cell and spatial transcriptomics are currently revolutionising our understanding of the cellular diversity of embryonic and adult organs. These techniques are fully applicable to chondrichthyans, with large organ size even becoming advantageous in spatially resolved approaches. Due to its experimental accessibility and key phylogenetic position, the small-spotted catshark is an excellent model to understand the origin and development of cell-type and organ complexity during vertebrate evolution.

## ACKNOWLEDGEMENTS

TB was supported by a studentship from the Biotechnology and Biological Sciences Research Council-funded South West Biosciences Doctoral Training Partnership (BB/M009122/1). AH was supported by a Biotechnology and Biological Sciences Research Council (BBSRC) David Phillips Fellowship (BB/N020146/1). E.L.A. was supported by grants from the Welch Foundation (Q-1866), an NIH Encyclopedia of DNA Elements Mapping Center Award (UM1HG009375), a US-Israel Binational Science Foundation Award (2019276), the Behavioral Plasticity Research Institute (NSF DBI-2021795), NSF Physics Frontiers Center Award (NSF PHY-2210291), and an NIH CEGS (RM1HG011016-01A1). Genome assembly was performed in association with the DNA Zoo Consortium (www.dnazoo.org), which acknowledges support from Illumina, IBM, and Pawsey Supercomputing Center.

We thank Dr. Paul Tafforeau and Dr. Kathleen N. Dollman at the European Synchrotron Radiation Facility for their assistance in the data acquisition and tomographic reconstruction. Beamline at BM05 was granted as part of proposal LS3021. We acknowledge ABIMS and Genotoul for access to bioinformatic facilities, Pascal Romans and the Aquariology Service of the Banyuls Oceanological Observatory (OOB) for providing catshark specimens. Catshark infrastructures of OOB and collaborations were funded by EMBRC-France and EMBRC-Europe. To generate the histology data, we acknowledge the “Réseau d’Histologie Expérimentale de Montpellier” - RHEM facility supported by SIRIC Montpellier Cancer Grant INCa_Inserm_DGOS_12553, the European regional development foundation and the Occitanie region (FEDER-FSE 2014-2020 Languedoc Roussillon), REACT-EU (Recovery Assistance for Cohesion and the Territories of Europe), IBiSA and Ligue contre le cancer for histology technics.

