## Supplementary for "The sensory shark: high-quality morphological, genomic and transcriptomic data for the small-spotted catshark *Scyliorhinus canicula* reveal the molecular bases of sensory organ evolution in jawed vertebrates"

This is the list of supplementary materials for the manuscript "The sensory shark: high-quality phenotypic, genomic and transcriptomic data for the small spotted catshark *Scyliorhinus canicula* reveals the molecular bases of sensory organs evolution in jawed vertebrates" from Mayeur et al.

### SUPPLEMENTARY MATERIAL: Supplementary Dataset List (separate files, available on request)

#### Supplementary Dataset 1 SDataset1.xls

Table of all TPM values for all genes in all organs/developmental stages. When more than one RNA library was available for an organ, their values were averaged.

#### Supplementary Dataset 2 SDataset2.xls

Table of Z-scores obtained by comparison of TPM values for all genes in all organs/developmental stages.

#### Supplementary Dataset 3 SDataset3.xls

Table of Z-scores obtained by comparison of TPM values for all genes in embryonic developmental stages.

#### Supplementary Dataset 4 SDataset4.xls

Table of Z-scores obtained by comparison of TPM values for all genes in adult tissues.

#### Supplementary Dataset 5 SDataset5.xls

Autocorrelation value within embryonic stages for each gene.

#### Supplementary Dataset 6 SDataset6.xls

GOterm association analysis in embryonic stage RNAseq data.

#### Supplementary Dataset 7 SDataset7.xls

List of most highly and most specifically expressed genes in each adult tissue RNAseq.

#### Supplementary Dataset 8 SDataset8.xls

GOterm association analysis in adult tissues.

#### Supplementary Dataset 9 SDataset9.avi

Video to see olfactory nerve, frontal view.

#### Supplementary Dataset 10 SDataset10.avi

Video to see olfactory nerve, ventral view.

#### Supplementary Dataset 11 SDataset11.xls

Table Top-50 genes in 3 sensory organs

### SUPPLEMENTARY MATERIAL: Supplementary Table List

#### Supplementary Table 1 List of RNA libraries generated on three different adult individuals (named male1, male2, female) and six individual embryos and the detail of organs sampled for each library.

| NCBI SRA reference | Individual | Organ |
|---|---|---|
| ERX5970093 | male1 | ampullae of Lorenzini |
| ERX5970102 | female | ampullae of Lorenzini |
| ERX5970120 | male1 | blood |
| ERX5970123 | female | blood |
| ERX5970075 | male1 | brain & olfactory epithelium |
| ERX5970090 | female | brain & olfactory epithelium |
| ERX5970101 | male2 | brain & olfactory epithelium |
| ERX5970078 | male2 | chondrocranium |
| ERX5970095 | male1 | chondrocranium |
| ERX5970114 | female | chondrocranium |
| ERX5970077 | male1 | dental lamina |
| ERX5970097 | female | dental lamina |
| ERX5970112 | male2 | dental lamina |
| ERX5970082 | male1 | esophagus |
| ERX5970087 | male1 | eye |
| ERX5970096 | female | eye |
| ERX5970076 | male1 | gills |
| ERX5970110 | male1 | heart |
| ERX5970115 | male1 | hypaxial muscle |
| ERX5970118 | female | hypaxial muscle |
| ERX5970121 | male1 | kidney |
| ERX5970106 | male1 | liver |
| ERX5970089 | male1 | Meckel's cartilage |
| ERX5970109 | female | Meckel's cartilage |
| ERX5970122 | male2 | Meckel's cartilage |
| ERX5970108 | female | ovary |
| ERX5970091 | female | pancreas |
| ERX5970116 | male1 | pancreas |
| ERX5970085 | female | rectal gland |
| ERX5970100 | male1 | rectal gland |
| ERX5970105 | male1 | seminal vesicle |
| ERX5970083 | male1 | skin denticle |
| ERX5970103 | female | skin denticle |
| ERX5970117 | male2 | skin denticle |
| ERX5970081 | male1 | spinal cord |
| ERX5970079 | female | spiral intestine |
| ERX5970094 | male1 | spiral intestine |
| ERX5970111 | male1 | spleen |
| ERX5970088 | male1 | stomach |
| ERX5970099 | male1 | testis |
| ERX5970107 | male2 | testis |
| ERX5970113 | female | uterus |
| ERX5970084 | male2 | vertebra |
| ERX5970119 | female | vertebra |
| ERX5970124 | embryo st12 | stage 12 |
| ERX5970080 | embryo st22 | stage 22 |
| ERX5970086 | embryo st24 | stage 24 |
| ERX5970092 | embryo st26 | stage 26 |
| ERX5970098 | embryo st30 | stage 30 |
| ERX5970104 | embryo st31 | stage 31 |

Supplementary Table 2 Assembly accession numbers and species names used in genome structure analysis

| Assembly Accession | Species name |
|---|---|
| GCA_000001405 | *Homo sapiens* |
| GCA_000242695 | *Lepisosteus oculatus* |
| GCA_004010195 | *Chiloscyllium plagiosum* |
| GCA_009764475 | *Pristis pectinate* |
| GCA_010909765 | *Amblyraja radiata* |
| GCA_017639515 | *Carcharodon carcharias* |
| GCA_018977255 | *Callorhinchus milii* |
| GCA_020745735 | *Hemiscyllium ocellatum* |
| GCA_021869965 | *Rhincodon typus* |
| GCA_022316705 | *Stegostoma fasciatum* |
| GCA_028641065 | *Leucoraja erinacea* |
| GCA_030035685 | *Mobula birostris* |
| GCA_030144855 | *Hypanus sabinus* |
| GCA_030390025 | *Squalus acanthias* |
| GCA_902713615 | *Scyliorhinus canicula* |
| GCF_010993605 | *Petromyzon marinus* |

Supplementary Table 3 Hox query sequences used to search the catshark genome assembly.

| Accession | Gene | Species |
|---|---|---|
| ACU32554.1 | homeobox protein HoxA1 | *Callorhinchus milii* |
| ACU32546.1 | homeobox protein HoxA10 | *Callorhinchus milii* |
| ACU32545.1 | homeobox protein HoxA11 | *Callorhinchus milii* |
| ACU32544.1 | homeobox protein HoxA13 | *Callorhinchus milii* |
| ACU32553.1 | homeobox protein HoxA2 | *Callorhinchus milii* |
| ACU32552.1 | homeobox protein HoxA3 | *Callorhinchus milii* |
| ACU32551.1 | homeobox protein HoxA4 | *Callorhinchus milii* |
| ACU32550.1 | homeobox protein HoxA5 | *Callorhinchus milii* |
| ACU32549.1 | homeobox protein HoxA6 | *Callorhinchus milii* |
| ACU32548.1 | homeobox protein HoxA7 | *Callorhinchus milii* |
| ACU32547.1 | homeobox protein HoxA9 | *Callorhinchus milii* |
| ACU32555.1 | homeobox protein HoxB1 | *Callorhinchus milii* |
| ACU32565.1 | homeobox protein HoxB10 | *Callorhinchus milii* |
| ACU32564.1 | homeobox protein HoxB13 | *Callorhinchus milii* |
| ACU32556.1 | homeobox protein HoxB2 | *Callorhinchus milii* |
| ACU32557.1 | homeobox protein HoxB3 | *Callorhinchus milii* |
| ACU32558.1 | homeobox protein HoxB4 | *Callorhinchus milii* |
| ACU32559.1 | homeobox protein HoxB5 | *Callorhinchus milii* |
| ACU32560.1 | homeobox protein HoxB6 | *Callorhinchus milii* |
| ACU32561.1 | homeobox protein HoxB7 | *Callorhinchus milii* |
| ACU32562.1 | homeobox protein HoxB8 | *Callorhinchus milii* |
| ACU32563.1 | homeobox protein HoxB9 | *Callorhinchus milii* |
| ACU32578.1 | homeobox protein HoxC1 | *Callorhinchus milii* |
| ACU32569.1 | homeobox protein HoxC10 | *Callorhinchus milii* |
| ACU32570.1 | homeobox protein HoxC11 | *Callorhinchus milii* |
| ACU32571.2 | homeobox protein HoxC12 | *Callorhinchus milii* |
| ACU32572.1 | homeobox protein HoxC13 | *Callorhinchus milii* |
| ACU32575.1 | homeobox protein HoxC3 | *Callorhinchus milii* |
| ACU32574.1 | homeobox protein HoxC4 | *Callorhinchus milii* |
| ACU32576.1 | homeobox protein HoxC5 | *Callorhinchus milii* |
| ACU32577.1 | homeobox protein HoxC6 | *Callorhinchus milii* |
| ACU32573.1 | homeobox protein HoxC8 | *Callorhinchus milii* |
| ACU32568.1 | homeobox protein HoxC9 | *Callorhinchus milii* |
| ACU32592.1 | homeobox protein HoxD1 | *Callorhinchus milii* |
| ACU32585.1 | homeobox protein HoxD10 | *Callorhinchus milii* |
| ACU32584.1 | homeobox protein HoxD11 | *Callorhinchus milii* |
| ACU32583.1 | homeobox protein HoxD12 | *Callorhinchus milii* |
| ACU32582.1 | homeobox protein HoxD13 | *Callorhinchus milii* |
| ACU32581.1 | homeobox protein HoxD14 | *Callorhinchus milii* |
| ACU32591.1 | homeobox protein HoxD2 | *Callorhinchus milii* |
| ACU32590.1 | homeobox protein HoxD3 | *Callorhinchus milii* |
| ACU32589.1 | homeobox protein HoxD4 | *Callorhinchus milii* |
| ACU32588.1 | homeobox protein HoxD5 | *Callorhinchus milii* |
| ACU32587.1 | homeobox protein HoxD8 | *Callorhinchus milii* |
| ACU32586.1 | homeobox protein HoxD9 | *Callorhinchus milii* |
| XP_041039348.1 | homeobox protein Hox-A1 | *Carcharodon carcharias* |
| XP_041040401.1 | homeobox protein Hox-A11b | *Carcharodon carcharias* |
| XP_041040525.1 | homeobox protein Hox-A13b | *Carcharodon carcharias* |
| XP_041039347.1 | homeobox protein Hox-A2b | *Carcharodon carcharias* |
| XP_041073562.1 | homeobox protein Hox-A2-like | *Carcharodon carcharias* |
| XP_041057354.1 | homeobox protein Hox-A2-like | *Carcharodon carcharias* |
| XP_041039346.1 | homeobox protein Hox-A3 | *Carcharodon carcharias* |
| XP_041039344.1 | homeobox protein Hox-A3 | *Carcharodon carcharias* |
| XP_041039343.1 | homeobox protein Hox-A3 | *Carcharodon carcharias* |
| XP_041039342.1 | homeobox protein Hox-A3 | *Carcharodon carcharias* |
| XP_041039341.1 | homeobox protein Hox-A3 | *Carcharodon carcharias* |
| XP_041039340.1 | homeobox protein Hox-A3 | *Carcharodon carcharias* |
| XP_041039339.1 | homeobox protein Hox-A3 | *Carcharodon carcharias* |
| XP_041039338.1 | homeobox protein Hox-A3 | *Carcharodon carcharias* |
| XP_041039337.1 | homeobox protein Hox-A3 | *Carcharodon carcharias* |
| XP_041039336.1 | homeobox protein Hox-A3 | *Carcharodon carcharias* |
| XP_041056252.1 | homeobox protein Hox-A4-like | *Carcharodon carcharias* |
| XP_041039349.1 | homeobox protein Hox-A5 | *Carcharodon carcharias* |
| XP_041040938.1 | homeobox protein Hox-A7 | *Carcharodon carcharias* |
| XP_041040251.1 | homeobox protein Hox-A9 | *Carcharodon carcharias* |
| XP_041029720.1 | homeobox protein Hox-B10a | *Carcharodon carcharias* |
| XP_041029457.1 | homeobox protein Hox-B1a | *Carcharodon carcharias* |

| | | |
|---|---|---|
| XP_041073561.1 | homeobox protein Hox-B3a-like | *Carcharodon carcharias* |
| XP_041073560.1 | homeobox protein Hox-B3a-like | *Carcharodon carcharias* |
| XP_041029680.1 | homeobox protein Hox-B4a | *Carcharodon carcharias* |
| XP_041029678.1 | homeobox protein Hox-B5a isoform X1 | *Carcharodon carcharias* |
| XP_041029679.1 | homeobox protein Hox-B5a isoform X2 | *Carcharodon carcharias* |
| XP_041029683.1 | homeobox protein Hox-B7a | *Carcharodon carcharias* |
| XP_041029298.1 | homeobox protein Hox-B8a isoform X1 | *Carcharodon carcharias* |
| XP_041029299.1 | homeobox protein Hox-B8a isoform X2 | *Carcharodon carcharias* |
| XP_041029297.1 | homeobox protein Hox-B9a | *Carcharodon carcharias* |
| XP_041036501.1 | homeobox protein Hox-C10-like, partial | *Carcharodon carcharias* |
| XP_041036500.1 | homeobox protein Hox-C11-like | *Carcharodon carcharias* |
| XP_041036496.1 | homeobox protein Hox-C13-like | *Carcharodon carcharias* |
| XP_041036503.1 | homeobox protein Hox-C6-like | *Carcharodon carcharias* |
| XP_041036502.1 | homeobox protein Hox-C9-like | *Carcharodon carcharias* |
| XP_041057237.1 | homeobox protein Hox-D11a | *Carcharodon carcharias* |
| XP_041057220.1 | homeobox protein Hox-D13 | *Carcharodon carcharias* |
| XP_041057355.1 | homeobox protein Hox-D1-like | *Carcharodon carcharias* |
| XP_041056706.1 | homeobox protein Hox-D3a-like isoform X1 | *Carcharodon carcharias* |
| XP_041056705.1 | homeobox protein Hox-D3a-like isoform X1 | *Carcharodon carcharias* |
| XP_041056707.1 | homeobox protein Hox-D4a-like isoform X2 | *Carcharodon carcharias* |
| XP_041056708.1 | homeobox protein Hox-D4a-like isoform X3 | *Carcharodon carcharias* |
| XP_041056709.1 | homeobox protein Hox-D5 isoform X1 | *Carcharodon carcharias* |
| XP_041056710.1 | homeobox protein Hox-D5 isoform X2 | *Carcharodon carcharias* |
| XP_041056983.1 | homeobox protein Hox-D8 | *Carcharodon carcharias* |
| XP_041056993.1 | homeobox protein Hox-D9a | *Carcharodon carcharias* |
| XP_041057239.1 | LOW QUALITY PROTEIN: homeobox protein Hox-D10a | *Carcharodon carcharias* |
| P49639.2 | Homeobox protein Hox-A1 | *Homo sapiens* |
| P31260.3 | Homeobox protein Hox-A10 | *Homo sapiens* |
| P31270.2 | Homeobox protein Hox-A11 | *Homo sapiens* |
| P31271.3 | Homeobox protein Hox-A13 | *Homo sapiens* |
| O43364.1 | Homeobox protein Hox-A2 | *Homo sapiens* |
| O43365.1 | Homeobox protein Hox-A3 | *Homo sapiens* |
| Q00056.3 | Homeobox protein Hox-A4 | *Homo sapiens* |
| P20719.2 | Homeobox protein Hox-A5 | *Homo sapiens* |
| P31267.2 | Homeobox protein Hox-A6 | *Homo sapiens* |
| P31268.3 | Homeobox protein Hox-A7 | *Homo sapiens* |
| P31269.4 | Homeobox protein Hox-A9 | *Homo sapiens* |
| P14653.2 | Homeobox protein Hox-B1 | *Homo sapiens* |
| Q92826.2 | Homeobox protein Hox-B13 | *Homo sapiens* |
| P14652.1 | Homeobox protein Hox-B2 | *Homo sapiens* |
| P14651.2 | Homeobox protein Hox-B3 | *Homo sapiens* |
| P17483.2 | Homeobox protein Hox-B4 | *Homo sapiens* |
| P09067.3 | Homeobox protein Hox-B5 | *Homo sapiens* |
| P17509.4 | Homeobox protein Hox-B6 | *Homo sapiens* |
| P09629.4 | Homeobox protein Hox-B7 | *Homo sapiens* |
| P17481.2 | Homeobox protein Hox-B8 | *Homo sapiens* |
| P17482.2 | Homeobox protein Hox-B9 | *Homo sapiens* |
| Q9NYD6.2 | Homeobox protein Hox-C10 | *Homo sapiens* |
| O43248.1 | Homeobox protein Hox-C11 | *Homo sapiens* |
| P31275.2 | Homeobox protein Hox-C12 | *Homo sapiens* |
| P31276.3 | Homeobox protein Hox-C13 | *Homo sapiens* |
| P09017.2 | Homeobox protein Hox-C4 | *Homo sapiens* |
| Q00444.1 | Homeobox protein Hox-C5 | *Homo sapiens* |
| P09630.3 | Homeobox protein Hox-C6 | *Homo sapiens* |
| P31273.2 | Homeobox protein Hox-C8 | *Homo sapiens* |
| P31274.3 | Homeobox protein Hox-C9 | *Homo sapiens* |
| Q9GZZ0.1 | Homeobox protein Hox-D1 | *Homo sapiens* |
| P28358.2 | Homeobox protein Hox-D10 | *Homo sapiens* |
| P31277.3 | Homeobox protein Hox-D11 | *Homo sapiens* |
| P35452.3 | Homeobox protein Hox-D12 | *Homo sapiens* |
| P35453.3 | Homeobox protein Hox-D13 | *Homo sapiens* |
| P31249.3 | Homeobox protein Hox-D3 | *Homo sapiens* |
| P09016.3 | Homeobox protein Hox-D4 | *Homo sapiens* |
| P13378.2 | Homeobox protein Hox-D8 | *Homo sapiens* |
| P28356.5 | Homeobox protein Hox-D9 | *Homo sapiens* |
| CBL59343.1 | HoxA1 | *Scyliorhinus canicula* |
| CBL59335.1 | HoxA10 | *Scyliorhinus canicula* |
| CBL59334.1 | HoxA11 | *Scyliorhinus canicula* |
| CBL59333.1 | HoxA13 | *Scyliorhinus canicula* |
| CBL59342.1 | HoxA2 | *Scyliorhinus canicula* |
| CBL59341.1 | HoxA3 | *Scyliorhinus canicula* |
| CBL59340.1 | HoxA4 | *Scyliorhinus canicula* |

| | | |
|---|---|---|
| CBL59339.1 | HoxA5 | *Scyliorhinus canicula* |
| CBL59338.1 | HoxA6 | *Scyliorhinus canicula* |
| CBL59337.1 | HoxA7 | *Scyliorhinus canicula* |
| CBL59336.1 | HoxA9 | *Scyliorhinus canicula* |
| CBL59354.1 | HoxB1 | *Scyliorhinus canicula* |
| CBL59345.1 | HoxB10 | *Scyliorhinus canicula* |
| CBL59344.1 | HoxB13 | *Scyliorhinus canicula* |
| CBL59353.1 | HoxB2 | *Scyliorhinus canicula* |
| CBL59352.1 | HoxB3 | *Scyliorhinus canicula* |
| CBL59351.1 | HoxB4 | *Scyliorhinus canicula* |
| CBL59349.1 | HoxB6 | *Scyliorhinus canicula* |
| CBL59348.1 | HoxB7 | *Scyliorhinus canicula* |
| CBL59347.1 | HoxB8 | *Scyliorhinus canicula* |
| CBL59346.1 | HoxB9 | *Scyliorhinus canicula* |
| CBL59367.1 | HoxD1 | *Scyliorhinus canicula* |
| CBL59360.1 | HoxD10 | *Scyliorhinus canicula* |
| CBL59359.1 | HoxD11 | *Scyliorhinus canicula* |
| CBL59358.1 | HoxD12 | *Scyliorhinus canicula* |
| CBL59357.1 | HoxD13 partial | *Scyliorhinus canicula* |
| CBL59356.1 | HoxD14 | *Scyliorhinus canicula* |
| CBL59366.1 | HoxD2 | *Scyliorhinus canicula* |
| CBL59365.1 | HoxD3 | *Scyliorhinus canicula* |
| CBL59364.1 | HoxD4 | *Scyliorhinus canicula* |
| CBL59363.1 | HoxD5 | *Scyliorhinus canicula* |
| CBL59362.1 | HoxD8 | *Scyliorhinus canicula* |
| CBL59361.1 | HoxD9 | *Scyliorhinus canicula* |
| ASS31193.1 | homeobox protein Hox-A1 | *Scyliorhinus torazame* |
| ASS31201.1 | homeobox protein Hox-A10 | *Scyliorhinus torazame* |
| ASS31202.1 | homeobox protein Hox-A11 | *Scyliorhinus torazame* |
| ASS31203.1 | homeobox protein Hox-A13 | *Scyliorhinus torazame* |
| ASS31194.1 | homeobox protein Hox-A2 | *Scyliorhinus torazame* |
| ASS31195.1 | homeobox protein Hox-A3 | *Scyliorhinus torazame* |
| ASS31196.1 | homeobox protein Hox-A4 | *Scyliorhinus torazame* |
| ASS31197.1 | homeobox protein Hox-A5 | *Scyliorhinus torazame* |
| ASS31198.1 | homeobox protein Hox-A6 | *Scyliorhinus torazame* |
| ASS31199.1 | homeobox protein Hox-A7 | *Scyliorhinus torazame* |
| ASS31200.1 | homeobox protein Hox-A9 | *Scyliorhinus torazame* |
| ASS31204.1 | homeobox protein Hox-B1 | *Scyliorhinus torazame* |
| ASS31212.1 | homeobox protein Hox-B10 | *Scyliorhinus torazame* |
| ASS31213.1 | homeobox protein Hox-B13 | *Scyliorhinus torazame* |
| ASS31205.1 | homeobox protein Hox-B2 | *Scyliorhinus torazame* |
| ASS31206.1 | homeobox protein Hox-B3 | *Scyliorhinus torazame* |
| ASS31207.1 | homeobox protein Hox-B4 | *Scyliorhinus torazame* |
| ASS31208.1 | homeobox protein Hox-B5 | *Scyliorhinus torazame* |
| ASS31209.1 | homeobox protein Hox-B6 | *Scyliorhinus torazame* |
| ASS31210.1 | homeobox protein Hox-B8 | *Scyliorhinus torazame* |
| ASS31211.1 | homeobox protein Hox-B9 | *Scyliorhinus torazame* |
| ASS31214.1 | homeobox protein Hox-D1 | *Scyliorhinus torazame* |
| ASS31225.1 | homeobox protein Hox-D1, partial | *Scyliorhinus torazame* |
| ASS31221.1 | homeobox protein Hox-D10 | *Scyliorhinus torazame* |
| ASS31222.1 | homeobox protein Hox-D11 | *Scyliorhinus torazame* |
| ASS31223.1 | homeobox protein Hox-D12 | *Scyliorhinus torazame* |
| ASS31224.1 | homeobox protein Hox-D13 | *Scyliorhinus torazame* |
| ASS31215.1 | homeobox protein Hox-D2, partial | *Scyliorhinus torazame* |
| ASS31216.1 | homeobox protein Hox-D3 | *Scyliorhinus torazame* |
| ASS31217.1 | homeobox protein Hox-D4 | *Scyliorhinus torazame* |
| ASS31218.1 | homeobox protein Hox-D5 | *Scyliorhinus torazame* |
| ASS31219.1 | homeobox protein Hox-D8 | *Scyliorhinus torazame* |
| ASS31220.1 | homeobox protein Hox-D9 | *Scyliorhinus torazame* |

#### Supplementary Table 4 List of RNAseq data used to build the *'ncbi-UTRs'* reference gene model

| Identification | Origin of cDNAs | Sequencing | Number of reads |
|---|---|---|---|
| To be submitted | whole embryo | Illumina Solexa, paired | 729,461,418 |
| **SRX22532267-365** | Habenula, stage 18 head | Illumina Hi-Seq 2500, paired | 946,712,650 |
| **SRX2495300-1** | developing jaw | Illumina Hi-seq 1500, paired | 290,232,228 |
| **SRX651773-5** | pancreas, brain and liver | Illumina MiSeq + Illumina HiSeq 2500, paired | 101,136,796 |
| **SRX036537** | whole embryo, stages 24 to 30 | Illumina Genome Analyzer II, non-paired | 50,760,902 |
| **SRX11963304-400** | embryo stages 11-34 | Cel-Seq2, Illumina, paired | 611,794,460 |
| To be submitted | embryo stages 15-21 | Illumina, paired | 932,528,270 |
| **SRR8753342** | 7cm embryo vertebrae | Illumina, paired | 537,310,337 |
| To be submitted | juvenile nose | Illumina, paired | 204,831,376 |
| **ERX5970075-124** | embryo and adult (this study) | Illumina, paired | 5,764,089,680 |
| **SRR8179289-91** | mature and immature gonads | Illumina HiSeq 2000, paired | 168,389,576 |
| **SRR8077742** | multiple tissues | Ion Torrent Proton, non-paired | 134,841,605 |

#### Supplementary Table 5 DNA matrix sequences used for RNA probes synthesis

First 23bp are the sequence for T3 primer (blue); last 23bp are the sequence for T7 primer (red).

| Target gene | Sequence (blue: T3 sequence; red: T7 sequence) |
|---|---|
| *otof* | CGAAATTAACCCTCACTAAAGGGTGCAAGTCAACGCCCCTAACAGCTGGATGAAGTCAATTGGCGCCTATTGAATGACAGTAAATATGATAGTCTAAG<br>TAGCTTGGAAAAAAAACAATATATGTCAACGGTGCACGAAACCCGCTCTGTTCATAAAGCCTCTAAATTATTTAGGGACGGCGAACAGCAAACGGGAT<br>GGACTGCAGTGAGTTGCACTGAGAGGAGGAGACGCTGGGCAGCAGTTTCTCTCGCCGCGGTGTGCCTGAACTGGCCGGGAACAGCTAGTCAGAGCGCA<br>GCATCCCAGAGCAACACTGTAACTGTGGCCAGGCTCAGCGCTCAGCATGGCTCTCATGGTCCACCTGAAAACGGTCACCGATCTCCGAGGGAAAGGCG<br>ACAGGATCGCCAAAGTAGCCTTTCGAGGGCTCTCATTTTATACAAGAGTACTCGAAAACTGTGAAGAAGAAGCACAATTTGATGAGACTTTCCGATGG<br>CCTATTGCCAGTAATATTGATGGGAATGAGATGCTGGAGATTCAAGTTTACAACTACAGCAAAGTCTTTACAAATAGATTAATTGGGACTTTCGGGAT<br>GGTATTACAGAAGGTAGTGGAAGAAGGACATCTGGAAATTGCAGACACACTAATAGATGACAACAACAGCTCAATCAGGACCTGTGTCTCCATTGAGA<br>TAAAATACCAGGCAATGGATGGTACTGTTGGAGCCTGGAATGAAAGTGATTTCTTAGAAACCCTATAGTGAGTCGTATTACAA |
| *slc1a3* | CGAAATTAACCCTCACTAAAGGGGGTCTGCTCATTCACGCCCTCATCGTGCTGCCGCTTCTCTACTTTGCCATCACACACAAGAATCCCTTTGTTTTC<br>ATTGGTGGATTGGTACAGGCCCTGGTTACTGCACTCGGCACTTCCTCCAGCTCAGCCACCTTGCCGATCACCTTTCGTTGCCTGGAGGACAAGATTGG<br>GCTGGATAAACGAATCACCAGATTTGTGTTGCCTGTTGGTGCCACCATTAACATGGATGGCACTGCCTTGTACGAAGCTCTTGCTGCCATCTTTATCG<br>CACAAGTTAACAACTTTGAACTCAACTTTGGACAAATCCTCACTATCAGTATCACTGCCACAGCTGCTAGTATTGGTGCAGCCGGCATTCCGCAGGCA<br>GGCCTGGTAACCATGGTAATCGTCCTGACATCTGTGGGGCTTCCAACAGACGACATCACGTTGATCATCGCCGTGGATTGGTTTCTGGACCGGCTGCG<br>TACGATGACCAATGTGCTGGGAGACTCCCTGGGAGCCGGGATCGTGGAGCACCTCTCACGCCATGAACTTCAGAAAATGGACGCCGAGGCTCTCAACT<br>CTGTCATTGAGGAAAATGAGAGGAAGAAACCCTATCAGCTGATCGGTCAGGACAACGAGAAACCAACAGACAATGAGACCAAGATGTAGCCACCGTGT<br>TGAAGCTACCTTCCTGACAGACAGTCTCTTCCACTATCTGCATTGAGGCTGATCAGATTGGCCCTGCTGCGCCTTCCCTATAGTGAGTCGTATTACAA |
| *vglut3*<br>=<br>*slc17a8* | CGAAATTAACCCTCACTAAAGGGAGGAAATTGAGTTGAATGAAGATGGTCAGCCGGTGAAGACGGCAGTATTGAAGCAGCCATTATGTGATTGTCACT<br>GCTTTGGGATGCCCAAGCGTTACATGATTGCCATCATGAGTGGACTTGGATTCTGCATCTCCTTCGGCATCAGGTGCAACTTGGGCGTGGCAATAGTT<br>GAAATGGTCAACAACAACACCGTCTATGTGAATGGCAAGGCAGAGATTGAGAAACCTCAATTCAACTGGAACCCTGAGACTGTAGGGTTGATTCATGG<br>GTCCTTTTTCTGGGGCTATATTGTAACTCAAATCCCAGGGGGATTTATCTCCAACAAGTTTGCTGCAAACAGAGTTTTTGGAGTTGCTATATTTTTGA<br>CCTCAACACTGAACATGCTCATCCCATCAGCAGCAAGAATACATTTTGGCTGTGTTTTGTTTGTAAGGATACTTCAAGGCCTTGTGGAGGGTGTCACC<br>TACCCAGCATGCCATGGGCTCTGGAGCATGTGGGCCCCACCACTTGAAAGGAGTCGATTGGCTACTACATCTTTCTGTGGTTCTTACGCAGGGGCTGT<br>TGTGGCAATGCCACTCGCAGGATTGTTGGTCCAGTACAAAGGCTGGTCTTCTGCCCTATAGTGAGTCGTATTACAA |
| *moxd2.1* | CGAAATTAACCCTCACTAAAGGGATGGGAACAGGTGTCGTTGACAATTGTGGATTTAAATTACATCTGACGACTAATCTTCGTAAATTTGACGTTGGG<br>ATTTTCTGGACGGGTGTTCAAGTTGCTGAGTTCTTAGTCATCCCTCCAAAGGCATCATCCTTCAAGACCTATGCATATTGTGACACATCTCCTGTTGA<br>TAAGGAGGAAGGCAAGAAATATACGGATATGCAAATCTTTGGATCGATACTCCACACGCACCTTACAGGGTCTAAAGTTCGCATCTTGCAGTTCAGAG<br>GTGGGGAGCAGATCAGGACAATTGCGCAGGACAAATATTATGATTTTAATCTACAGGAGTCACGGATGCTCAGGGAGCCCGTCACCGTGCGCCGGGGG<br>GATGTGCTGATTACAGAATGCACATATAACACGGAGAATCGATCAAACATAACGTGGGGTGGATTGGCAACAACTGAGGAAATGTGTCTCGCCTTCAT<br>GTTCTATTATCCAAAGAATGAACTGCTGACGTGTTTGAGTTATGTGACAGCTCGCACTGCTTCAAGTGTCTTGAACATCACAGGGGACACGACCGAAG<br>AGATAATCACAAGCATGTGGTCATGGGACTGGAACCAGACTCTCATTGACGAGGTGGAGACCAAACTGCGAGATGCCGATCAGGATCAGGTCATCGGC<br>ACCAACACGGCTTACAAACACTTTAAGGGTAAGAAGACCCTATAGTGAGTCGTATTACAA |
| *moxd2.2* | CGAAATTAACCCTCACTAAAGGGATGAGAGATGGGAAACAGATCGGAATCCTTGCACAAGATACAAACTACGATTTCAATCTTCAGGAAACAAAAACT<br>TTGAAGGAACCAGCCCGTGTGAAGAGGGGGGATGTGCTGATTACAGAATGCACATATAACACGGAGAATCGATCAACAATAACGTGGGGTGGATTGAG<br>TACCACTGATGAAATGTGTCTTGCCTACATGTGGTATTACCCCCGGATTGAGTTCACCGGCTGCAACAGTTTCCAGACGTTTCAGACAGTGGCTGCAG<br>CTTTGAATGTTTCAGCAACCAGCAAAGAAGATCTGTTCGCGATATTGCAGAATTGGAACTGGACAGAAAGATCTGCCGCAGACATGGAAGTGAAGCTC<br>CGTGATGCCGATCACTACCTGTTCACCATGGATCGCACTGGTCACCGATTCACCAAGGTGAAGATTCCGGAGATCCCGGAGGAGAAAGTAATCCCATG<br>TTCGAGGCTGGGAGCTTTTGGAGGGTCTGTGCGTCAGGAGAGTGTGCCCCTCACCCTCGCAGCGAGTCTCCTCATGGTCGCCTGGTACCAGGCGGTCC<br>CCTGAGGGTGCAGCTGATCGTCATGGGTGACTTCACGAGTCAGGACCTGTCACTCCATCCCTATAGTGAGTCGTATTACAA |
| *s100z* | CGAAATTAACCCTCACTAAAGGGAAGTCAGCTTGCTCACCACAACTGAGAAGGGGATTGCAGAGAATCCAGTGTCAAATGATGTGTGTGCCATGCAAG<br>ATGTGGGAGGATTTTGATGCACAAGGCAGCCAGGACAGTCATATCTGCAAGAAATGTCAGAGAACTCTGCCAAGATGCCATCCCAGCTGGAAGGTGCC<br>ATGGACTCGCTGATCCACATCTTCCACAATTATTCCGGCAAAGAGGGGGACAAGTACAAACTCAACAAGGGCGAAATGAGGGAACTTCTGCAAAGTGA<br>GCTGGGTAATTTTCTGGCGGCTCAGAAGGATCCATCTCTTGTGGACAATATCATGAAAGATCTTGATTCCAATAAAGACAATGAGGTGGATTTCAATG<br>AATTTGTTATTTTGGTTGCTGCTCTGACTGTTGCCTGCAATGACTTCTTTGAAGACCAACTCAGAAAAAAAGGAAAATCCCAATAGAACTGAAAATAT<br>CTTAAACAGATATCTTCATGAACACATGTTAAAAAATGATACACACCTGTCCCCTTTCCTTTGTAGAACCGCTGTAACTTCTATGTCTATGTATTTTG<br>TAAGTAAGAATGTATGATTCACTTTGAAGGTAATAAAATGCCTCAAGTTGCACAAACCCTATAGTGAGTCGTATTACAA |
| *trpc2* | CGAAATTAACCCTCACTAAAGGGATGGAGCATCTAACGAAAAAAAAGCCTGAGTACCTGAATCTGGAGGAACTCTGTGAGGACCTGGCCGTGGAATTG<br>CTCGGAATGTGCCGGAATCAGAGCGAAGTGAACGCTGTGCTGAACGACTGCGGTGATGAGAATGTGGAAGATGTGGACAACCAGACCTTTGAAGAAGG<br>GATTCCCAACCTGGGCAGGCTTCGCCTTGCTGTCAACTACAATCAGAAGAGGTTTGTGGCTCACCCCATCAGCCAGCAGGTACTCACCTCGATCTGGT<br>GCGGCAAGCTGACCTGGTGGCGAGGCAGCAAAACCATCTCAATCATCGGTGTCTCCTTGGGGATGTTCTTCACCATGCCCCTGATCTGTATCTGCTAC<br>TGGATTGCTCCCAAATCCAAGCTGGGACAGTTCGTTCGGACTCCAGTCATTAAATTCCTCCTTCACTCCTCTTCCTACATCTGGTTCCTGGTCTTGCT<br>CCTGGTGGAATCAATAGTCGCCCAACAATTCCGGGAACTGACCTCCTCCAGGCATGAACCGATCTACCTCAACTCTATACACATGATTTGGGTCGTGG<br>GCTTTTTCTGGTACGAATGCAAAGAGGTGTGGATCGAGGGTTTGCGCAGTTACCTACTGGACTGGTGGAATTTCCTGGATATTGTCATTCTGAGTATG<br>TACCTGGCTTCCTTCGCACTACGTATAGTGGTTTATCCCCTATAGTGAGTCGTATTACAA |
| *or3* | CGAAATTAACCCTCACTAAAGGGCACACACTCTCTTTGGGCTGCAAGCAGCTAACAATCCAGTAAAATGTATTTTTGATAAATAATTATCATAAAAGC<br>CCACTTCAAGGAAGAACATGAAAATAAGCATACGTGTTGAAGATCGCATGGAAGAAACTGGAAAAGTAAGAATTGCAGTTACAGACATCAACGCACTG<br>CGAAATGCATAAACCAAAATGACAAGATGCCCATGACAACTGATGAAGGAAAAGCTTTACACAATTTGTTCAGTCCAGTCAGAGAAGAACATTAAACA<br>GAATCATGAACAGCTCCAATGAGACTTCTGATTACATTTTAAATGTGGTGAAGACTTCAATTTATCTGACTGATTTTATTGTCACTAGTATGTGCAGT<br>CACATAATAATAAGTACTGTGCAGCAAGAGTTGGAGTTAAAGCGTGAGACCAGATACTTCCTCCTGTGCCAACATCTCATTTATGCTTCTATATATTT<br>TGCCCTGTGCACCCTAACAAATGGTTTGCGACTCTTCAGTATCAACTCACCCAGGATTCTTTGCTGGATCCTGTTTGCATTTCAAATAACATTCGCAC<br>AATGCATAATGCTAACTTTAACTCTAATGTCCTTGAACGCTTGTATAGCTATCTGTTGGCCTTTGCGATATACTTCACTTGTTGATTCAACAAATAAG<br>AAAATAACTGCAATAATGTGGGTGTTAGCGGTGCAGACCCTATAGTGAGTCGTATTACAA |
| *taar2* | CGAAATTAACCCTCACTAAAGGGATTTAATCTTCTTAAAGTGCTATTAACTGTAGTTGAAGAATTGCATTGGTTCATAAACACATCCAGAGAAATGGT<br>ATTAATAGTGTCGGACTATGCAGTTCAGATCCTGGGAACATCCTTCAAAGACAGTATCCATAATGATAACATCAAGAAAGATTAACCTCAAATGAAGG<br>AAATTGCAGTCTTAGTAAAGACCACACCAATCAATAACTCTGGAAGATCGGCTTTGTCATCTCACCGTGATCCAAGGAATTGGATTATCAATGTGAAA<br>AAGAATGAATTTAGCAGATCTTGAAAATTCAGAGGATGTGCAATATTGTTTTGAATTTATCAACACGTCCTGTCCCAGAGTCACCAGGTCAACAGCAG<br>CCAATGCAACGATGTATATTTTTATTACCATTTCAATACTCATTACTATATTTGGAAATCTGACGGTGATCATTTCTGTTTTACATTTCAAACAACTG<br>CAGACATCCACCAACTGTCTTCTCTTATCTTTGGCTGTTGTGGACTTTCTGGTGGGGTTTATTGTGTTGCCTTACAGTATGGTTAAGTCTGTAGAAAC<br>ATGCTGGTATTTTGGAGAAATGTTTTGTAAAGTTCACTCAATTATAGATATTGTGTTGACCGTAGTGTCAATTTATACTTTATGTTTTATTGCCATTG<br>ATCGTTACTATGCAATGTGTGACCCTTTGCTCCACCCCCCTATAGTGAGTCGTATTACAA |
| *v2rl1* | CGAAATTAACCCTCACTAAAGGGGGGTATGATGTGCGAGGCTTGGTAAAAGCTATTGCCTTGATCTACTCCATTGAGATGGTAAATAATTCAACTCTG<br>TTACCTGGGATCAAACTGGGGTACGAAATTTATGACACTTGCACGGAGCCTACAAAGGCACTGAAAGCAACTTTGAGGTTCCTTTCCGTATTCAACTC<br>GACAGACAACTGCGTCAAAGTTCAGTGCAATTATGCGGATTATCAGCCAGCAGTAAAGGCAGTTATTGGAGCTGGAACCTCAGAAGTTAGCATTGCCG |

| | |
|---|---|
| | TTGCAAGGTTGTTAAACATTCAACTTATTCCTCAGATCAGTTACTCAGCTTCTGCTAAAATCCTCAGTGACAAGGCTCGATTTCCTGCTTTTTTAAGA<br>ACTATTCCAAGTGACGACTATCAGACCAAAGCCATGGTCAAGATGGTTCAAAGATTCCAGTGGAACTGGGTGGGAACTATTGCAAGTGACGATGAATA<br>CAGCCGGTCTGGAATAGATAACTTCATTACACAAGCAGAAAGCCTTGGTATATGCATTGCGTTTCAAGAAGTGATTCCATTTTATTCATCTATTCAAG<br>TAACTAATTCCAGAATAAACGAAATAGCAAACATGGTCATCAACCGAACAAAAGTGAACGTCATTGTTATTTTTGGAAAAAGCACGCACGTCATTGAA<br>CTGTTTAAGATATTAGGCACTCACAACATAAGCAAAGCCCTATAGTGAGTCGTATTACAA |
| *v2rl4* | CGAAATTAACCCTCACTAAAGGGCCCAAGATCAAAAGAGGAAGCTGTTTTAATTTACTCTCAGACATCCCAATAAATACATCTAAGCAGAAGGAATAA<br>TGCTCGCTGTGAAGCCTGTCGATGTTTGGTGCTGTGTGTTTCTATGCTGGTTTCTATGCTGGACCGTGAGAGCCTGTGACACTGTAGGAGCCCGAAGC<br>CAAGGAGATATCAACATTGGTGGAATCTTCCCAATCCATCGTCAAGTGGAACATTTAGAAATTCGCTCTAAACCAGACCCATTGACATGTAAAGTGTT<br>TGATGTTCATAGATTTATCTGGCTACAGGCTATGATACATACAATTGAAATGACCAACAACTCCACGTTACTTCCTGGTATTAAACTTGGATATGAAA<br>TATATGATACCTGTACAGATGTCTCCACGGCTGTAACAGCCTCAATGAAGTTACTTTCTAAAGTGAATATCTCAGATAATTGTCTGGAGGTTCGATGC<br>AACTACACAGATTACCAACCGATTGTAAAAGTTGTTGTAGGTGAAATCTTTTCGGAGTTGTCGATTTCCACCGCAAGAATATTTAATGTAGCACTCAT<br>GCCTCAGATCAGTCCTGCATCATCTGCTGCAATCCTTAGCGACAAGCTGAGATTTCCATCATTGATACGCACAGTTCCAAGTGACATCCATCAGACAA<br>AGGCCTTGGCTCGATTTGTCAGCAACTCTCGTTGGAACCCTATAGTGAGTCGTATTACAA |
| *rh2* | CGAAATTAACCCTCACTAAAGGGATGAACGGCACAGAAGGAGAAAATTTTTATGTCCCGCTCTCCAACAAGACTGGAGTGGTTCGCAGTCCTTACGAG<br>TTCTCCCAGTATTACCTGGCTGTACCCTGGGAGTTCTCTGTCATCGCAGCTTACGTGTTTCTGCTGATGGTCATCGGCTTGCCCATCAATTGTTTGAC<br>TATGGCGGTCACTTTCAAGCACAAGAAACTAAGGTAGCCTCTCAATTACATTTAGTGAACCTGGCGGTGGCCAACCTGTTCATGATTCTCTTTGGGT<br>TTGTGGTCACATTTTACACAGCCTTGCATGGCTACTTTGTCTTTGGGCCCGTCGGCTGTGCGATTGAGGGCTTCTTCGCTACTCTACGTGGCGAAGTT<br>GCTCTTTGGTCCTTGGTGATCTTGGCTATAGAACGATACACCGTGGTCTGCAAACCGATGGGGAATTTCAGGTTTGGAGTGACTCATGCCTTTATTGG<br>AATCGACAGCACTTGGACCCTGGCTTTGGCCGCTGCCGGTCCTCCTCTGGTAGGTTGGTCCAGGTACATACCAGAAGGCTATCAGTGCTCATGTGGTC<br>CGGATTACTACACCATGAATCCACGTTACAACAATAAGTCTTATGTCATATATTTGTTTGTCTTTTACTTTCTGATTCCAGTTGCCCTCATATTCTTT<br>TCGTACGGAAGATTGATATGTAAAGTACAAGAGGCTGCCCTATAGTGAGTCGTATTACAA |
| *pinopsi n* | CGAAATTAACCCTCACTAAAGGGATGGGTTTTTCCACAGTTGCCATCATCACTCACAATACGACCATCGGTACTTTTGATGGCCCCCAGTGGCCACAT<br>GTTGCTCCAAAGAGTACCTACATGGCCGTGGCCACCCTCATGGGTACGGTGGTCATCTTGGCCTCCTTCATGAACGGGATGGTCATGCTCGTGTCCAT<br>CAAATACAAGAGGCTGCGCTCCCCGCTCAACTACATCCTGGTGAATCTGGCCATCGCTGACTTGTTGGTCACTTTCTTCGGCAGCACCATCAGTTTCT<br>CCAACAATGTCCATGGCTACTTCACTCTGGGGGAGTCGATGTGTCAGTTCGAAGGTTTCATGGTGTCCCTGACAGGTATTGTGGGACTGTGGTCCCTG<br>GCAATCTTGGCCTTCGAGCGGTACATTGTTATTTGCAAACCGATGGGCGATTTTCGATTCCAACAAAGGCACGCTGTGGGGGGGTGCCTCTTCACCTG<br>GATCTGGTCATTCATCTGGACTGTGCCACCTCTCATTGGCTGGTGCAGCTACGTACCTGAGGGTTTGCGACCCTATAGTGAGTCGTATTACAA |
| *cryg1* XM_ 038788722. 1 | CGAAATTAACCCTCACTAAAGGGGTGTTATATCAGTTACTCATCTAACTGTGAGCTCAAAATGGGAAAGATTATCTTTTACGAGGACAGGGACTTCCA<br>GGGTCGGCACTATGAGTGCAGCAGTGAGTGTGCTGATCTGTCCCCTTTCTTCAGCCGCTGTAACTCCATCCGCGTTGAGAGTGATTGGTGGGTGGTCT<br>ATGAGAGACCCAATTACATGGGATACCAGTATGTTCTGAGCAGGGGAGAATATACTGACTACCAGCGCTGGATGGGATTCAATGACAGCATTGGGTCA<br>TGTCGCTCTTACCCATATTACCGAGGTGGAAACTACAGAATGAAGATTTACGAGAGGCCCGACTTTGGAGGACAGATGATGGAATTCATGGATGACTG<br>TCCATCCGTCTATGATCGTTTCCGTTACCGTGACATCCACTCCTGCCATGTGATGGACGGTTACTGGATCTTCTATGAACATCCCAACTACAGAGGCC<br>GACAGTACTTCATGAGACCCGGTGAATACAGGAGATACAGTGACTGGGGTGGCTACAATTCAACTATCGGATCTTTCAGACGCATGAGGGATTTCTAG<br>ACTGTTCCTGACAAATATGGCTTTGTTTTTCTGAGACATGTGTCAGCAGAATAAATATACACAAATGTACCCTATAGTGAGTCGTATTACAA |
| *cryg2* XM_ 038788824. 1 | CGAAATTAACCCTCACTAAAGGGGTTCTCTTTTTGCTGACGTTCAGCATCAAATCATTTGAATAACAAAGGAACAGAGTAGAGAGCATTTTGTACAGC<br>ATATATATACCACACTGTGCAATGTTTTGCAATGTTAAAGCAGTTACTCATCTAACTGTGAGCTCAAAATGGCAAAGATCATCTTTTACGAGGACAGG<br>AACTTCCAGGGAAGGCATTATGAATGCAGTAATGACTGTGCTGACCTGTCCCCTTACTTCAGCCGCTGTAACTCCATGCGTGTCATGAGTGACTGGTG<br>GGTGATGTATGAGAGACCCAATTACATGGGATACCAGTATGTTCTGAGCAGGGGAGAATATCCTGACTACCAGCGCTGGATGGGATTCAATGACAACA<br>TTGGCTCATGTCGCACCTACCCATATTACCGAGGTGGAAACTACAGAATGAAGATTTATGAGAGGCCTGACTTTGGAGGACAGATGATGGAATTCATG<br>GATGACTGTCCATCTGTCTACGATCGTTTCCGTTACCGTGACATCCACTCCAGCCATGTGATGGACGGTTACTGGACCTTCTATGAACATCCCAACTA<br>CAGAGGTCGACAGTACTTCATGAGACCCGGTGAATACAGGAGATACAGTGACTGGGGCGGCTACAACTCAACTATCGGATCTTTCAGACGCATGAGGG<br>ATTTCTAGACTGTTCCTGACAAATATGATTTTGATTTTCTAACATATGTGTCAGCAGAATAAATATACATAATTTAACTGCTACCCTATAGTGAGTCG<br>TATTACAA |
| *trpv8.2* | CGAAATTAACCCTCACTAAAGGGATGTATATCACGGTGAACATGAACGATACTGCCAGGCATGGAAGTGAGTTTTTTAAAGCAGCTCACATAAATGAC<br>ACTGTCAGCATGAAGTCCTTGTTAACAAGTAAAGATTTTGATCCTGCCATTCGAGGGAATTTAGAAGAAACAATACTACATACAGCATTATTAAATGG<br>AAATAAGGAAATCGCAGAGTTCATTCTGGATATGATGCCATCACTGATCAATGAACCAATGACGTCTGATATGTACAGAGGTGAGACACCCATGCACA<br>TTGCTGTTCTGAAACAAGATGTGGAAATGGTTAAAGAACTTTTGAAGCGAGGTGCTGATGTGATGAATGCTCGTGCAACAGGATCCTGCTTTGTTCCC<br>GGAGAAGAATGTCAGTGTTACTATGGTGAATATTTGTTGTCGTTTGCAGCATGCATTGGCAATGAAGAGATTGTCAGAACTCTTGTTAAGCATTGTGC<br>ACCTTTAGAGGCACAGGATTCATTAGGTAACACAGTGCTTCATGTGTTAGTTCTACAACCAGAAAAAGAACAGGCATATTCCATGTATGATCTCATCA<br>CATCTCTTGTGTCTGAGAAACATCATCAATTTGTAGAAAACATTGTGAACAATGATGGCTACACTCCATTGAAGCTTGCAGCAGCTGAGGGTGACTTT<br>GTAATGTTCAACTACTTGGTGCAACCCTATAGTGAGTCGTATTACAA |
| *trpw1* | CGAAATTAACCCTCACTAAAGGGACGCTGGTTGTGACCATACCAGGGAACGAGGACGATGCAGAAGACAAGTGGAAGGACTATGTGCGAAAACGACGG<br>CTTCCACATAAACATATACCATTTAAAACGGATGCTGAGCTCGAGACCACAGATTCTGCCACCTATATTGGCTTGAAGAGGATTACAGATGCAGGACA<br>GCAAGTTGCGACATCGAATCAAGCAGTCCAGGGACAAGCAGGGACAAGCAGTCAGAACCAAAGTACTAGTACTTCCCCCACCACAGCCTTTCTTCAGT<br>ATTTCATCATAGCGTCAAAGAGAGGCAATGATAGTGAGATTGATTTGGAGCGCTTAAATACGATGCTGGAAAATGGAGCCGACATCAATGCTACTGAC<br>AGATACGGACAGGGTGTTTTGCACGGGGCAGCCCGAACATGGCACCCTGATGTTGCCAAGTTTCTGATCGAGAGAAATGCTGACGTGAACAAGCCTGA<br>TAACTACGGTGTGACCCCACTGCATGTGGCCGCAGCAAAAAATTATCCCGAAATGGTTGACTTTCTCCTGGATTGCGGCGCGGACGTTGAAGCAAAGA<br>CATCTGGAATGCTGCAGACACCAGTTCATTATGCGGCTAAGTATGATGCTAATAATTCCCTGCACTGTCTATTAAAACATGACGCCAACATGAAGGCT<br>CGAGACTACAAACAACGGACCCCTTTGCAACTGGCAACAGAGCTTGATCGATCTGAGTCAGCTCAATTGTTGCTGCAGTGTCAAGCAGATGCAGGAGT<br>TCATGATAATACAGCCCTATAGTGAGTCGTATTACAA |

#### Supplementary Table 6 sScyCan1.1 genome assembly summary statistics

| | |
|---|---|
| Scaffold L50 | 9 |
| Scaffold N50 | 198790641 |
| Scaffold L90 | 20 |
| Scaffold N90 | 98274498 |
| Scaffold len_max | 313568160 |
| Scaffold len_min | 995 |
| Scaffold len_mean | 6543240 |
| Scaffold len_median | 41167 |
| Scaffold len_std | 34693293 |
| Scaffold num_A | 1087862218 |
| Scaffold num_T | 1088048818 |
| Scaffold num_C | 990583604 |
| Scaffold num_G | 989674543 |
| Scaffold num_N | 64220747 |
| Scaffold num_bp | 4220389930 |
| Scaffold num_bp_not_N | 4156169183 |
| Scaffold num_seq | 645 |
| Scaffold GC content overall | 46.92 |
| Contig L50 | 616 |
| Contig N50 | 1855773 |
| Contig L90 | 2881 |
| Contig N90 | 254691 |
| Contig len_max | 13395245 |
| Contig len_min | 31 |
| Contig len_mean | 533457 |
| Contig len_median | 139317 |
| Contig len_std | 1028067 |
| Contig num_bp | 4156169183 |
| Contig num_seq | 7791 |
| Number of gaps | 7146 |

Supplementary Table 7 Length and population genetics statistics for *Scyliorhinus canicula* chromosomes.

$d_{XY}$ and π: absolute values for the whole genome or the whole chromosome; $F_{ST}$ value is the mean for all SNPs in the genome, or the chromosome.

| Chromosome | SeqID | Length (Mb) | $\pi_{Atl}$ | $\pi_{Med}$ | $d_{XY}$ | $F_{ST}$ |
|---|---|---|---|---|---|---|
| **(all genome)** | | | 0.001542286 | 0.0017562 | 0.001738406 | 0.044039844 |
| Chr1 | NC_052146.1 | 313.57 | 0.001201454 | 0.00137903 | 0.001371618 | 0.05084552 |
| Chr2 | NC_052147.1 | 289.50 | 0.001177534 | 0.001392878 | 0.001370372 | 0.054494517 |
| Chr3 | NC_052148.1 | 277.25 | 0.001399305 | 0.001551178 | 0.00155341 | 0.041699264 |
| Chr4 | NC_052149.1 | 244.32 | 0.001367927 | 0.001541541 | 0.001544124 | 0.052799902 |
| Chr5 | NC_052150.1 | 233.86 | 0.001448788 | 0.001649188 | 0.001639829 | 0.047476951 |
| Chr6 | NC_052151.1 | 225.97 | 0.001566239 | 0.001732001 | 0.001733123 | 0.045115562 |
| Chr7 | NC_052152.1 | 211.67 | 0.001336016 | 0.001484142 | 0.001476558 | 0.037521834 |
| Chr8 | NC_052153.1 | 199.96 | 0.001314683 | 0.00147339 | 0.00147393 | 0.041454005 |
| Chr9 | NC_052154.1 | 198.79 | 0.001470345 | 0.001640858 | 0.001639337 | 0.042336586 |
| Chr10 | NC_052155.1 | 191.88 | 0.001287479 | 0.001403209 | 0.001419262 | 0.044433329 |
| Chr11 | NC_052156.1 | 169.80 | 0.001450576 | 0.001598791 | 0.001606636 | 0.043272974 |
| Chr12 | NC_052157.1 | 165.19 | 0.00165505 | 0.001845942 | 0.001833786 | 0.040401988 |
| Chr13 | NC_052158.1 | 163.47 | 0.002153436 | 0.002507315 | 0.002440999 | 0.03483952 |
| Chr14 | NC_052159.1 | 160.85 | 0.001449808 | 0.001650224 | 0.001648518 | 0.050364088 |
| Chr15 | NC_052160.1 | 147.04 | 0.001404549 | 0.001563551 | 0.001568854 | 0.048335481 |
| Chr16 | NC_052161.1 | 144.97 | 0.00153299 | 0.001744982 | 0.001744633 | 0.051218034 |
| Chr17 | NC_052162.1 | 133.84 | 0.00212667 | 0.002847901 | 0.002604566 | 0.04342731 |
| Chr18 | NC_052163.1 | 131.92 | 0.001573778 | 0.0017472 | 0.001757485 | 0.050341696 |
| Chr19 | NC_052164.1 | 106.71 | 0.001447487 | 0.001639691 | 0.001630908 | 0.043732262 |
| Chr20 | NC_052165.1 | 98.27 | 0.001527127 | 0.001671391 | 0.001664758 | 0.028944284 |
| Chr21 | NC_052166.1 | 71.05 | 0.001753373 | 0.002017886 | 0.001991876 | 0.046768802 |
| Chr22 | NC_052167.1 | 30.79 | 0.002207089 | 0.002280123 | 0.002348847 | 0.039764935 |
| Chr23 | NC_052168.1 | 27.69 | 0.002446304 | 0.002668196 | 0.002645188 | 0.026448762 |
| Chr24 | NC_052169.1 | 27.32 | 0.001967814 | 0.002175314 | 0.002158729 | 0.034106878 |
| Chr25 | NC_052170.1 | 26.01 | 0.00218309 | 0.002338468 | 0.002383542 | 0.042868212 |
| Chr26 | NC_052171.1 | 24.91 | 0.002581717 | 0.002731119 | 0.002754937 | 0.029663898 |
| Chr27 | NC_052172.1 | 24.35 | 0.002508492 | 0.002722625 | 0.00273425 | 0.040446506 |
| Chr28 (X) | NC_052173.1 | 20.09 | 0.001143211 | 0.001380839 | 0.001384389 | 0.046883998 |
| Chr29 | NC_052174.1 | 16.13 | 0.003558252 | 0.003761555 | 0.003837485 | 0.034016374 |
| Chr30 | NC_052175.1 | 15.59 | 0.003456362 | 0.003566165 | 0.003675443 | 0.043657458 |
| Chr31 | NC_052176.1 | 12.11 | 0.003006646 | 0.003485276 | 0.003421928 | 0.048386055 |

#### Supplementary Table 8 Transposable element identification and counts.

| TE classification | coverage | count | proportion | Number_of_Distinct_Classifications |
|---|---|---|---|---|
| DNA | 253267242 | 323446 | 0,06001015 | 3521 |
| LINE | 1444957965 | 2491720 | 0,34237411 | 4123 |
| LTR | 492541379 | 536307 | 0,11670472 | 2482 |
| Other (Simple Repeat, Microsatellite, RNA) | 12245685 | 7888 | 0,00290154 | 330 |
| Penelope | 53256945 | 130137 | 0,01261891 | 241 |
| Rolling Circle | 8010116 | 18745 | 0,00189795 | 237 |
| SINE | 172129885 | 509312 | 0,04078514 | 338 |
| Unclassified | 439377534 | 1059955 | 0,10410787 | 1987 |
| SUM | 2875786751 | 5077510 | 0,6814004 | 13259 |

#### Supplementary Table 9 Small spotted catshark Hox genes.

The three Hox clusters (A, B, D) are located on chromosomes 5, 19 and 2 respectively.

| Gene ID | Gene symbol | NCBI gene symbol | NCBI description | Chromosome | Location |
|---|---|---|---|---|---|
| 119966064 | **hoxa1** | LOC119966064 | homeobox protein Hox-A1 | 5 | NC_052150.1 (78134776..78136249, complement) |
| 119966063 | **hoxa2** | hoxa2b | homeobox A2b | 5 | NC_052150.1 (78140155..78142273, complement) |
| 119966062 | **hoxa3** | LOC119966062 | homeobox protein Hox-A3 | 5 | NC_052150.1 (78145987..78190440, complement) |
| 119966066 | **hoxa4** | LOC119966066 | homeobox protein Hox-A4 | 5 | NC_052150.1 (78166954..78169027, complement) |
| 119966065 | **hoxa5** | LOC119966065 | homeobox protein Hox-A5 | 5 | NC_052150.1 (78179944..78183331, complement) |
| 119966067 | **hoxa6** | LOC119966067 | homeobox protein Hox-A6 | 5 | NC_052150.1 (78183448..78185283, complement) |
| 119966073 | **hoxa7** | LOC119966073 | homeobox protein Hox-A7 | 5 | NC_052150.1 (78191473..78194871, complement) |
| 119966072 | **hoxa9** | LOC119966072 | homeobox protein Hox-A9 | 5 | NC_052150.1 (78201022..78209928, complement) |
| 119966076 | **hoxa11** | LOC119966076 | homeobox protein Hox-A11-like | 5 | NC_052150.1 (78222714..78226938, complement) |
| 119965617 | **hoxa13** | LOC119965617 | homeobox protein Hox-A13-like | 5 | NC_052150.1 (78238035..78240447, complement) |
| 119954273 | **hoxb1** | hoxb1a | homeobox B1a | 19 | NC_052164.1 (5086117..5088982, complement) |
| 119954272 | **hoxb2** | LOC119954272 | homeobox protein Hox-A2-like | 19 | NC_052164.1 (5096966..5099248, complement) |
| 119954271 | **hoxb3** | LOC119954271 | homeobox protein Hox-B3a-like | 19 | NC_052164.1 (5103180..5109398, complement) |
| 119954277 | **hoxb4** | hoxb4a | homeobox B4a | 19 | NC_052164.1 (5124148..5126817, complement) |
| 119954275 | **hoxb5** | hoxb5a | homeobox B5a | 19 | NC_052164.1 (5138512..5141012, complement) |
| 119954274 | **hoxb6** | hoxb6a | homeobox B6a | 19 | NC_052164.1 (5142790..5145294, complement) |
| 119954279 | **hoxb7** | hoxb7a | homeobox B7a | 19 | NC_052164.1 (5150737..5153573, complement) |
| 119954278 | **hoxb8** | hoxb8a | homeobox B8a | 19 | NC_052164.1 (5154155..5156521, complement) |
| 119954372 | **hoxb9** | hoxb9a | homeobox B9a | 19 | NC_052164.1 (5164081..5166982, complement) |
| 119954371 | **hoxb10** | LOC119954371 | homeobox protein Hox-A10-like | 19 | NC_052164.1 (5174988..5181610, complement) |
| 119954291 | **hoxb13** | hoxb13a | homeobox B13a | 19 | NC_052164.1 (5267490..5269305, complement) |
| 119961873 | **hoxd1** | LOC119961873 | homeobox protein Hox-A1-like | 2 | NC_052147.1 (199582073..199584741, complement) |
| 119961872 | **hoxd2** | LOC119961872 | homeobox protein Hox-A2-like | 2 | NC_052147.1 (199589430..199592786, complement) |
| 119961875 | **hoxd3** | hoxd3a | homeobox D3a | 2 | NC_052147.1 (199595029..199633262, complement) |
| 119961879 | **hoxd4** | hoxd4a | homeobox D4a | 2 | NC_052147.1 (199615039..199623861, complement) |
| 119961877 | **hoxd5** | LOC119961877 | homeobox protein Hox-D5 | 2 | NC_052147.1 (199624916..199626729, complement) |
| 119961878 | **hoxd8** | LOC119961878 | homeobox protein Hox-D8 | 2 | NC_052147.1 (199636365..199638319, complement) |
| 119961876 | **hoxd9** | LOC119961876 | homeobox protein Hox-D9 | 2 | NC_052147.1 (199643590..199646176, complement) |
| 119961881 | **hoxd10** | LOC119961881 | homeobox protein Hox-D10-like | 2 | NC_052147.1 (199647178..199653254, complement) |
| 119962438 | **hoxd11** | hoxd11a | homeobox D11a | 2 | NC_052147.1 (199660937..199663041, complement) |
| 119962439 | **hoxd12** | hoxd12a | homeobox D12a | 2 | NC_052147.1 (199669103..199671289, complement) |
| 119962440 | **hoxd13** | LOC119962440 | homeobox protein Hox-D13 | 2 | NC_052147.1 (199676792..199678087, complement) |
| 119962441 | **hoxd14** | LOC119962441 | hematopoietically-expressed homeobox protein hhex-like | 2 | NC_052147.1 (199686837..199689125, complement) |

### SUPPLEMENTARY MATERIAL: Supplementary Figure List

#### Supplementary Figure 1

Variation of GC content in genomes, coding sequences or third codon in thirteen chondrichthyan species, in comparison to two osteichthyans species (the human *Homo sapiens* and the gar *Lepisosteus oculatus*) and a cyclostome, the marine lamprey *Petromyzon marinus*.

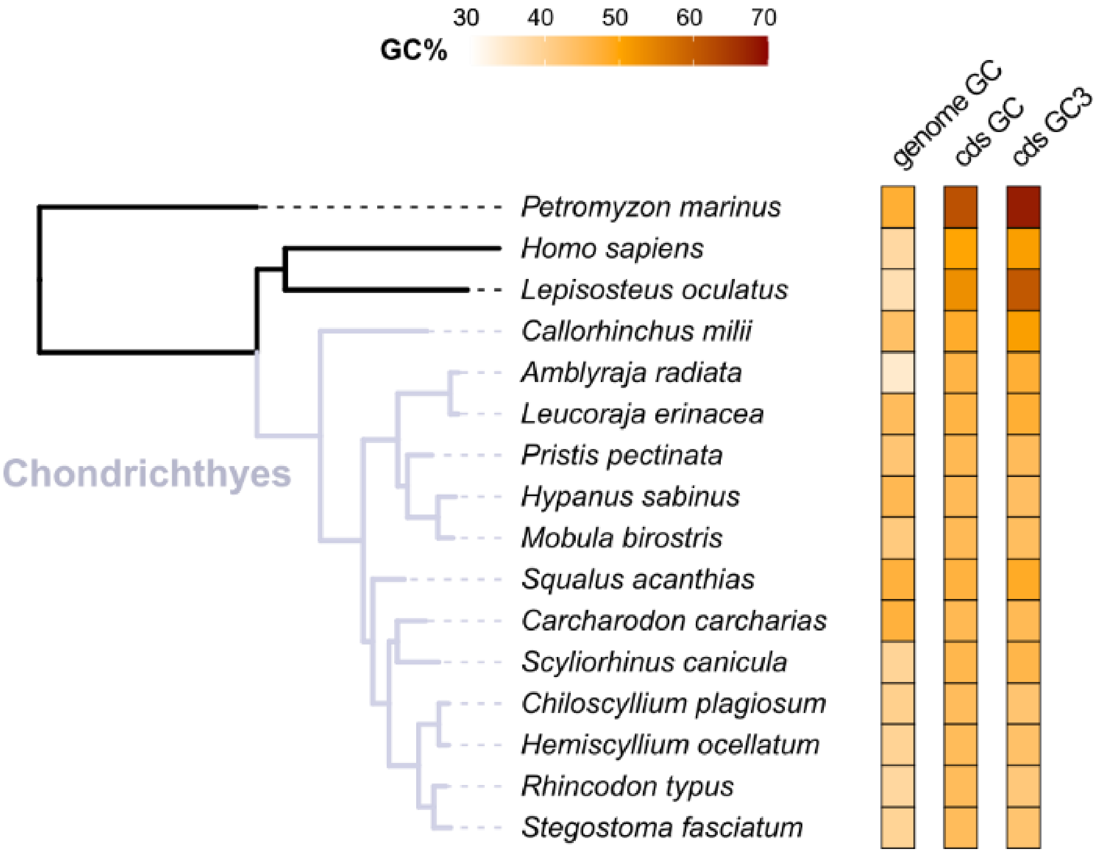

Supplementary Figure 2 Variation of GC content depending on chromosome size in thirteen chondrichthyan species, in comparison to two osteichthyans species (the human *Homo sapiens* and the gar *Lepisosteus oculatus*) and a cyclostome, the marine lamprey *Petromyzon marinus*.

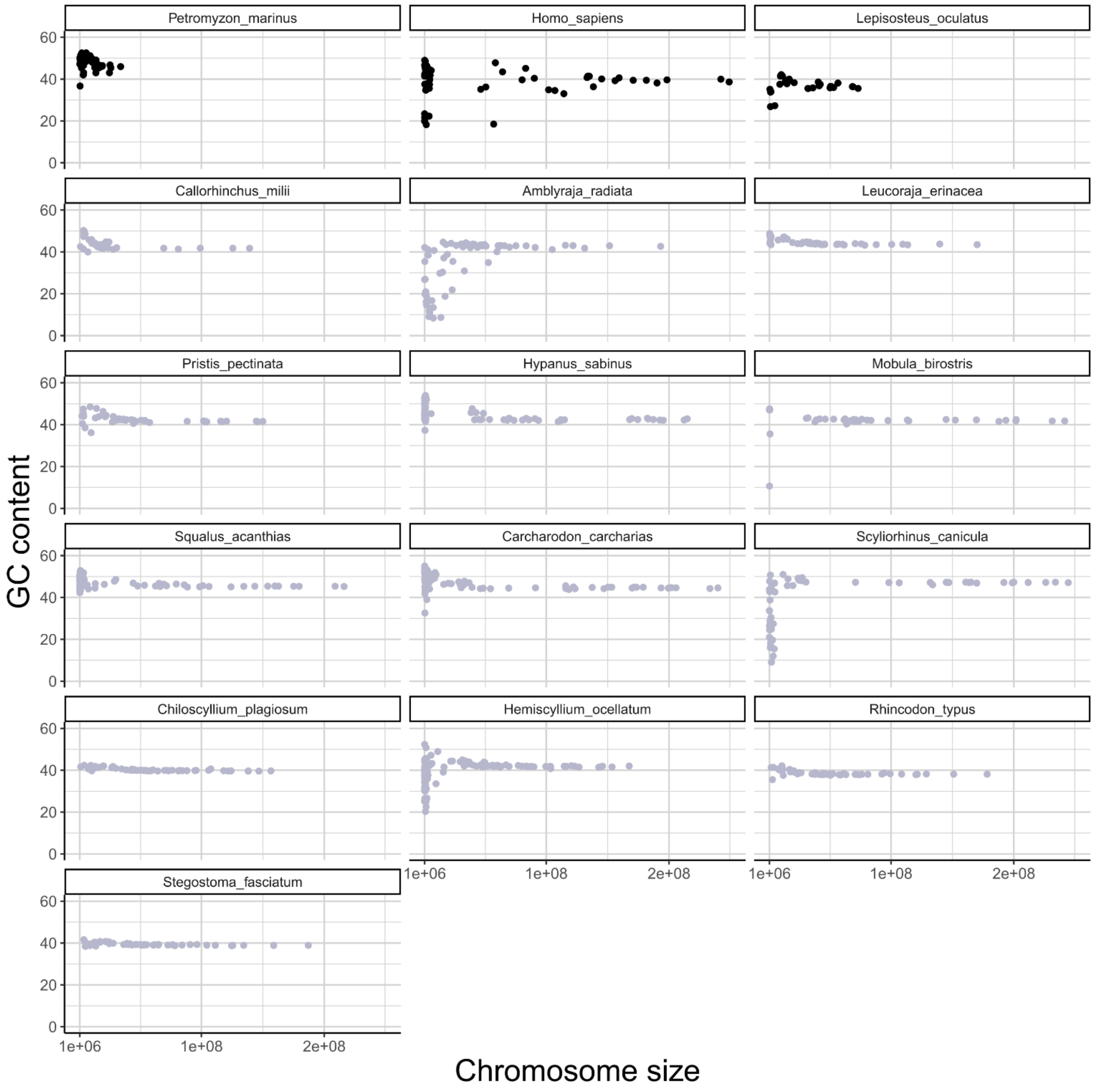

Supplementary Figure 3 BlobToolKit output for GC-coverage (A) and cumulative sequence plots (B).

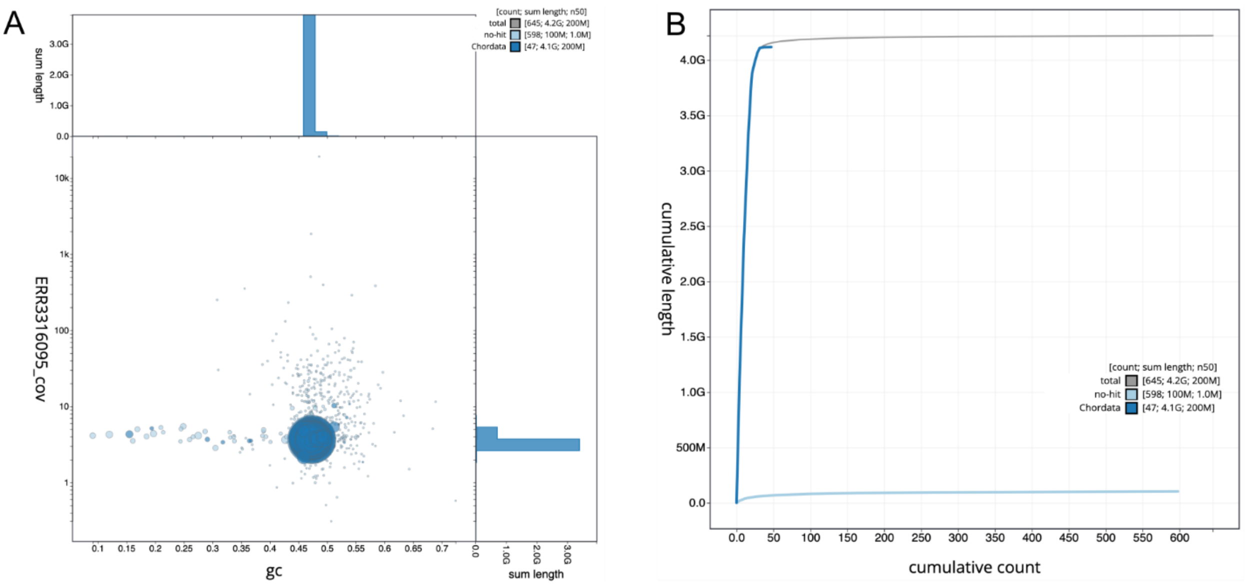

Supplementary Figure 4 Pearson correlation between statistics of the genomic landscape.

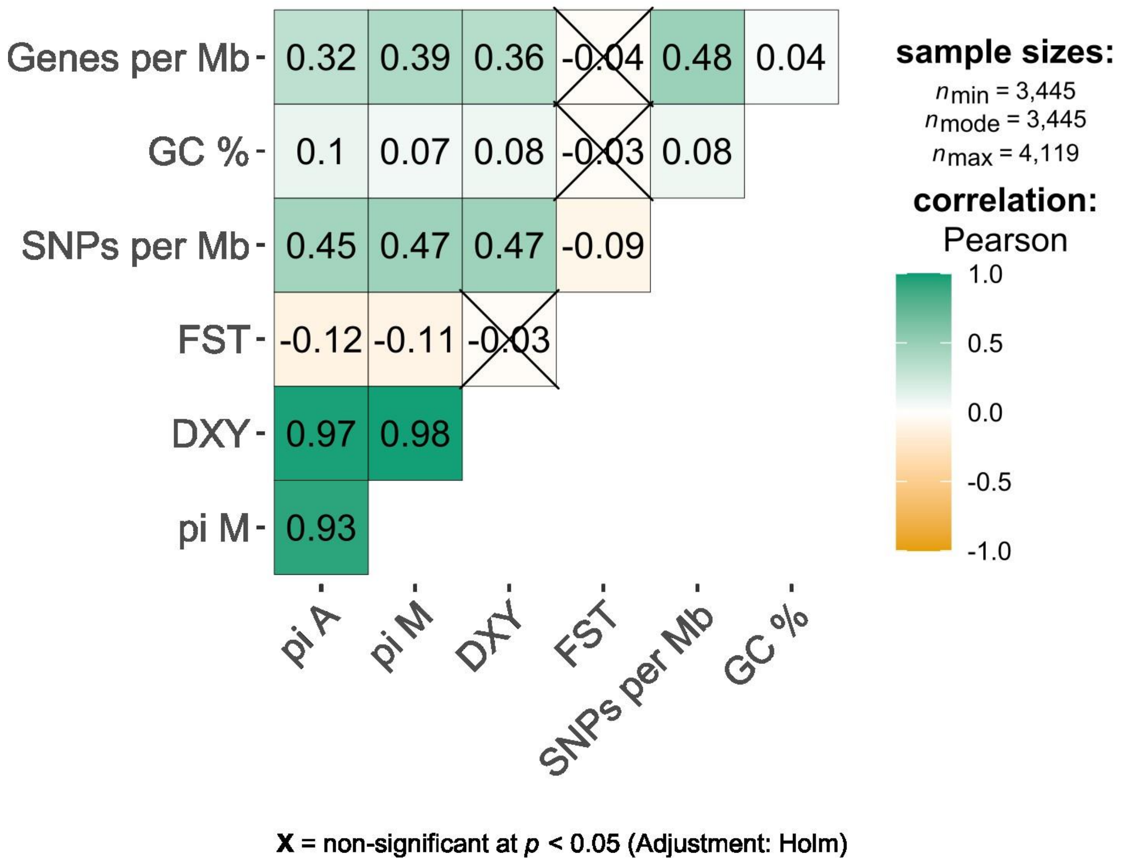

Supplementary Figure 5 Genomic organization of Hox gene clusters (A) and gene expression patterns in RNAseq data (B). A: Position of Hox genes and associated long non-coding RNAs along chromosomes 2, 5, and 19 in the small-spotted catshark. B: TPM values for each Hox gene and their associated long non-coding RNAs (highlighted blue) in a selection of tissues from the RNAseq data (only TPM values >5 shown (green), only in tissues where at least one TPM value in>10 (TPM value in orange). TPM values > 50 shown in red and >100 shown in brown. Gene names with asterisks belong to the gene cluster #1 in the autocorrelation analysis of embryonic expression (Supplementary Dataset 5).

A

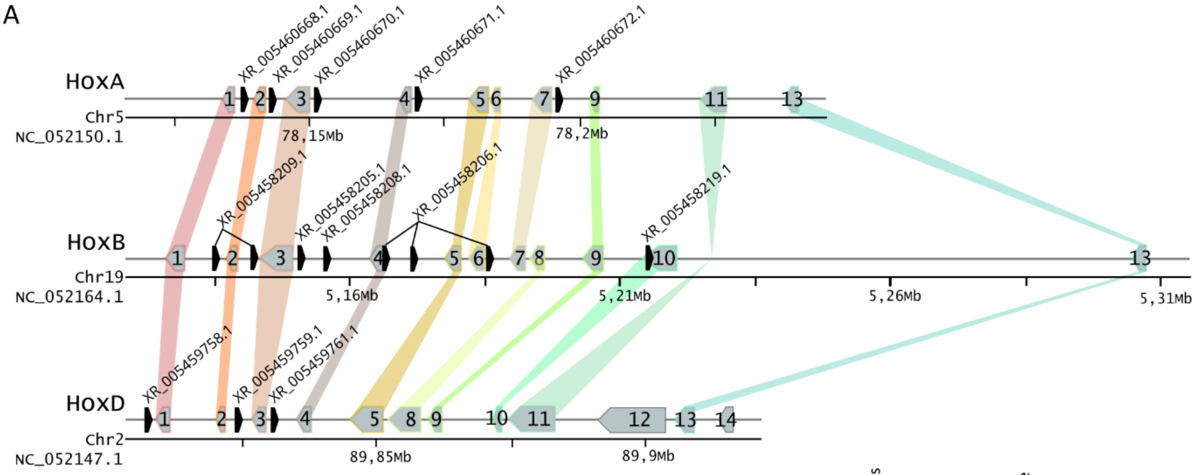

B

| | | E-St-12 | E-St-22 | E-St-24 | E-St-26 | E-St-30 | E-St-31 | Spinal chord | Kidney | Uterus | Seminal vesicle | Rectal gland | Esophagus | Gills | Denticles | Vertebrae | Muscle | Ovary | Testis |
|---|---|---|---|---|---|---|---|---|---|---|---|---|---|---|---|---|---|---|---|
| *hoxa1* * | XM_038797261.1 | 16,0 | 11,9 | 7,4 | | | | | | | 5,7 | | | | | | | | |
| | XR_005460668.1 | 37,2 | 46,3 | 28,4 | 32,0 | 25,6 | 25,5 | 20,1 | | 10,5 | 58,4 | 68,6 | 39,4 | 15,0 | 13,5 | 16,5 | 12,1 | | |
| *hoxa2* * | XM_038797260.1 | 20,9 | 70,9 | 51,1 | 55,3 | 24,8 | 12,5 | 15,5 | | | 19,0 | | 5,9 | 10,0 | | 9,0 | | | |
| * | XR_005460669.1 | 5,9 | 77,8 | 63,8 | 79,6 | 53,0 | 18,2 | 32,5 | | | 26,7 | 5,8 | 15,0 | 5,5 | | 12,2 | | | |
| *hoxa3* * | XM_038797256.1 | 7,9 | 57,6 | 62,8 | 56,2 | 46,3 | 22,1 | 21,6 | 18,1 | 13,3 | 31,6 | 16,2 | 6,2 | 18,5 | 12,7 | 5,7 | 5,4 | 7,1 | |
| * | XR_005460670.1 | 21,6 | 38,9 | 31,1 | 28,9 | 17,6 | 11,1 | 13,7 | | | 13,3 | | 16,5 | 19,4 | 9,8 | 19,9 | 15,9 | | |
| *hoxa4* * | XM_038797263.1 | | 18,8 | 19,8 | 16,6 | 8,9 | | 18,4 | | | 13,5 | 5,3 | 6,8 | | | | | | |
| * | XR_005460671.1 | | 6,3 | 7,3 | | | | 7,9 | | | | | 5,6 | | | | | | |
| *hoxa5* | XM_038797262.1 | | 19,7 | 33,8 | 19,5 | 17,0 | 10,3 | **155,4** | 34,0 | 5,4 | 34,1 | 12,3 | 5,9 | | 8,7 | | 6,5 | 5,1 | |
| *hoxa6* | XM_038797264.1 | | 8,2 | 15,6 | 8,6 | 7,1 | | 48,7 | 14,8 | | 10,6 | | | | | | | | |
| *hoxa7* * | XM_038797266.1 | | 18,4 | 17,0 | 16,1 | 12,6 | 8,9 | 13,7 | 14,2 | | | 6,4 | | | 12,9 | | | | |
| * | XR_005460672.1 | | 6,0 | | | | | | | | | | | | | | | | |
| *hoxa9* * | XM_038797265.1 | | 56,6 | 39,3 | 42,4 | 29,6 | 13,1 | | 41,5 | 16,1 | 9,2 | 11,6 | | | | | 6,1 | | |
| *hoxa11* | XM_038797267.1 | | 22,8 | 24,6 | 25,6 | 22,7 | 8,3 | | | 14,9 | 13,8 | 8,2 | | | | | | | |
| *hoxa13* | XM_038796400.1 | | | 8,3 | 10,2 | 13,7 | 6,4 | | | **223,0** | 5,5 | 44,3 | | | | | | | |
| *hoxb1* * | XM_038779368.1 | 51,1 | 38,4 | 21,1 | 17,9 | 5,2 | | 6,8 | | | | | | | | | | | |
| | XR_005458209.1 | | | | | | | | | | | | | | | | | | |
| *hoxb2* | XM_038779367.1 | | | | | | | | | | | | | | | | | | |
| *hoxb3* * | XM_038779366.1 | | 17,0 | 21,9 | 17,0 | 11,6 | 5,2 | 6,8 | 6,5 | | 10,9 | 6,2 | | | | 9,9 | | | |
| * | XR_005458205.1 | 17,7 | 71,1 | 53,4 | 46,2 | 27,6 | 14,7 | 63,9 | 8,7 | | 23,8 | 11,9 | 9,2 | 14,7 | | 55,7 | 5,4 | 17,8 | |
| * | XR_005458208.1 | 5,1 | 20,2 | 19,9 | 14,4 | 10,9 | 5,3 | 10,8 | | | 12,1 | | | | | 10,2 | | | |
| *hoxb4* * | XM_038779373.1 | 7,4 | 22,3 | 27,0 | 21,5 | 8,8 | 6,5 | 8,1 | | | 5,7 | | 9,6 | 6,2 | 5,4 | 13,0 | | | |
| * | XR_005458206.1 | 33,3 | 37,1 | 35,1 | 29,7 | 15,7 | 8,3 | 11,9 | 15,1 | | 12,3 | 46,8 | | | | 15,3 | | 11,1 | |
| *hoxb5* | XM_038779371.1 | | | | | | | | | | | | | | | | | | |
| *hoxb6* * | XM_038779369.1 | | | 6,4 | | | | 5,3 | | | | | | | | | | | |
| *hoxb7* * | XM_038779377.1 | | 37,9 | 36,9 | 34,0 | 27,6 | 20,6 | 60,3 | 31,1 | | 22,5 | 21,4 | 8,6 | | | | 9,3 | 9,4 | |
| *hoxb8* | XM_038779374.1 | | 34,6 | 33,9 | 31,4 | 32,6 | 18,5 | **141,1** | 50,8 | | 44,2 | 26,3 | | | | | 5,3 | 21,8 | |
| *hoxb9* * | XM_038779565.1 | | 62,2 | 57,9 | 51,5 | 26,2 | 13,3 | 24,3 | 83,7 | | 43,8 | **181,9** | | | | | 6,9 | 18,2 | |
| *hoxb10* * | XM_038779564.1 | | 15,6 | 24,1 | 16,6 | 13,7 | 6,1 | | 19,9 | | 11,3 | 62,2 | | | | | 16,2 | | |
| | XR_005458219.1 | | | 5,3 | | | | | 13,8 | | | 13,5 | | | | | 9,6 | | |
| *hoxb13* | XM_038779389.1 | | | 16,4 | 16,7 | 19,0 | | | | | 48,4 | **720,9** | | | | | | | |
| * | XR_005459758.1 | | | | | | | | | | | | | | | | | | 10,3 |
| *hoxd1* * | XM_038789329.1 | 68,4 | 29,1 | 16,4 | 11,6 | | | | | | | 6,0 | | | | | | | |
| *hoxd2* | XM_038789328.1 | 5,8 | 29,2 | 18,0 | 28,7 | 9,9 | | 6,7 | | | | | | | | | | | |
| | XR_005459759.1 | 10,3 | 8,5 | 5,3 | 6,5 | 8,1 | 5,7 | 6,1 | | | 5,0 | 10,7 | | | | 6,7 | | | |
| *hoxd3* * | XM_038789330.1 | | 39,4 | 27,1 | 28,4 | 13,4 | 6,9 | 8,8 | 21,3 | | 9,4 | 6,5 | | | | 7,4 | | | |
| * | XR_005459761.1 | | 27,7 | 18,9 | 18,8 | 5,7 | | | | | | | | | | 2,5 | | | |
| *hoxd4* * | XM_038789341.1 | 9,8 | 71,8 | 58,0 | 50,5 | 14,3 | 7,2 | 11,0 | 19,0 | | 10,1 | 9,0 | | | | 5,6 | | | |
| *hoxd5* * | XM_038789337.1 | | 18,2 | 26,2 | 18,2 | 12,2 | 10,2 | 20,2 | 43,3 | | 13,6 | | | | | 9,1 | 6,5 | | |
| *hoxd8* | XM_038789339.1 | | 13,7 | 19,0 | 17,8 | 17,3 | 8,3 | 9,1 | **126,0** | | 34,5 | | | | | | | 22,2 | |
| *hoxd9* * | XM_038789336.1 | | 21,2 | 21,0 | 21,8 | 18,1 | 9,6 | | 61,0 | | 15,0 | 8,1 | | | | | | 6,9 | |
| *hoxd10* | XM_038789343.1 | | 13,9 | 17,8 | 12,8 | 12,9 | 5,7 | | 17,0 | | | 8,0 | | | | | | | |
| *hoxd11* * | XM_038790391.1 | | 12,8 | 15,4 | 9,2 | 6,6 | | | | 37,4 | | 11,1 | | | | | | | |
| *hoxd12* | XM_038790392.1 | | 21,8 | 41,0 | 40,4 | 44,3 | 16,6 | | | **161,6** | | **123,7** | | | | | | | |
| *hoxd13* | XM_038790396.1 | | | 5,9 | 7,7 | 5,9 | | | | **208,1** | | **166,4** | | | | | | | |
| *hoxd14* | XM_038790397.1 | | 6,8 | | | | | | | 80,9 | | 47,8 | | | | | | | |

Supplementary Figure 6 Analysis of RNAseq data in adult organ samples.

A: Z-scores versus ln-transformed TPM values highlight highly specific together with highly expressed genes in each sampled organ; B: Pearson correlations for each pair of tissues and across selected genes (Z-score>1 and TPM>5 for at least one tissue)

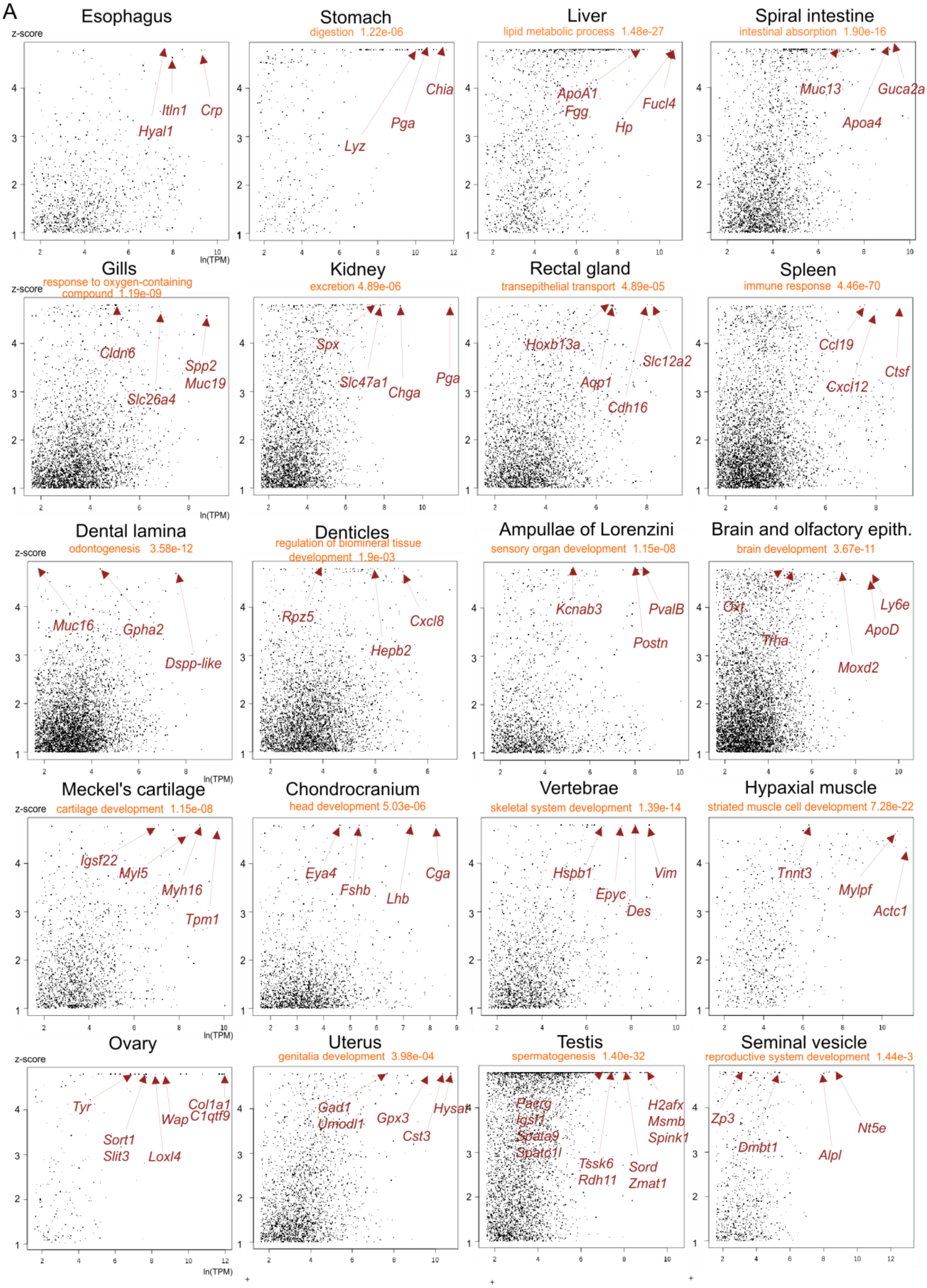

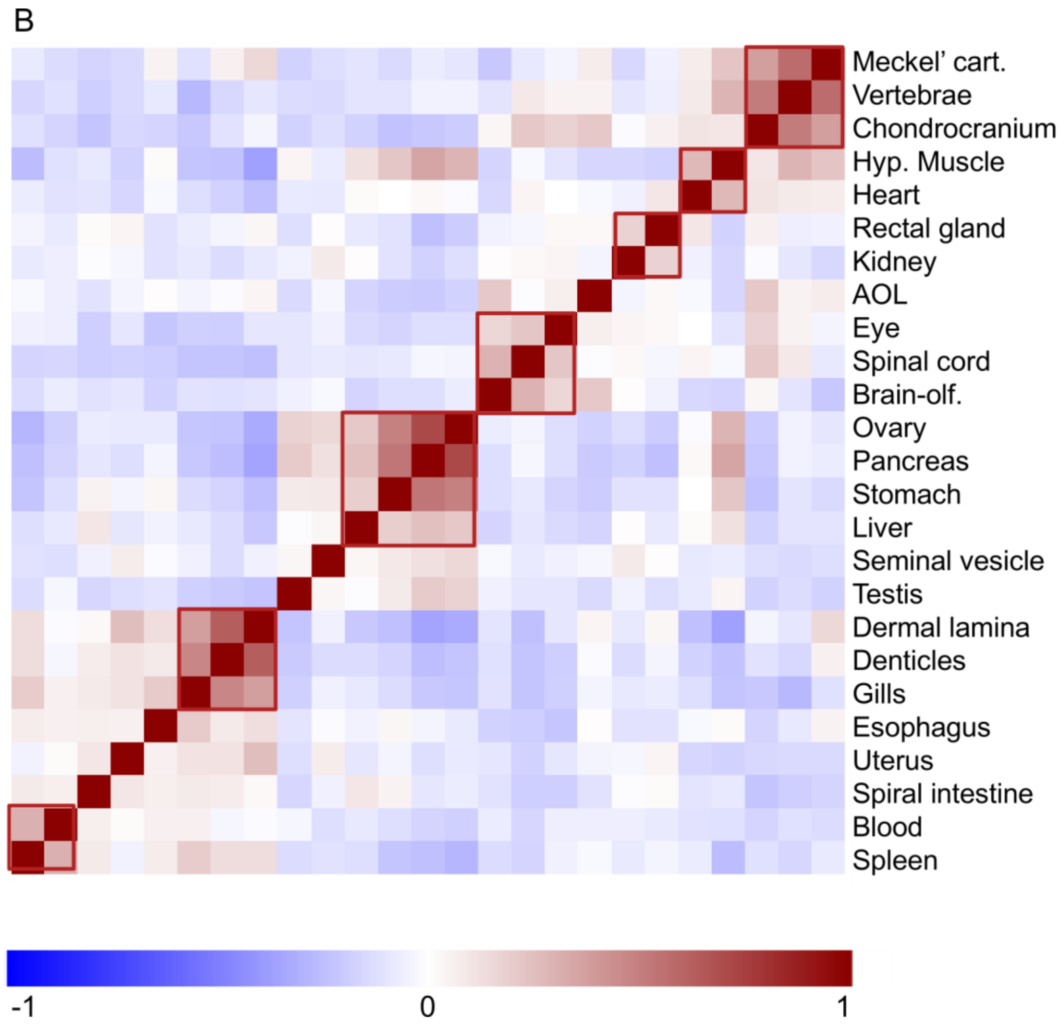
B
Meckel' cart.
Vertebrae
Chondrocranium
Hyp. Muscle
Heart
Rectal gland
Kidney
AOL
Eye
Spinal cord
Brain-olf.
Ovary
Pancreas
Stomach
Liver
Seminal vesicle
Testis
Dermal lamina
Denticles
Gills
Esophagus
Uterus
Spiral intestine
Blood
Spleen
-1
0
1

Supplementary Figure 7 Position of surface pores on the head of the small spotted catshark.

A. left lateral view; B. dorsal view; C. frontal view; D. ventral view. Black dots are lateral line pores; yellow dots are scales; purple and blue dots are pores of the ampullae of Lorenzini in the superficial ophthalmic (SO) and buccal (BUC) cluster respectively.

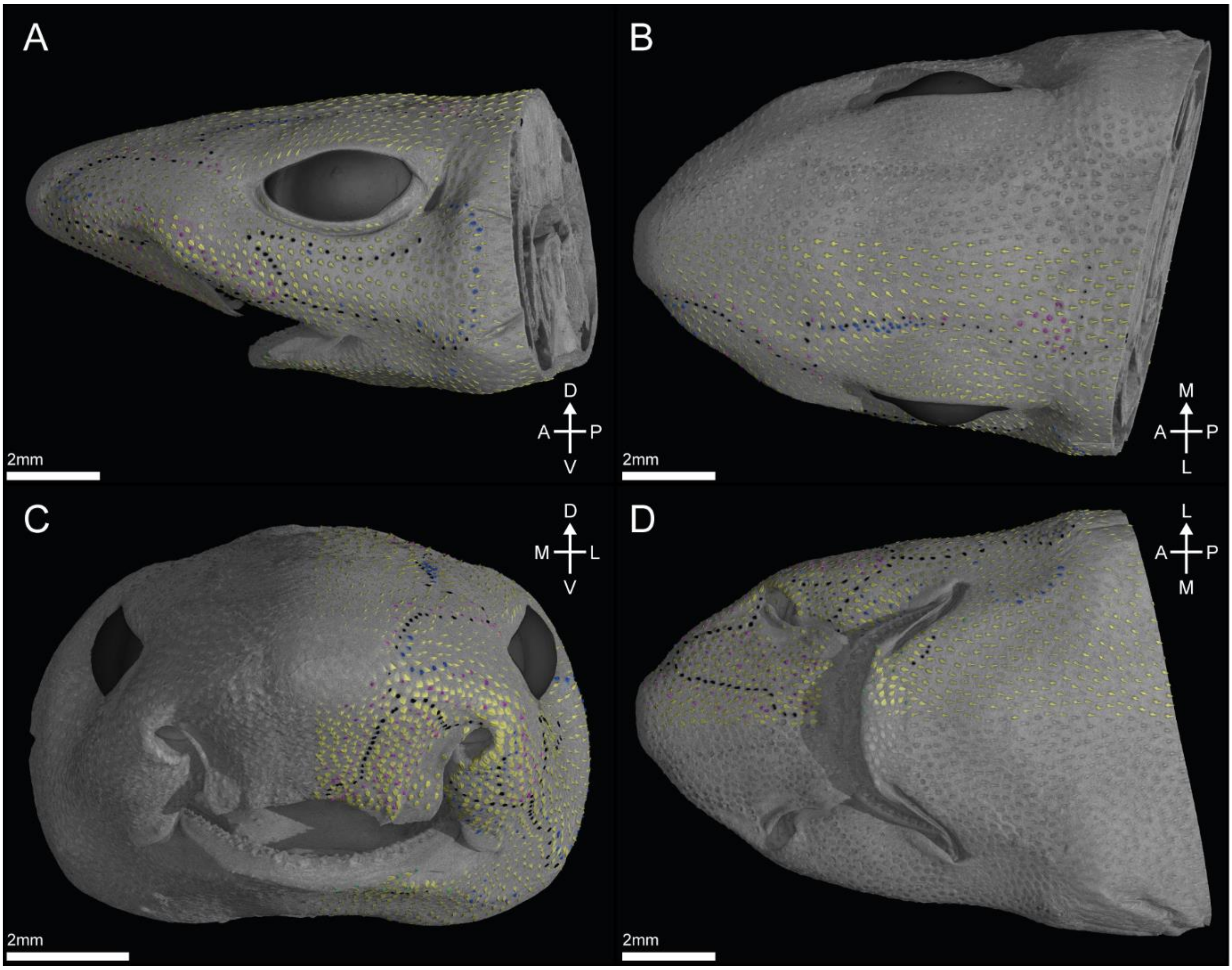

Supplementary Figure 8 Nervous connections between the olfactory epithelium and the olfactory bulb.

(A) Horizontal virtual thin section showing the lateral and medial olfactory nerves emanating posteriorly from the olfactory epithelium towards the braincase. (B) Dorsal view of the olfactory rosette in green, with the lateral and medial olfactory nerves innervating the lateral and medial olfactory lamellae, respectively. (C) Horizontal virtual thin section as in (A), but with the fibrous (blue) and cartilage-supported (yellow) braincase highlighted. (D) Anterior and slightly lateral view of the braincase showing the olfactory nerve foramen. The dotted line indicates the position of the horizontal sections in (A, C). c-BC: cartilage-supported braincase, f-BC: fibrous braincase, l-oln: lateral olfactory nerves, l-oln-fo: lateral olfactory nerve foramen, m-oln: medial olfactory nerves, m-oln-fo: medial olfactory nerve foramen, OE: olfactory epithelium, OB: olfactory bulb, tel: telencephalon, tng: terminal nerve ganglion. Orientation: A - anterior, P - posterior, M - medial, L - lateral, D - dorsal, V - ventral.

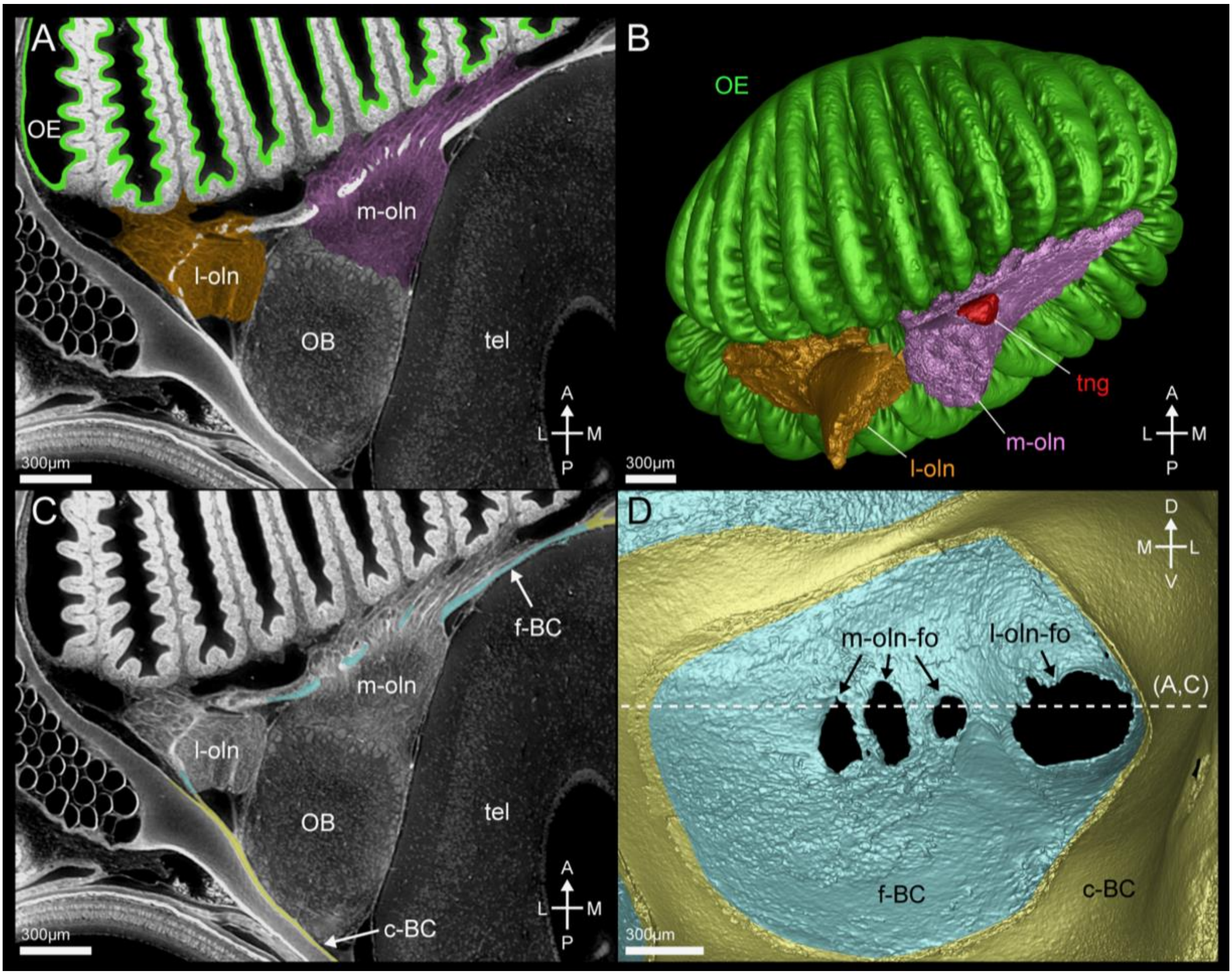

#### Supplementary Figure 9 Selected views of Supplementary dataset 9 with legends.

Horizontal virtual sections through the left olfactory bulb with anterior to the bottom, lateral to the right. (A) Section plane showing the passage of the lateral olfactory nerves through the braincase foramen to the lateral olfactory bulb. (B) Section plane showing the passage of the median olfactory nerves through the braincase foramen to the median olfactory bulb.

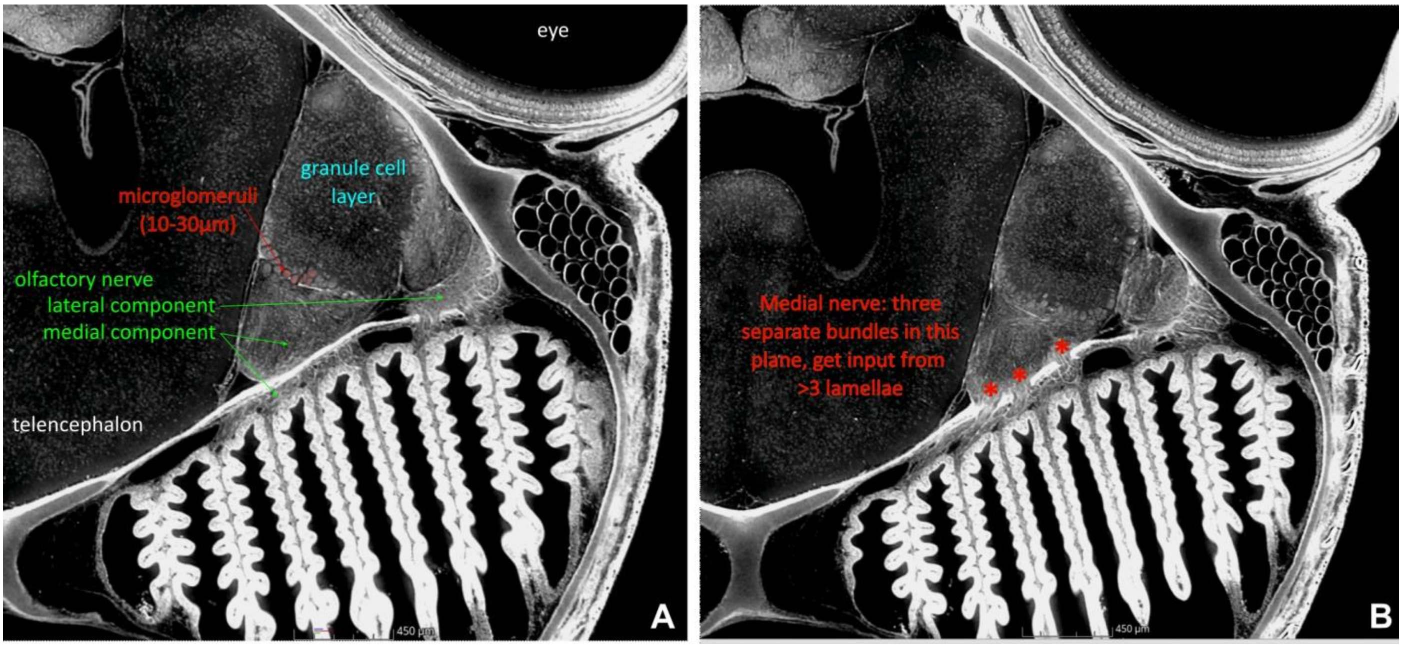

Supplementary Figure 10 Vertebrate Moxd gene tree inference by ML, rooted by amphioxus sequences.
The alignment was 753 positions, the best fit model was VT+R4. Branch supports: SH-aLRT/ultrafast bootstraps (percentages); see main text for species names. The catshark sequences are highlighted in color, gnathostome orthology groups are identified by the position of bullets.

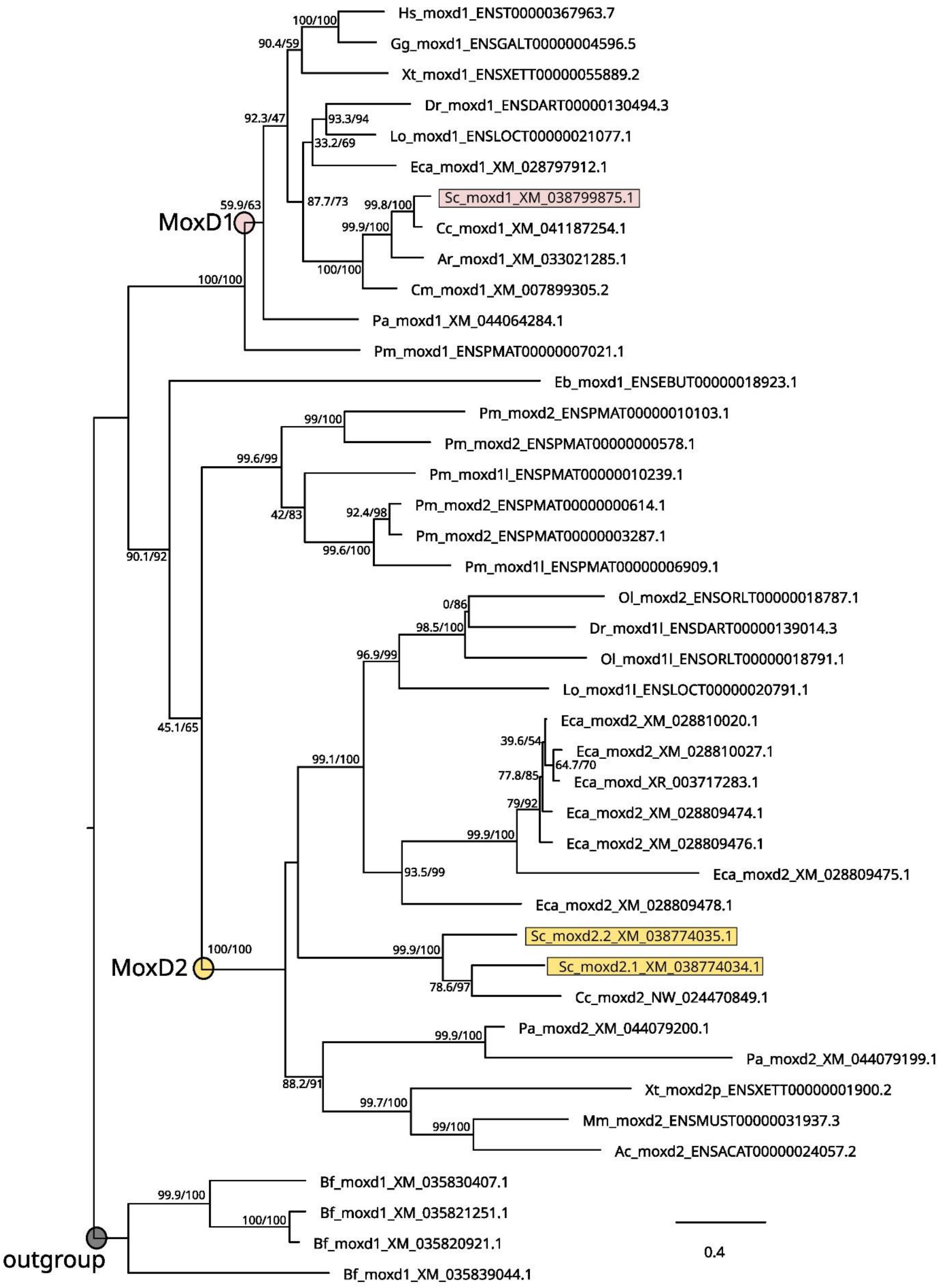

#### Supplementary Figure 11 Gene expression pattern for *moxd2.2*.

ecm: extracellular matrix of the basal conjunctive tissue; lu: lumen of the olfactory rosette; dotted line: basal lamina between the sensory epithelium and the underlying conjunctive tissue; open arrowhead: putative sensory cell bodies; black arrowhead: putative supporting cells. Scales in microns.

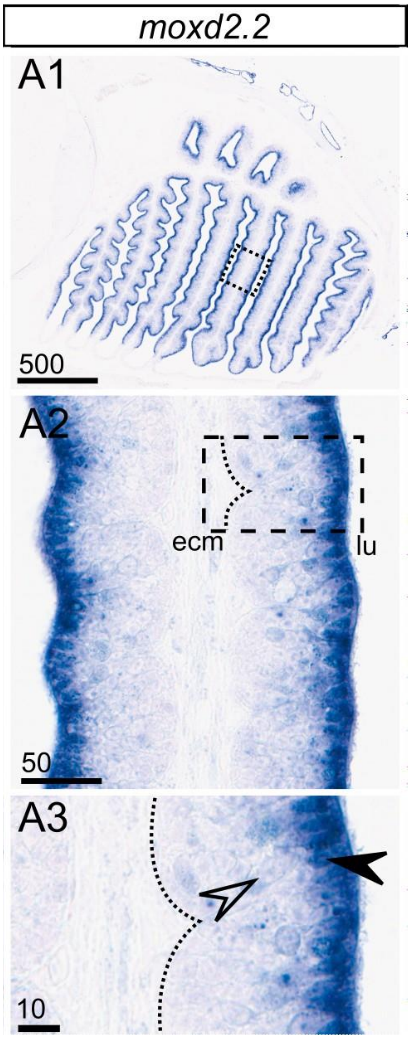

Supplementary Figure 12 Gnathostome s100z gene tree inference by ML, rooted by gnathostome s100a and lamprey sequences.

The alignment was 104 positions, the best fit model was Q.insect+G4. Branch supports: SH-aLRT/ultrafast bootstraps (percentages); see main text for species names. The catshark sequences are highlighted in color, gnathostome orthology groups are identified by the position of bullets.

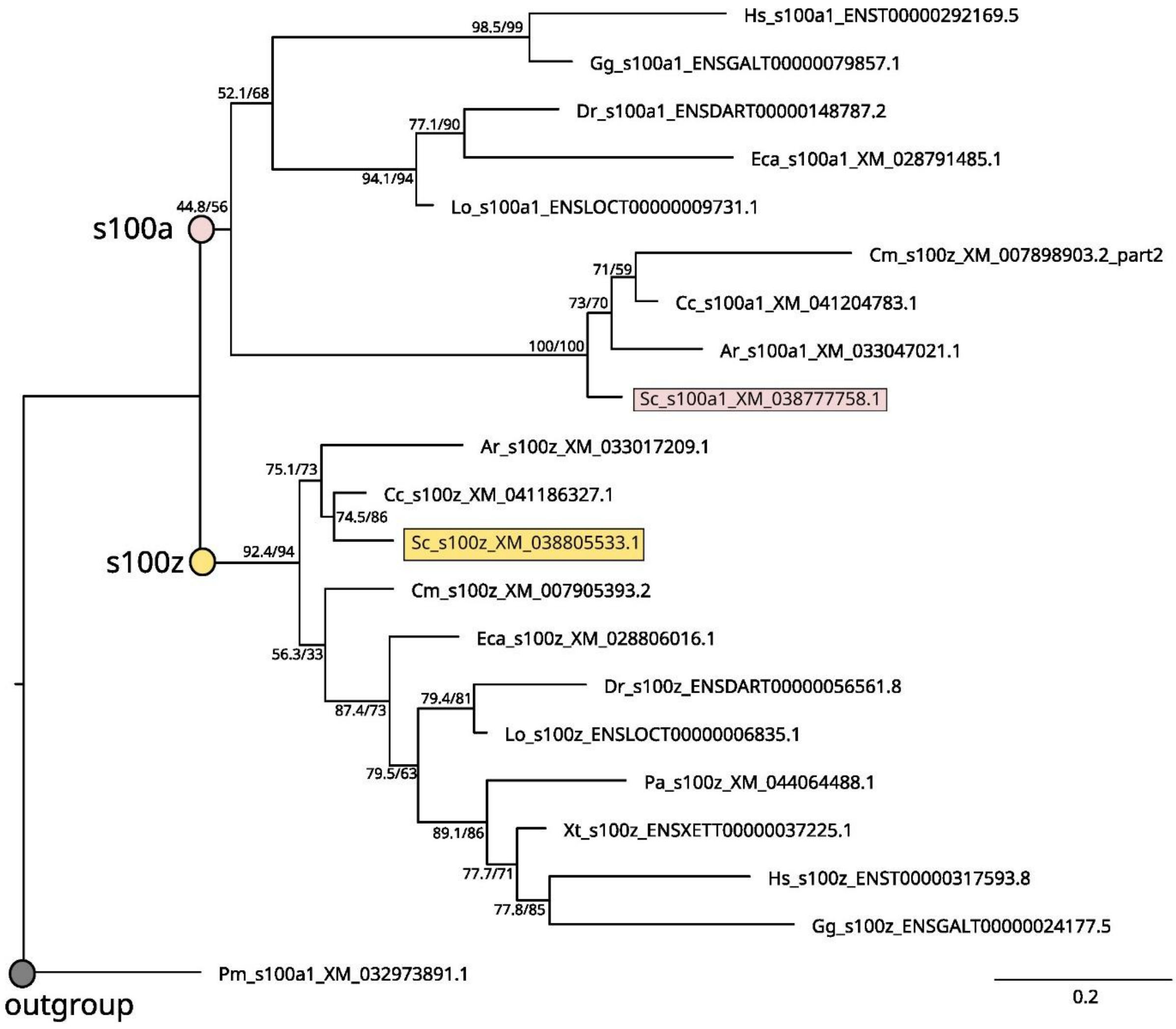

Supplementary Figure 13 Levels of expression (TPM values) of chemo-sensory receptor families in the reference RNAseq. Values are highlighted depending on the associated Z-score value (scale on the right side of the panel).

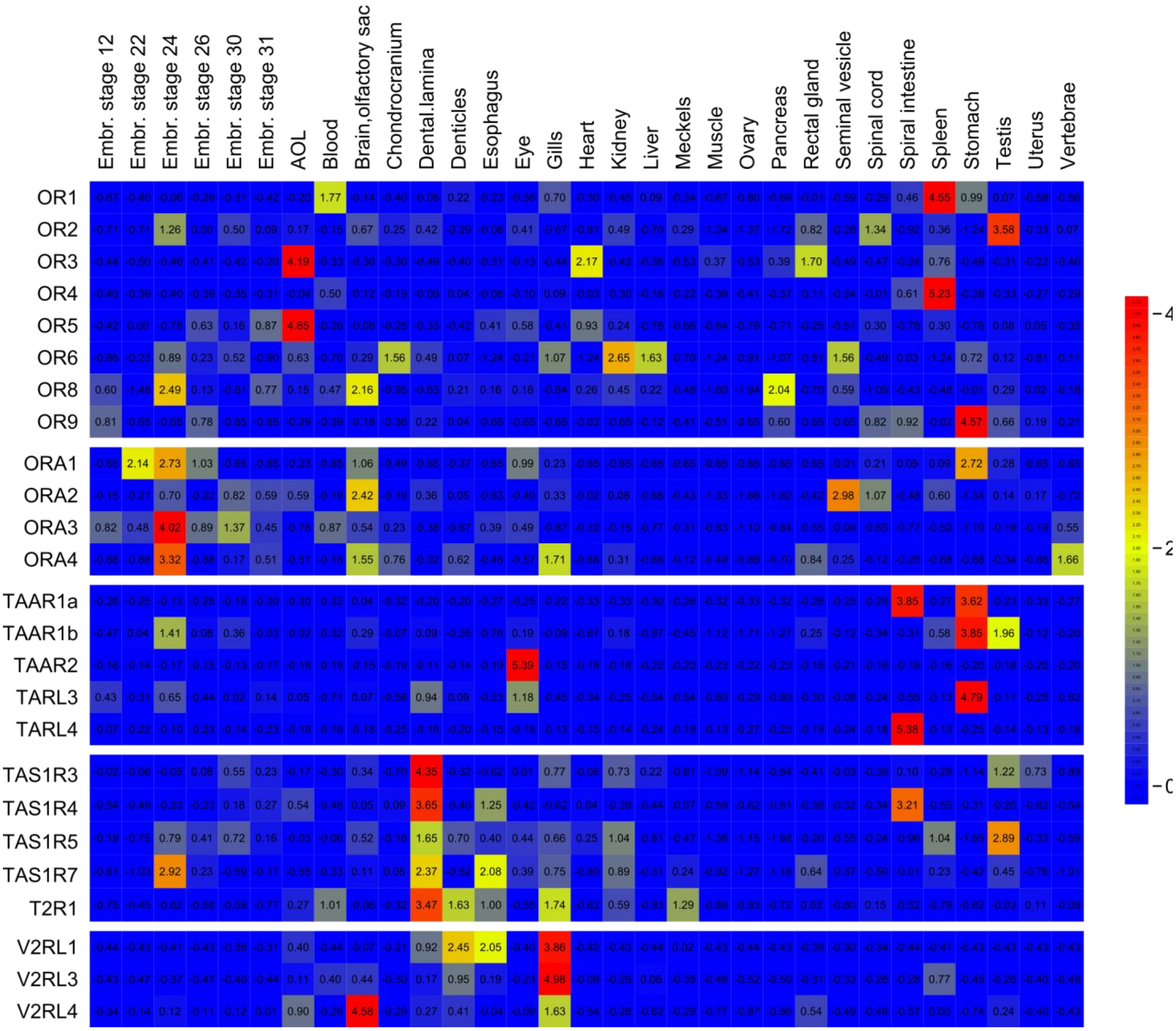

Supplementary Figure 14 Phylogenetic relationship and level of expression for V2R genes.

Expression levels are TPM values in the reference RNAseq data, highlighted depending on the associated Z-score value (scale on the right side of the panel). Phylogenetic relationships following Syed et al. 2023.

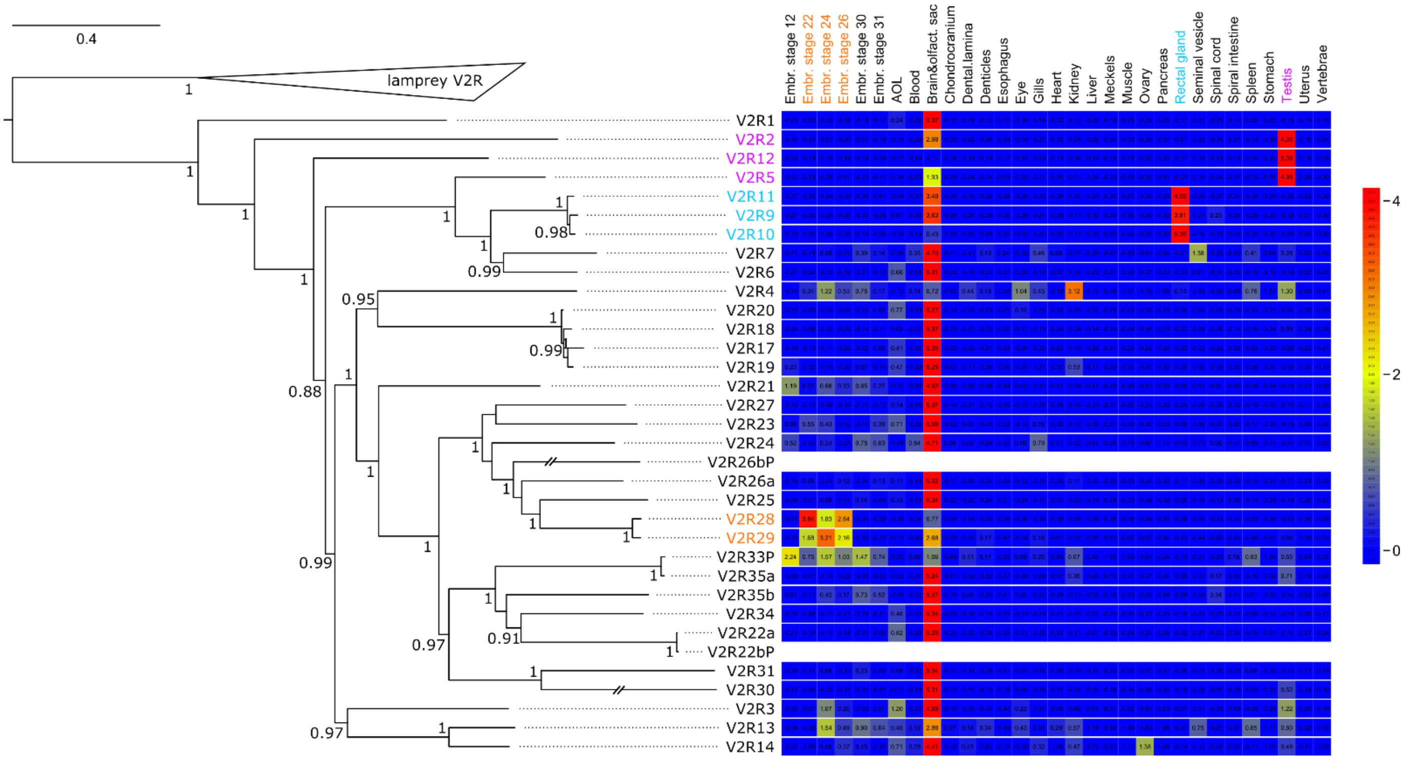

Supplementary Figure 15 Gene expression patterns for *v2rl4* in transverse sections of a juvenile catshark (except in **B, C**: stage 31 embryo).

Expression in the olfactory epithelium (**A**) and non-olfactory sites: undifferentiated retina (**B**, gc: ganglion cell progenitors); posterior brain (**C**, v: ventricle of the mesencephalon); taste bud (**D**); epiderm and lateral line (LL; **E**), sb: scale bud; gills (**F**) and muscle tissue (**G**). Black arrowheads point to scattered epithelial cells positive for gene expression; open arrowhead points to muscle (**G**) cells. Scales in microns.

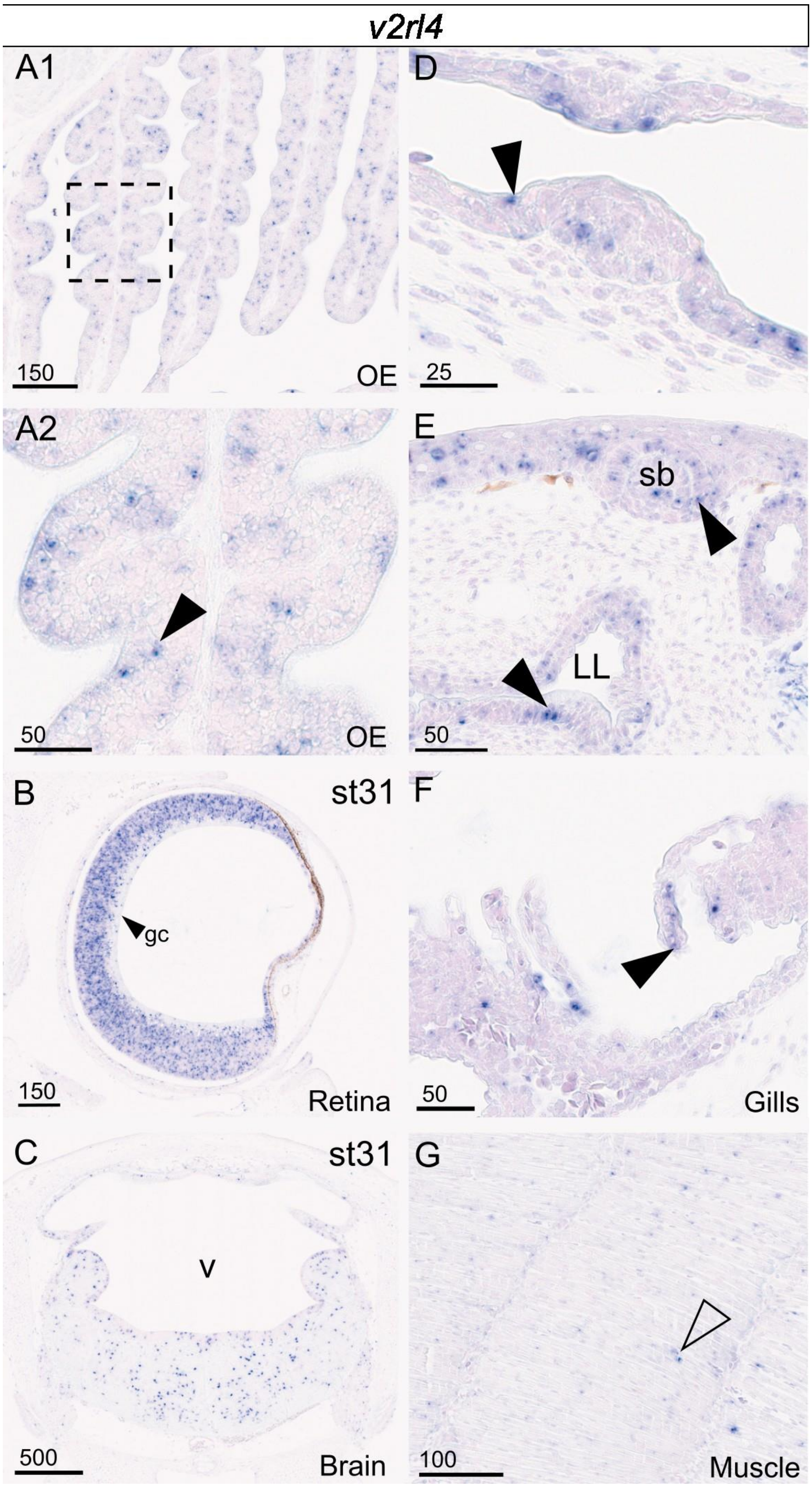

Supplementary Figure 16 Jawed vertebrate visual opsin gene tree inference by ML, rooted by parapinopsin/parietopsin sequences.

The alignment was 523 positions, the best fit model was LG+F+R5. Branch supports: SH-aLRT/ultrafast bootstraps (percentage); see main text for species names. The catshark sequences are highlighted in color, gnathostome orthology groups are identified by the position of bullets.

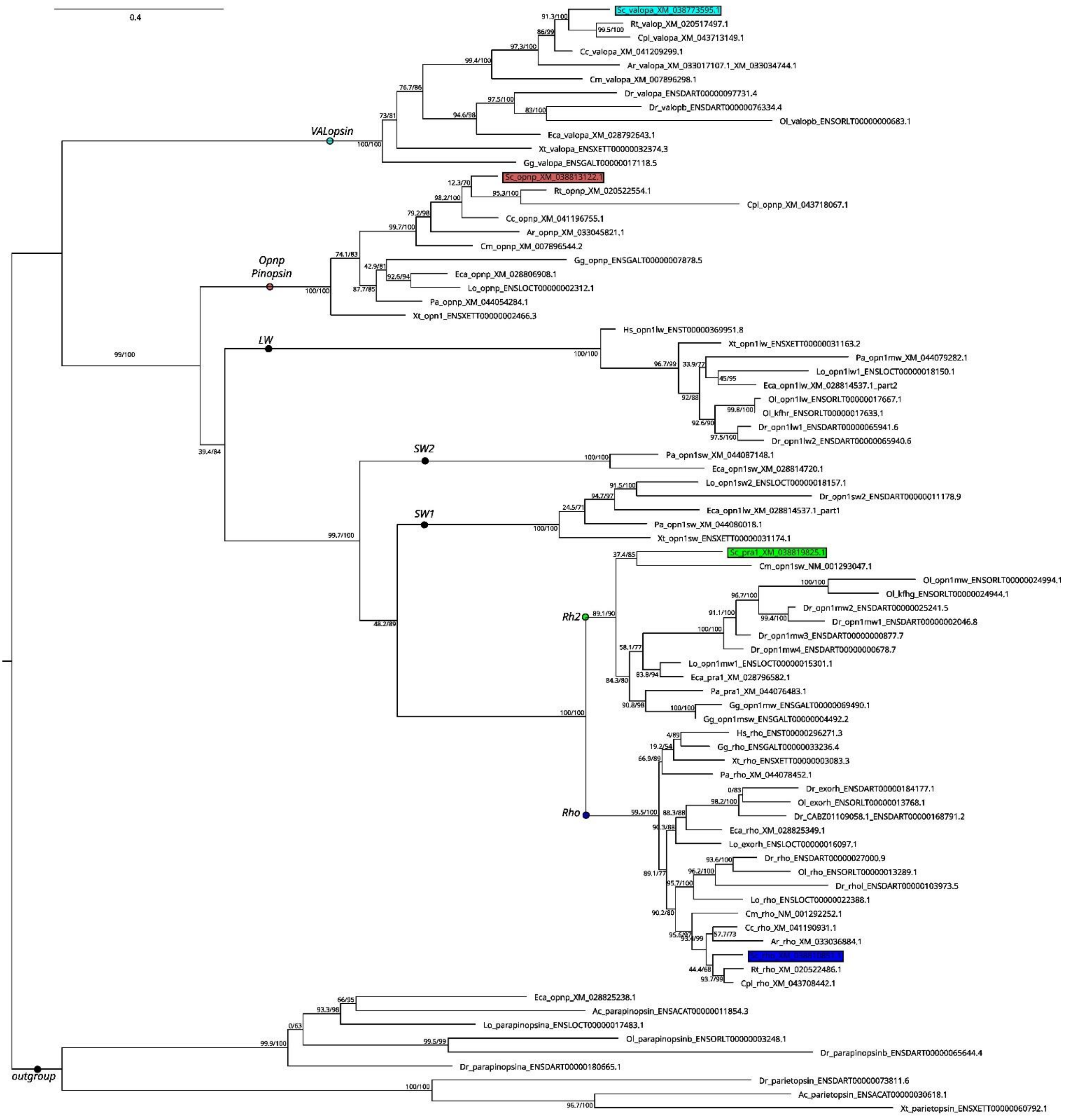

Supplementary Figure 17 Levels of expression (TPM values) of opsin gene families in the reference RNAseq. Values are highlighted depending on the associated Z-score value (scale on the bottom of the panel).

| | Embr. stage 12 | Embr. stage 22 | Embr. stage 24 | Embr. stage 26 | Embr. stage 30 | Embr. stage 31 | AOL | Blood | Brain, olfactory sac | Chondrocranium | Dental lamina | Denticles | Esophagus |
|---|---|---|---|---|---|---|---|---|---|---|---|---|---|
| *opn3* | 1.53 | 1.56 | 1.28 | 1.25 | 0.66 | 0.76 | -0.38 | -0.59 | 1.54 | -0.12 | -0.55 | -0.74 | -0.60 |
| *opn4* | -0.21 | -0.18 | -0.18 | -0.19 | -0.19 | -0.18 | -0.19 | -0.21 | -0.17 | -0.18 | -0.20 | 0.23 | -0.19 |
| *opn5L1B* | -0.76 | 0.05 | 0.29 | -0.18 | -0.12 | 0.37 | 0.25 | -0.02 | 0.45 | -0.04 | 0.09 | 2.78 | -0.24 |
| *pinopsin* | -0.61 | 0.12 | 0.02 | -0.16 | 0.84 | 0.79 | 2.88 | -0.07 | 1.25 | 0.31 | 1.18 | 0.09 | -0.67 |
| *va-opsin* | -0.64 | 1.79 | 0.56 | 0.09 | 1.26 | 0.82 | 1.32 | -0.52 | 0.18 | -0.01 | -0.47 | -0.13 | -0.13 |
| *rgr1* | -0.40 | -0.01 | 1.28 | 0.11 | 0.52 | 0.01 | 0.17 | -0.16 | 0.28 | 0.09 | 0.63 | -0.06 | -0.56 |
| *rh2* | 0.15 | 0.24 | 1.96 | -0.54 | 3.94 | -0.56 | -0.80 | -0.82 | -0.25 | 0.20 | 0.19 | -0.32 | -0.23 |
| *rho* | -0.18 | -0.18 | -0.18 | -0.18 | -0.18 | -0.18 | -0.18 | -0.18 | -0.18 | -0.18 | -0.18 | -0.18 | -0.18 |
| *rrh* | -0.30 | 0.16 | 0.88 | 0.25 | 0.94 | 0.44 | -0.19 | 0.02 | 0.89 | -0.16 | 0.20 | -0.10 | -0.30 |
| *tmt1* | -0.20 | -0.17 | 0.08 | 0.11 | -0.08 | -0.19 | -0.38 | -0.17 | 0.17 | 0.05 | -0.10 | -0.32 | -0.26 |

| | Eye | Gills | Heart | Kidney | Liver | Meckels | Muscle | Ovary | Pancreas | Rectal gland | Seminal vesicle | Spinal cord | Spiral intestine | Spleen | Stomach | Testis | Uterus | Vertebrae |
|---|---|---|---|---|---|---|---|---|---|---|---|---|---|---|---|---|---|---|
| *opn3* | -0.29 | -0.69 | -0.78 | -0.43 | -0.36 | -0.68 | -0.77 | -0.79 | -0.79 | 0.34 | -0.62 | -0.21 | -0.71 | -0.60 | -0.77 | 3.29 | -0.35 | -0.41 |
| *opn4* | 5.37 | -0.20 | -0.21 | -0.20 | -0.21 | -0.15 | -0.22 | -0.23 | -0.22 | -0.20 | -0.21 | -0.14 | -0.22 | -0.20 | -0.23 | -0.07 | -0.21 | -0.18 |
| *opn5L1B* | 2.18 | -0.06 | -0.48 | -0.21 | -0.14 | -0.50 | -1.09 | -1.49 | -1.44 | -0.24 | -0.25 | 1.29 | -0.31 | 0.08 | -1.28 | 2.41 | -0.60 | -0.79 |
| *pinopsin* | 0.89 | 0.08 | -1.19 | 0.59 | -1.10 | -0.24 | 0.13 | -1.93 | -1.96 | 0.04 | -0.52 | -0.17 | -1.02 | 0.64 | -1.52 | 0.41 | 0.01 | 0.88 |
| *va-opsin* | 4.04 | -0.53 | -0.75 | -0.34 | -0.59 | -0.34 | -0.66 | -0.75 | -0.72 | -0.64 | -0.59 | 0.11 | -0.66 | -0.38 | -0.75 | 0.21 | -0.38 | -0.40 |
| *rgr1* | 3.69 | 0.17 | -0.79 | 0.32 | -0.92 | -0.37 | -1.21 | -1.53 | -1.51 | -0.18 | 0.01 | -0.10 | -0.67 | 0.55 | -1.17 | 1.81 | 0.31 | -0.32 |
| *rh2* | -0.12 | 0.40 | -0.59 | -0.13 | -0.32 | 1.56 | -0.37 | -0.78 | -0.81 | -0.98 | 0.70 | 0.60 | -0.06 | -0.59 | -0.39 | 0.40 | -0.72 | -0.98 |
| *rho* | 5.39 | -0.18 | -0.18 | -0.18 | -0.18 | -0.18 | -0.18 | -0.18 | -0.18 | -0.18 | -0.18 | -0.18 | -0.18 | -0.18 | -0.18 | -0.18 | -0.18 | -0.18 |
| *rrh* | 3.98 | 0.14 | -0.79 | 0.30 | -0.73 | -0.66 | -1.17 | -1.48 | -1.60 | -0.05 | -0.40 | 0.96 | -0.31 | 0.33 | -1.26 | 0.36 | 0.17 | -0.52 |
| *tmt1* | 4.52 | -0.09 | -0.50 | 0.73 | -0.57 | -0.44 | -0.54 | -0.72 | -0.70 | -0.29 | -0.40 | 2.33 | -0.49 | 0.03 | -0.38 | -0.21 | -0.41 | -0.40 |

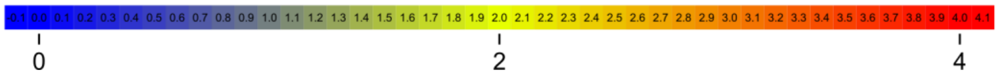

Supplementary Figure 18 Vertebrate Melanopsin-related (Opn4) gene tree inference by ML, rooted by an amphioxus sequence.

The alignment was 735 positions, the best fit model was Q.mammal+G4. Branch supports: SH-aLRT/ultrafast bootstraps (percentage); see main text for species names. The catshark sequences are highlighted in color, gnathostome orthology groups are identified by the position of bullets.

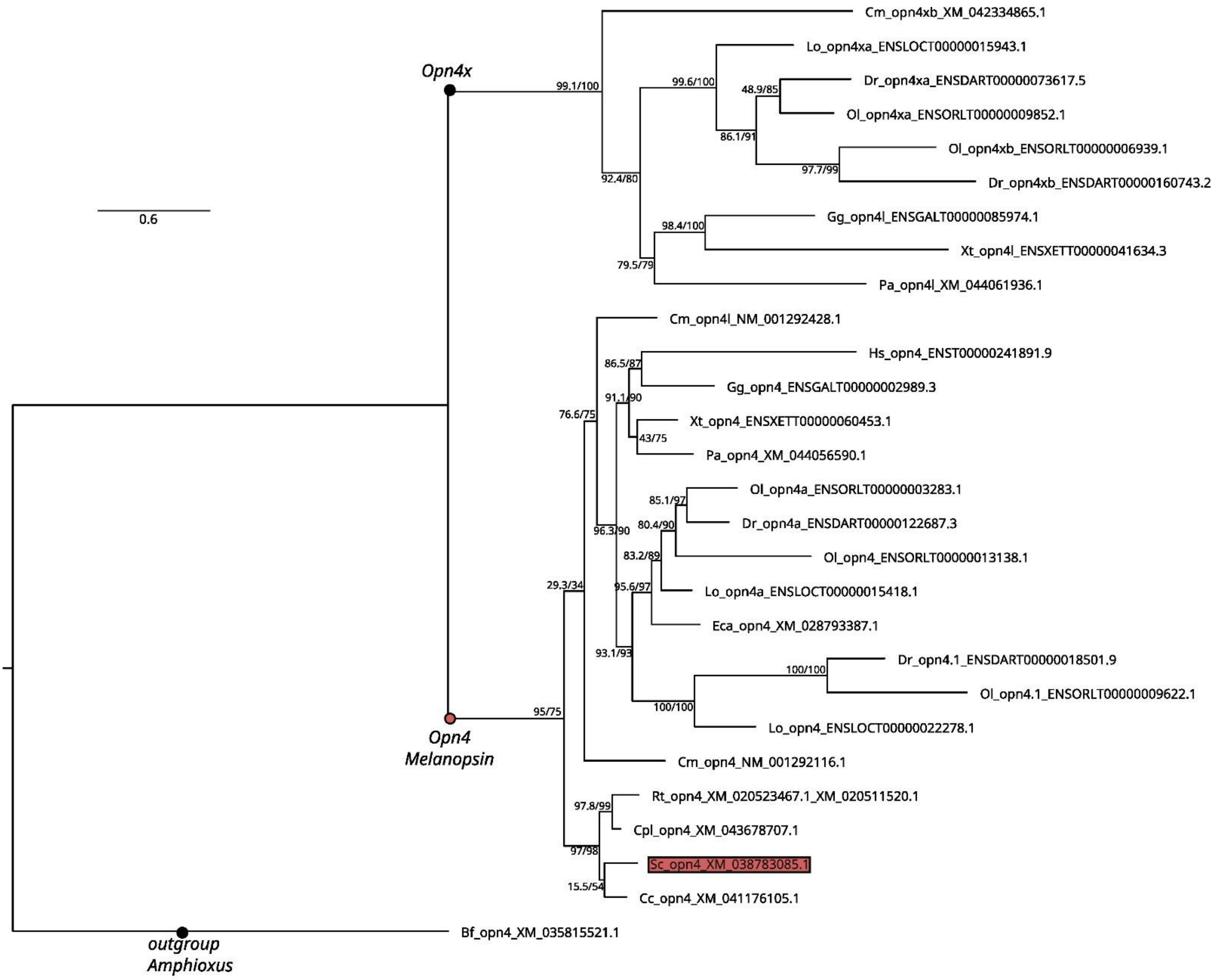

Supplementary Figure 19 Vertebrate Encephalopsin-related (Opn3) gene tree inference by ML, rooted by an amphioxus sequence.

The alignment was 505 positions, the best fit model was Q.plant+I+G4. Branch supports: SH-aLRT/ultrafast bootstraps (percentage); see main text for species names. The catshark sequences are highlighted in color, gnathostome orthology groups are identified by the position of bullets.

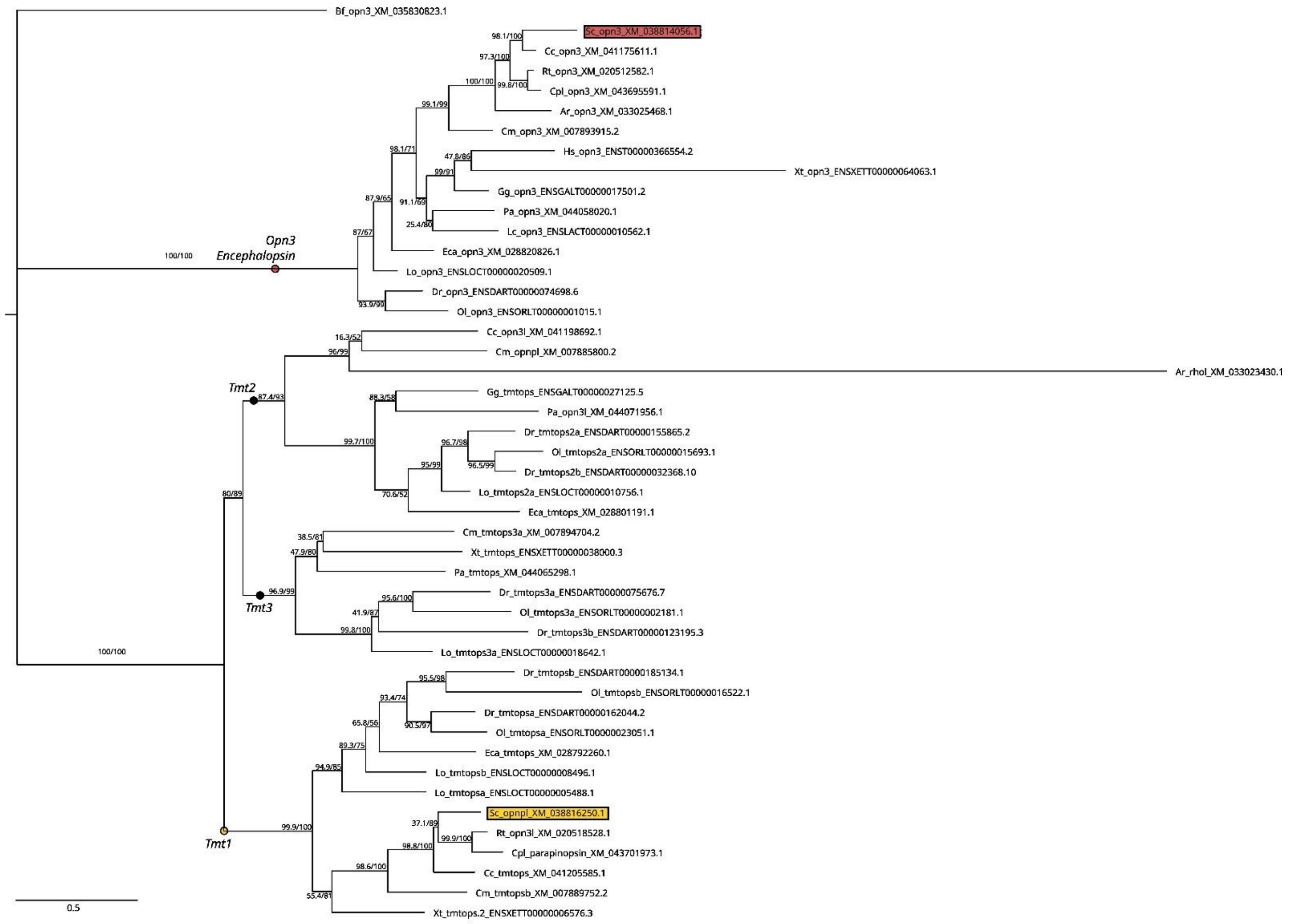

Supplementary Figure 20 Vertebrate Neuropsin-related (Opn5/Opn7) gene tree inference by ML, rooted by an amphioxus sequence.

The alignment was 719 positions, the best fit model was Q.plant+R5. Branch supports: SH-aLRT/ultrafast bootstraps (percentage); see main text for species names. The catshark sequences are highlighted in color, gnathostome orthology groups are identified by the position of bullets.

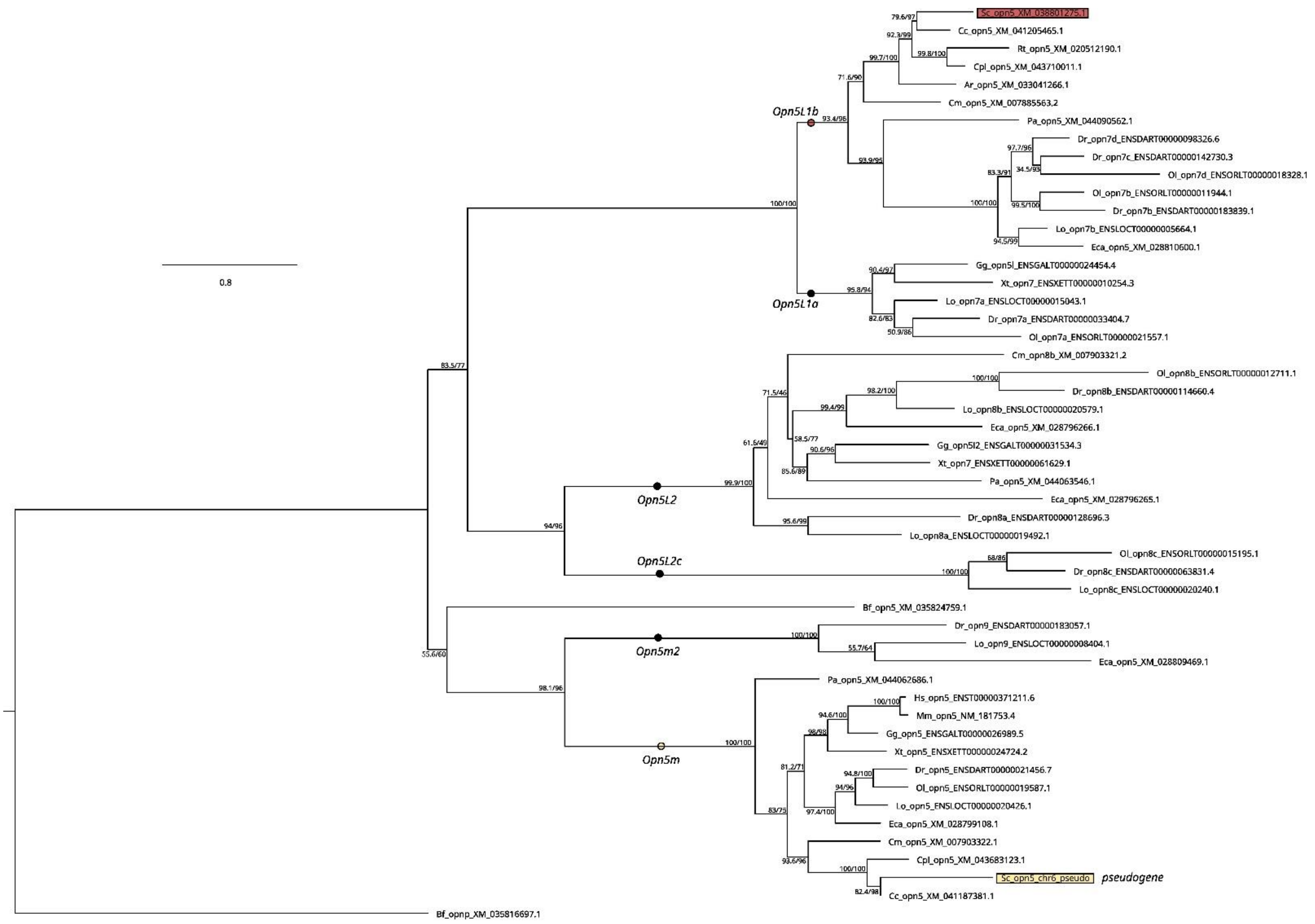

Supplementary Figure 21 Jawed vertebrate Peropsin-related (Rrh) gene tree inference by ML, rooted by a hagfish (Eb) sequence.

The alignment was 439 positions, the best fit model was Q.bird+G4. Branch supports: SH-aLRT/ultrafast bootstraps (percentage); see main text for species names. The catshark sequences are highlighted in color.

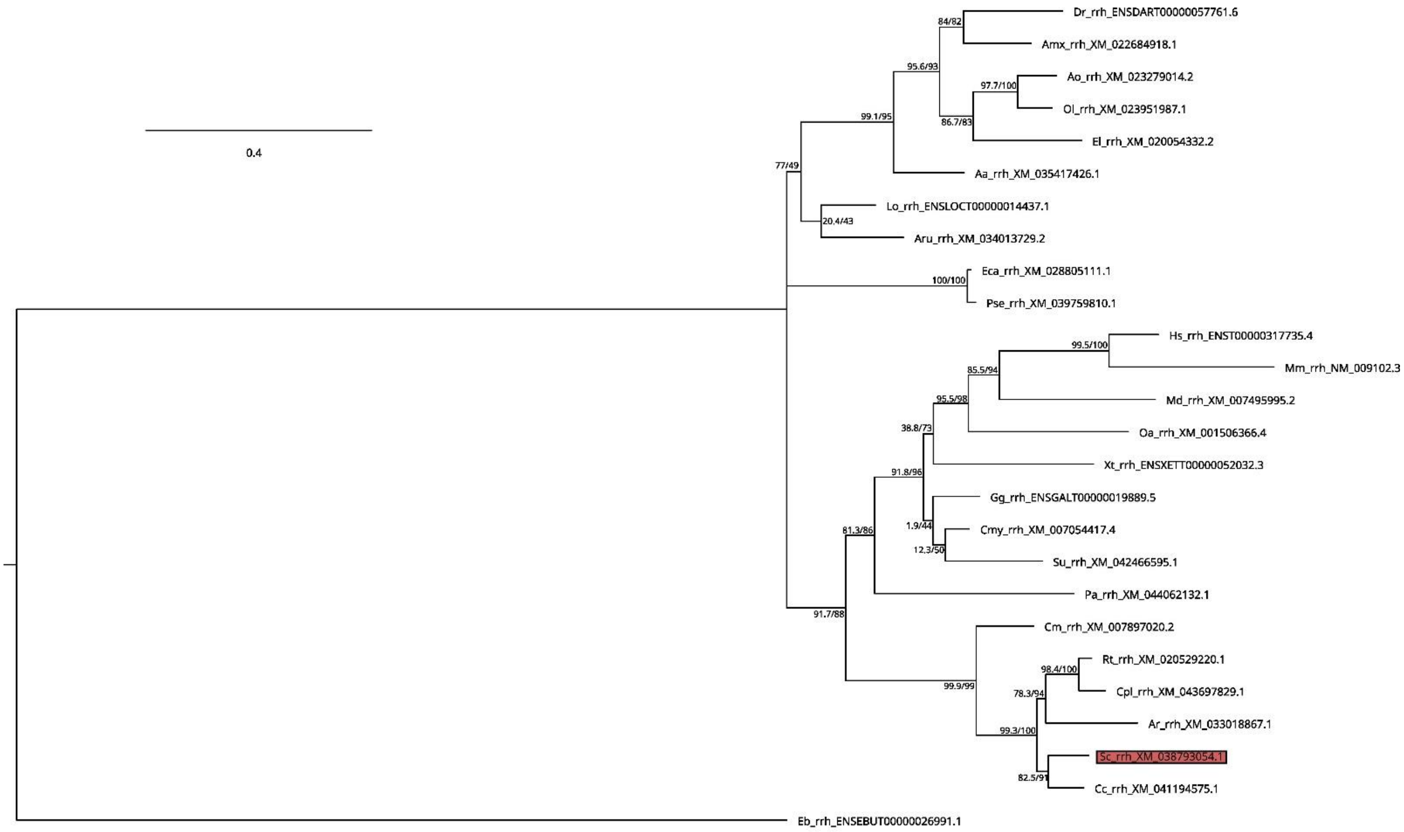

Supplementary Figure 22 Vertebrate RGR-opsin related gene tree inference by ML, rooted by amphioxus sequences. The alignment was 478 positions, the best fit model was Q.plant+G4. Branch supports: SH-aLRT/ultrafast bootstraps (percentage); see main text for species names. The catshark sequences are highlighted in color, gnathostome orthology groups are identified by the position of bullets.

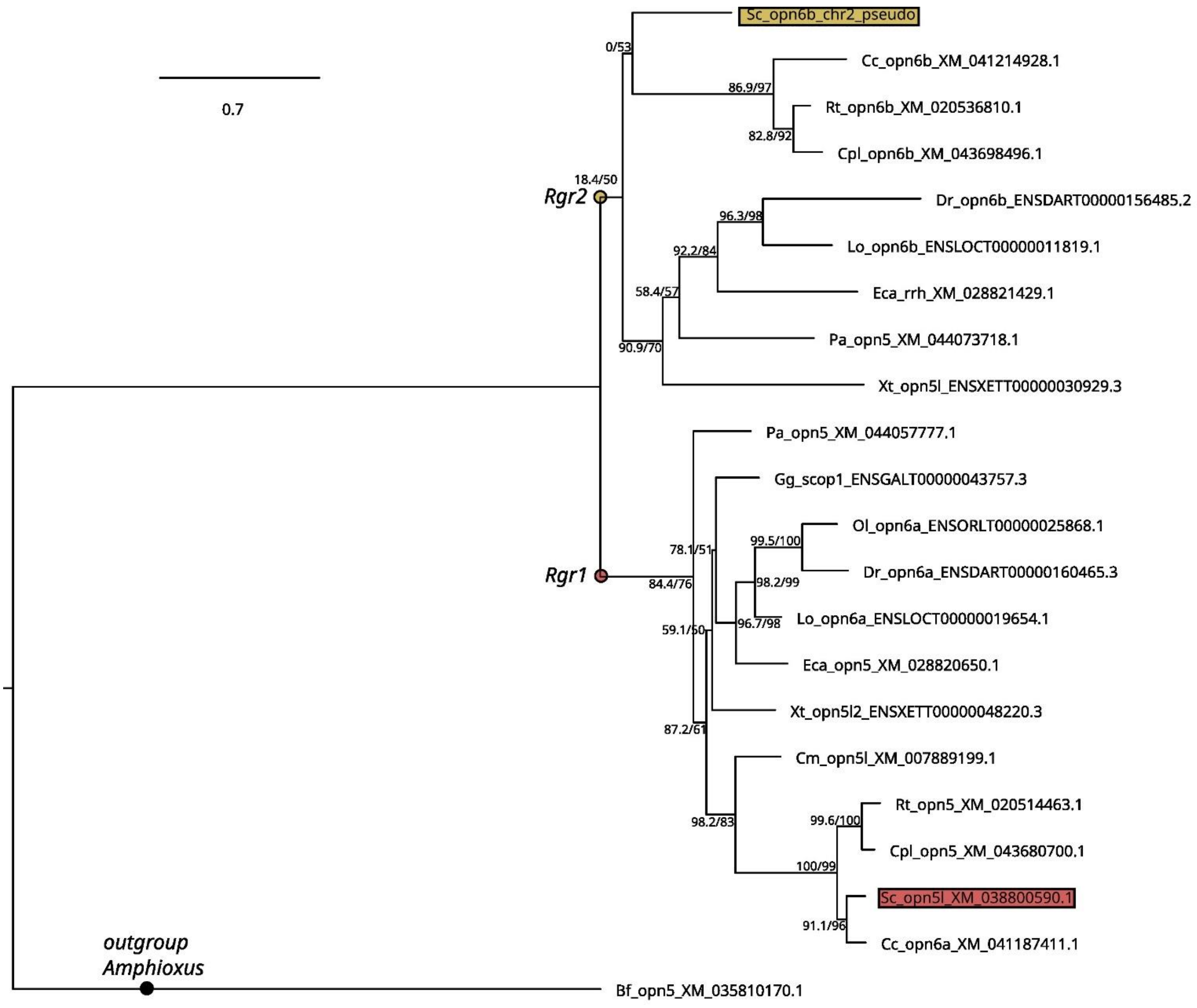

#### Supplementary Figure 23 Gnathostome CryG (Crystallin G) gene tree inference by ML, rooted by the clade of vertebrate CryGN sequences.

The alignment was 820 positions, the best fit model was Q.plant+R6. Branch supports: SH-aLRT/ultrafast bootstraps (percentage); see main text for species names. In the CryG clade, the catshark sequences are highlighted in blue, other clades made of sequences from a single species or taxon, are boxed, the clade made of only cartilaginous fish sequences is shown in a grey box.

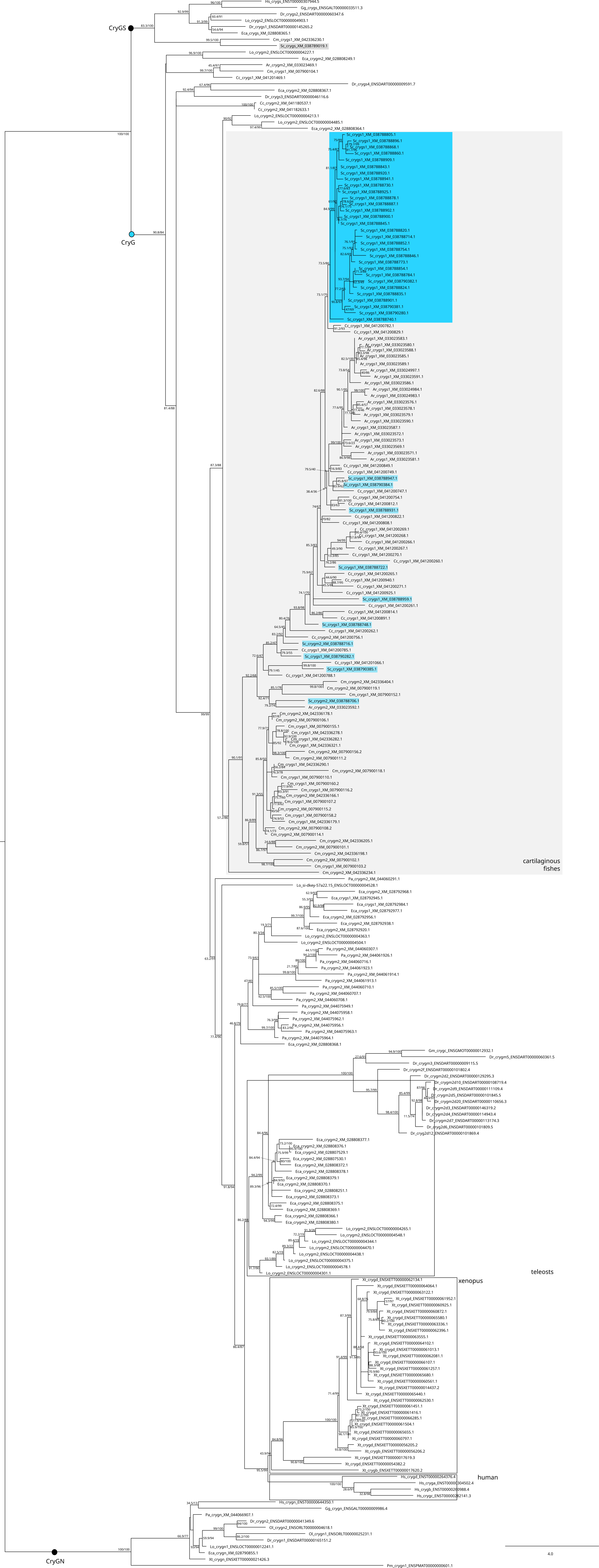
CryGS
CryG
CryGN
cartilaginous fishes
teleosts
xenopus
human
4.0

Supplementary Figure 24 Cartilaginous fish *cryg* gene tree inference by ML, rooted by bony fish *cryg* sequences.
The alignment was 426 codon sites, the best fit model was MG+F1X4+R9. Branch supports: SH-aLRT/ultrafast bootstraps (percentage); see main text for species names. The elephant shark (Cm) and skate (Ar) clades are boxed in grey, catshark sequences grouped in a clade are in a light blue box, catshark sequences with closer relationship with the great white shark (Cc) sequences are in deep blue boxes.

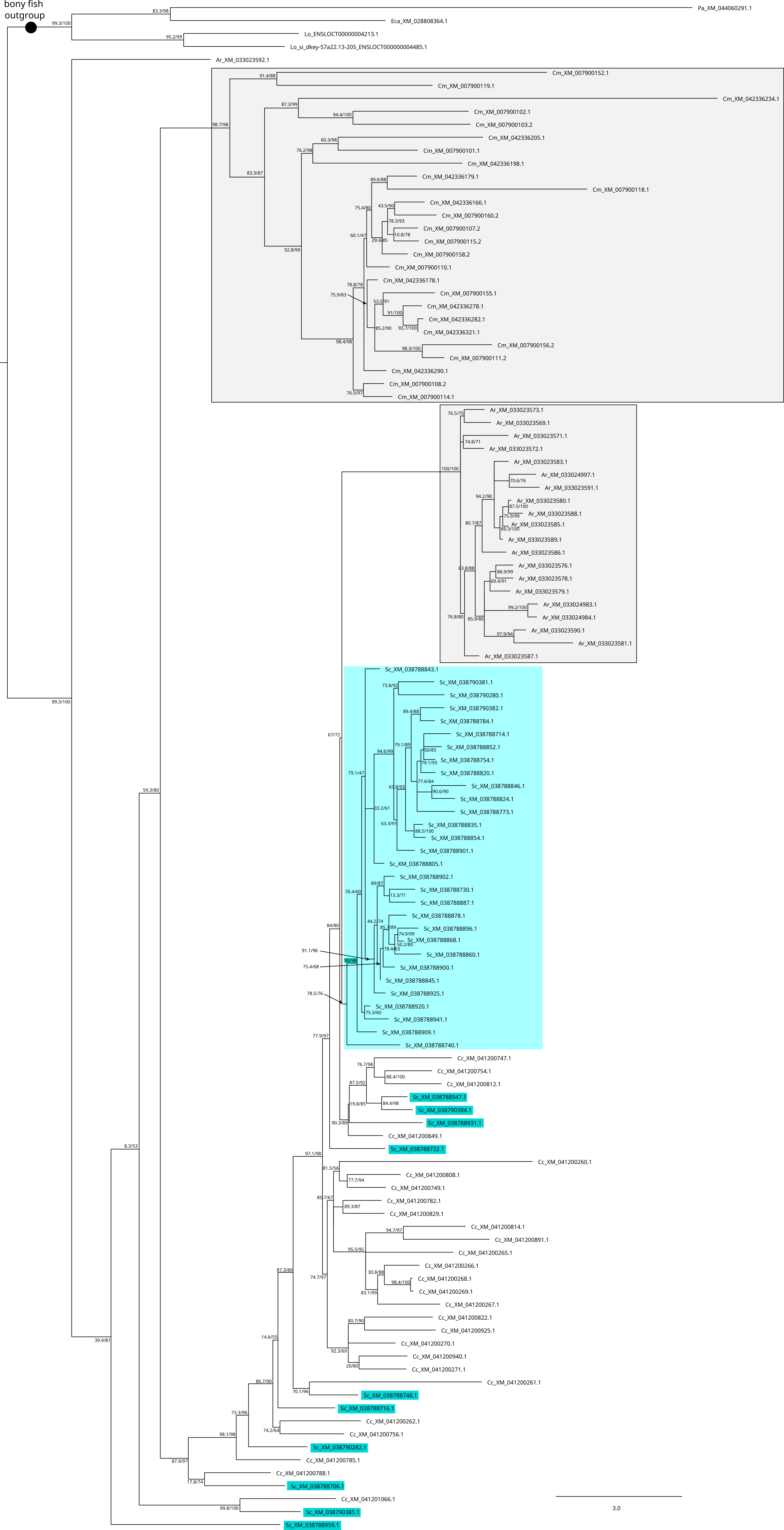

bony fish outgroup
Pa_XM_044060291.1
Eca_XM_028808364.1
Lo_ENSLOCT00000004213.1
Lo_si_dkey-57a22.13-205_ENSLOCT000000004485.1
Ar_XM_033023592.1
Cm_XM_007900152.1
Cm_XM_007900119.1
Cm_XM_042336234.1
Cm_XM_007900102.1
Cm_XM_007900103.2
Cm_XM_042336205.1
Cm_XM_007900101.1
Cm_XM_042336198.1
Cm_XM_042336179.1
Cm_XM_007900118.1
Cm_XM_042336166.1
Cm_XM_007900160.2
Cm_XM_007900107.2
Cm_XM_007900115.2
Cm_XM_007900158.2
Cm_XM_007900110.1
Cm_XM_042336178.1
Cm_XM_007900155.1
Cm_XM_042336278.1
Cm_XM_042336282.1
Cm_XM_042336321.1
Cm_XM_007900156.2
Cm_XM_007900111.2
Cm_XM_042336290.1
Cm_XM_007900108.2
Cm_XM_007900114.1
Ar_XM_033023573.1
Ar_XM_033023569.1
Ar_XM_033023571.1
Ar_XM_033023572.1
Ar_XM_033023583.1
Ar_XM_033024997.1
Ar_XM_033023591.1
Ar_XM_033023580.1
Ar_XM_033023588.1
Ar_XM_033023585.1
Ar_XM_033023589.1
Ar_XM_033023586.1
Ar_XM_033023576.1
Ar_XM_033023578.1
Ar_XM_033023579.1
Ar_XM_033024983.1
Ar_XM_033024984.1
Ar_XM_033023590.1
Ar_XM_033023581.1
Ar_XM_033023587.1
Sc_XM_038788843.1
Sc_XM_038790381.1
Sc_XM_038790280.1
Sc_XM_038790382.1
Sc_XM_038788784.1
Sc_XM_038788714.1
Sc_XM_038788852.1
Sc_XM_038788754.1
Sc_XM_038788820.1
Sc_XM_038788846.1
Sc_XM_038788824.1
Sc_XM_038788773.1
Sc_XM_038788835.1
Sc_XM_038788854.1
Sc_XM_038788901.1
Sc_XM_038788805.1
Sc_XM_038788902.1
Sc_XM_038788730.1
Sc_XM_038788887.1
Sc_XM_038788878.1
Sc_XM_038788896.1
Sc_XM_038788868.1
Sc_XM_038788860.1
Sc_XM_038788900.1
Sc_XM_038788845.1
Sc_XM_038788925.1
Sc_XM_038788920.1
Sc_XM_038788941.1
Sc_XM_038788909.1
Sc_XM_038788740.1
Cc_XM_041200747.1
Cc_XM_041200754.1
Cc_XM_041200812.1
Sc_XM_038788947.1
Sc_XM_038790384.1
Sc_XM_038788931.1
Cc_XM_041200849.1
Sc_XM_038788722.1
Cc_XM_041200260.1
Cc_XM_041200808.1
Cc_XM_041200749.1
Cc_XM_041200782.1
Cc_XM_041200829.1
Cc_XM_041200814.1
Cc_XM_041200891.1
Cc_XM_041200265.1
Cc_XM_041200266.1
Cc_XM_041200268.1
Cc_XM_041200269.1
Cc_XM_041200267.1
Cc_XM_041200822.1
Cc_XM_041200925.1
Cc_XM_041200270.1
Cc_XM_041200940.1
Cc_XM_041200271.1
Cc_XM_041200261.1
Sc_XM_038788748.1
Sc_XM_038788716.1
Cc_XM_041200262.1
Cc_XM_041200756.1
Sc_XM_038790282.1
Cc_XM_041200785.1
Cc_XM_041200788.1
Sc_XM_038788706.1
Cc_XM_041201066.1
Sc_XM_038790385.1
Sc_XM_038788959.1
3.0

**Supplementary Figure 25** Metazoan Trp gene tree inference by ML, rooted by a clade of Trpc-related bilaterian sequences. The alignment was 2431 positions, the best fit model was VT+F+R8. Branch supports: SH-aLRT/ultrafast bootstraps (percentage); see main text for species names, additional species names used here are other bilaterians (Ce: *Caenorhabditis elegans*, Cg: *Crassostrea gigas*, Ct: *Capitella teletea*, Dm: *Drosophila melanogaster*, Pd: *Platynereis dumerilii*; Sk: *Saccoglossus kowalevskii*) and cnidarians (Hv: *Hydra vulgaris*, Nv: *Nematostella vectensis*). The catshark sequences are highlighted in color, gnathostome orthology groups are identified by the position of bullets.

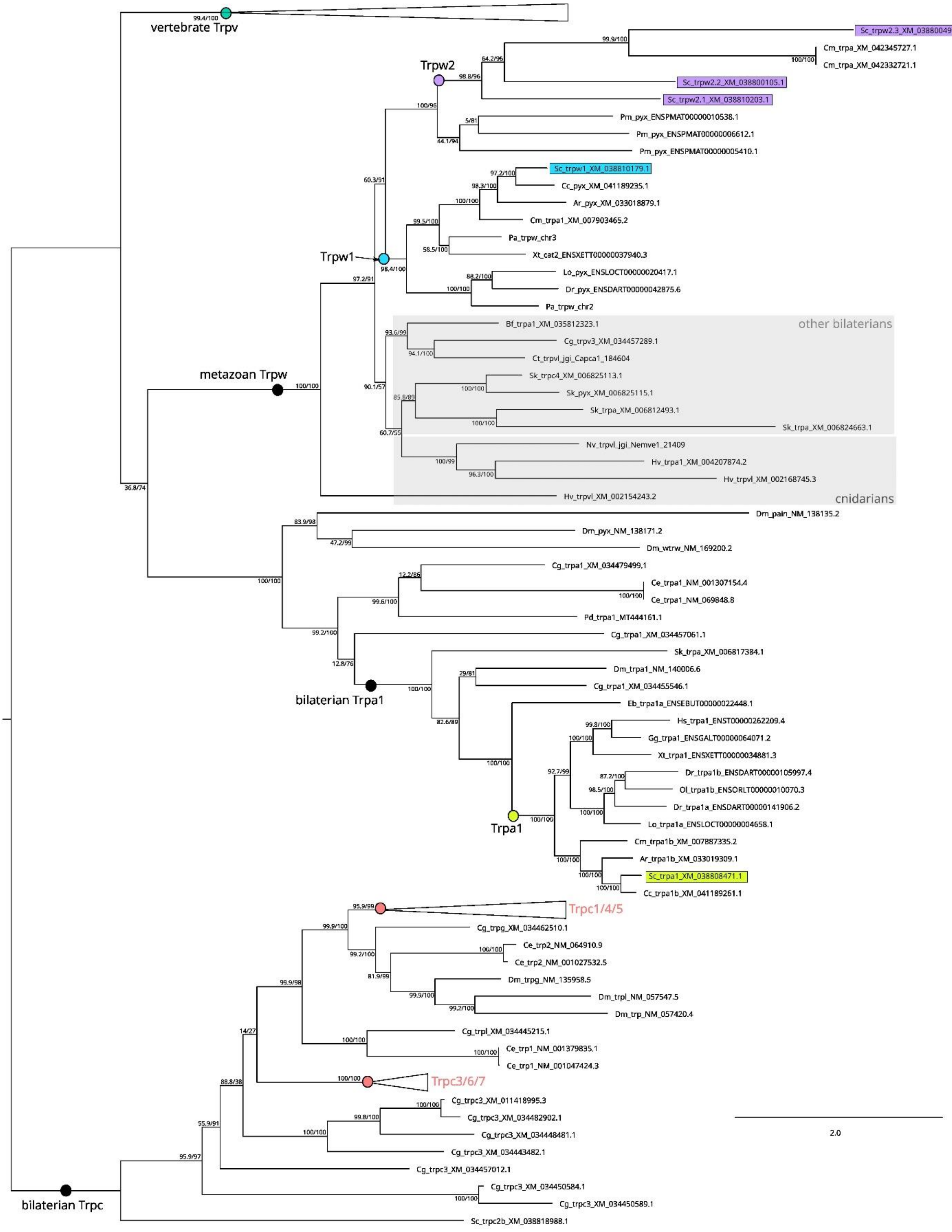

Supplementary Figure 26 Levels of expression (TPM values) of TRP gene families in the reference RNAseq. Values are highlighted depending on the associated Z-score value (scale on the bottom side of the panel).

| | Embr. stage 12 | Embr. Stage 22 | Embr. Stage 24 | Embr. Stage 26 | Embr. Stage 30 | Embr. Stage 31 | AOL | Blood | Brain+olfact. sac | Chondrocranium | Dental lamina | Denticles | Esophagus | Eye | Gills | Heart | Kidney | Liver | Meckels | Muscle | Ovary | Pancreas | Rectal gland | Seminal vesicle | Spinal chord | Spiral intestine | Spleen | Stomach | Testis | Uterus | Vertebrae |
|---|---|---|---|---|---|---|---|---|---|---|---|---|---|---|---|---|---|---|---|---|---|---|---|---|---|---|---|---|---|---|---|
| trpc1 | -0.76 | 0.08 | 0.76 | 0.61 | 1.29 | 0.57 | 0.63 | -1.05 | 0.86 | 1.50 | 0.86 | -0.80 | -0.30 | 0.55 | 0.19 | -0.36 | -0.51 | -1.41 | 1.48 | -1.06 | -0.71 | -1.68 | 1.58 | -0.35 | 0.85 | -1.34 | -0.58 | -1.53 | -0.46 | -0.62 | 1.66 |
| trpc2 | -0.21 | -0.21 | -0.21 | -0.21 | -0.21 | -0.21 | 0.97 | -0.22 | 5.27 | -0.21 | -0.21 | -0.22 | -0.22 | -0.21 | -0.21 | -0.22 | -0.22 | -0.22 | -0.22 | -0.22 | -0.22 | -0.22 | -0.22 | -0.22 | -0.22 | -0.22 | -0.22 | -0.22 | -0.21 | -0.22 | -0.21 |
| trpc3 | 2.64 | 1.17 | 0.77 | 0.67 | 0.24 | 0.12 | -0.63 | -0.39 | 3.06 | 0.68 | -0.30 | -0.16 | -0.57 | -0.47 | -0.31 | -0.75 | -0.28 | -0.72 | -0.59 | -0.87 | -0.98 | -0.94 | -0.47 | -0.70 | 1.34 | -0.67 | -0.13 | -0.57 | 1.08 | -0.79 | -0.51 |
| trpc4 | -0.58 | 0.35 | 0.99 | 0.75 | 1.67 | 1.18 | -0.55 | -0.82 | 0.85 | -0.51 | -0.67 | 1.23 | -0.72 | 2.75 | -0.75 | 0.06 | -0.34 | -0.77 | -0.55 | -0.78 | -0.82 | -0.73 | 0.14 | -0.75 | 2.49 | -0.62 | -0.41 | -0.51 | -0.47 | -0.44 | -0.64 |
| trpc5 | 0.56 | -0.26 | 0.03 | 0.07 | 0.92 | 0.82 | -0.25 | -0.32 | 1.72 | 0.62 | -0.55 | -0.42 | -0.60 | 1.66 | -0.63 | -0.63 | 0.68 | -0.84 | -0.33 | -0.43 | -0.96 | -1.00 | -0.59 | -0.76 | 3.59 | -0.78 | -0.56 | -0.91 | 0.94 | -0.60 | -0.19 |
| trpc6 | -0.52 | -0.23 | -0.38 | -0.54 | 0.56 | 1.08 | -0.42 | -0.71 | 0.02 | 0.08 | -0.51 | -0.65 | -0.60 | -0.37 | -0.69 | 0.63 | 0.78 | -0.70 | -0.23 | 0.28 | -0.52 | -0.68 | 4.62 | 0.50 | 0.55 | -0.50 | 0.36 | -0.68 | 0.14 | -0.44 | -0.23 |
| trpc7 | -0.39 | -0.26 | 0.09 | -0.05 | 0.45 | -0.01 | -0.30 | -0.27 | 0.11 | -0.30 | -0.06 | 0.23 | -0.09 | -0.17 | 0.01 | -0.36 | 5.19 | -0.45 | -0.34 | -0.45 | -0.51 | -0.51 | -0.13 | -0.47 | -0.08 | -0.05 | 0.58 | -0.45 | -0.20 | -0.37 | -0.38 |
| trpm1 | -0.27 | 0.28 | 4.45 | 0.64 | 0.56 | 0.01 | -0.05 | -0.06 | 0.38 | -0.19 | 0.26 | -0.03 | -0.52 | 0.10 | 0.38 | -0.61 | 0.50 | -0.78 | -0.61 | -1.06 | -1.17 | -1.31 | -0.13 | -0.11 | 0.78 | -0.60 | 0.48 | -1.08 | 0.38 | -0.12 | -0.50 |
| trpm1 | -0.27 | -0.22 | -0.09 | -0.22 | -0.15 | -0.21 | -0.18 | -0.26 | 0.03 | 0.04 | -0.16 | -0.23 | -0.29 | 5.33 | -0.22 | -0.29 | -0.12 | -0.34 | -0.18 | -0.32 | -0.06 | -0.38 | -0.17 | -0.29 | -0.02 | -0.29 | 0.31 | -0.35 | 0.09 | -0.28 | -0.16 |
| trpm2 | -0.27 | -0.28 | -0.26 | -0.27 | -0.26 | -0.24 | 0.71 | -0.27 | 5.20 | 0.61 | -0.25 | -0.22 | -0.23 | -0.24 | -0.21 | -0.26 | -0.26 | -0.29 | -0.27 | -0.29 | -0.29 | -0.29 | -0.27 | -0.28 | -0.25 | -0.28 | -0.19 | -0.29 | -0.26 | -0.29 | 0.53 |
| trpm3 | -0.33 | -0.22 | 0.34 | -0.01 | 0.01 | 0.07 | -0.16 | -0.34 | 1.04 | 0.30 | -0.33 | -0.34 | -0.33 | 5.13 | -0.34 | -0.35 | -0.28 | -0.35 | -0.34 | -0.33 | -0.37 | -0.37 | -0.34 | -0.28 | 0.20 | -0.34 | -0.34 | -0.37 | -0.29 | -0.35 | 0.03 |
| trpm4 | -0.48 | -0.30 | -0.39 | -0.44 | -0.39 | -0.20 | -0.08 | -0.53 | -0.38 | -0.04 | 4.01 | 2.11 | 0.63 | -0.33 | 1.21 | -0.53 | -0.50 | -0.53 | -0.01 | -0.47 | -0.54 | -0.52 | -0.37 | -0.24 | -0.52 | -0.38 | -0.52 | 1.79 | -0.51 | -0.08 | -0.48 |
| trpm5 | -0.60 | -0.45 | -0.41 | -0.40 | -0.31 | -0.15 | 0.46 | 0.00 | -0.13 | -0.43 | 0.67 | 1.08 | -0.07 | 0.75 | 3.88 | -0.59 | 1.39 | -0.64 | -0.54 | -0.88 | -0.95 | -0.94 | -0.13 | -0.64 | -0.39 | -0.41 | -0.09 | 2.11 | -0.26 | -0.26 | -0.68 |
| trpm6 | -0.54 | -0.50 | -0.21 | -0.39 | 0.15 | 0.30 | 3.39 | -0.25 | 0.17 | 0.40 | 0.65 | 0.17 | -0.34 | 2.44 | 0.11 | -0.49 | 1.70 | -0.75 | -0.27 | -0.98 | -1.22 | -1.23 | -0.37 | -0.62 | -0.18 | 1.04 | 0.02 | -1.03 | -0.36 | -0.34 | -0.47 |
| trpm7 | 0.77 | 0.03 | -0.35 | 0.20 | 0.35 | 0.13 | -0.03 | -0.41 | -0.38 | 0.86 | 1.00 | 0.77 | -0.86 | 1.32 | 0.41 | -0.52 | -0.05 | -0.96 | 2.70 | -1.46 | -1.61 | -1.72 | 0.04 | -0.77 | -0.88 | 0.97 | 0.42 | -1.01 | -0.63 | -0.26 | 1.92 |
| trpv1/2 | -0.72 | -0.78 | -0.44 | -0.50 | -0.29 | 0.18 | -0.32 | -0.68 | 0.12 | 2.95 | 0.73 | 1.84 | 1.24 | -0.59 | 0.91 | -0.59 | -0.45 | -0.76 | -0.49 | -0.72 | -0.78 | -0.78 | -0.47 | -0.65 | 0.78 | -0.31 | 0.05 | -0.54 | -0.12 | -0.52 | 2.72 |
| trpv3 | -0.12 | -0.37 | -0.33 | 0.31 | 0.39 | 0.64 | 0.33 | 0.48 | 1.22 | -0.62 | 0.46 | 0.46 | 0.89 | -0.44 | 3.51 | -0.82 | -0.23 | -0.74 | -0.61 | -0.92 | -1.05 | -1.03 | 0.74 | -0.68 | -0.46 | -0.54 | -0.35 | -1.03 | 2.22 | -0.60 | -0.72 |
| trpv4 | -0.28 | 0.01 | 0.27 | 0.05 | 0.24 | 0.01 | -0.25 | -0.08 | -0.19 | 0.03 | 0.14 | 0.03 | -0.37 | -0.16 | -0.13 | -0.50 | -0.10 | -0.58 | -0.24 | -0.62 | -0.67 | -0.76 | -0.06 | 5.10 | -0.24 | -0.47 | -0.04 | -0.68 | -0.03 | -0.23 | 0.77 |
| trpv5/6.1 | -0.40 | -0.34 | -0.15 | -0.33 | 0.19 | 1.85 | -0.49 | -0.48 | -0.27 | 0.11 | -0.48 | -0.45 | -0.46 | 0.07 | 3.86 | -0.24 | 2.78 | -0.49 | -0.47 | 0.26 | -0.51 | -0.50 | -0.19 | 0.00 | -0.22 | -0.50 | -0.47 | -0.46 | -0.40 | -0.50 | -0.31 |
| trpv5/6.2 | -0.32 | -0.39 | -0.34 | -0.23 | -0.17 | 1.61 | 0.32 | -0.23 | 2.64 | -0.23 | -0.48 | -0.54 | -0.46 | -0.07 | 1.53 | -0.51 | -0.06 | -0.57 | -0.51 | -0.57 | -0.57 | -0.55 | 0.15 | 3.66 | -0.40 | -0.54 | -0.37 | -0.57 | -0.26 | -0.57 | -0.42 |
| trpv5/6.3 | 3.07 | 0.84 | 0.20 | 0.99 | 0.86 | 0.42 | -0.44 | -0.43 | -0.42 | -0.44 | -0.44 | -0.42 | -0.43 | -0.23 | 3.70 | -0.44 | -0.42 | -0.45 | -0.42 | -0.45 | -0.44 | -0.45 | -0.42 | -0.42 | -0.42 | -0.44 | -0.42 | -0.44 | -0.35 | -0.40 | -0.45 |
| trpv7 | 1.54 | 1.73 | -1.35 | -1.36 | -1.29 | -1.31 | 0.24 | -1.33 | 0.59 | 0.21 | 0.29 | 0.66 | 1.26 | 0.00 | 2.10 | 0.72 | 1.05 | -0.25 | 0.36 | -0.43 | -1.39 | -1.19 | 0.26 | 0.36 | 0.85 | -0.20 | -0.33 | -0.29 | 0.37 | -1.38 | -0.50 |
| trpv8.1 | -0.54 | -0.45 | 1.07 | 0.08 | 0.81 | 0.67 | 1.46 | -0.34 | 2.77 | 0.57 | 0.21 | -0.55 | -0.14 | -0.12 | 1.76 | -0.61 | -0.46 | -1.10 | -0.26 | -0.73 | -1.59 | -1.67 | 0.05 | -0.16 | 0.22 | -0.87 | -0.15 | -1.41 | 1.70 | -0.24 | 0.03 |
| trpv8.2 | -0.55 | -0.22 | 0.15 | -0.33 | 0.18 | 0.26 | 2.48 | -0.16 | 1.11 | 0.06 | 0.07 | 0.02 | 0.08 | -0.09 | 4.08 | -0.50 | -0.11 | -0.66 | -0.41 | -0.79 | -0.96 | -0.89 | -0.23 | -0.31 | -0.41 | -0.53 | 0.01 | -0.91 | 0.54 | -0.51 | -0.44 |
| trpv9 | -0.66 | -0.12 | 0.45 | 0.11 | 0.63 | 0.04 | 0.27 | 0.91 | 0.42 | 0.15 | 0.02 | -0.05 | 0.11 | 2.28 | 0.80 | -0.45 | 0.18 | -1.12 | -0.58 | -1.41 | -1.64 | -1.81 | 0.37 | -0.71 | 0.92 | -0.68 | 2.16 | -1.59 | 1.68 | -0.26 | -0.42 |
| pkd1 | -0.64 | -0.39 | -0.41 | -0.21 | 0.04 | 0.13 | -0.49 | -0.90 | -0.43 | -0.34 | -0.02 | -0.50 | -0.51 | -0.16 | -0.19 | 2.51 | 1.82 | -0.54 | 0.64 | -0.62 | -0.82 | -0.72 | 3.17 | -0.07 | 0.19 | -0.73 | -0.37 | 0.12 | -0.71 | -0.73 | 1.89 |
| pkdrej | -0.23 | -0.20 | -0.20 | -0.20 | -0.18 | -0.15 | -0.18 | -0.13 | -0.11 | -0.18 | -0.19 | -0.19 | -0.18 | -0.18 | -0.13 | -0.18 | -0.17 | -0.20 | -0.21 | -0.23 | -0.22 | -0.23 | -0.18 | -0.19 | -0.18 | -0.19 | 0.06 | -0.22 | 5.38 | -0.18 | -0.21 |
| pkd1L2 | -0.34 | -0.35 | -0.30 | -0.32 | -0.01 | 0.17 | -0.33 | -0.33 | -0.29 | -0.29 | -0.33 | -0.28 | -0.34 | -0.28 | -0.33 | -0.35 | -0.09 | -0.32 | -0.33 | -0.35 | -0.34 | -0.35 | 3.11 | -0.35 | 4.22 | 0.01 | -0.33 | -0.10 | -0.31 | 0.16 | -0.34 |
| pkd1L3.1 | -0.93 | -0.84 | 0.04 | -0.50 | -0.44 | -0.44 | 0.46 | -0.64 | 0.07 | 0.02 | 0.15 | -0.58 | -0.10 | 1.10 | -0.02 | 1.50 | 2.22 | 0.34 | -0.47 | -1.31 | -0.85 | -1.56 | 1.23 | 0.05 | 1.44 | -0.98 | 0.22 | -1.08 | 2.63 | -0.01 | -0.68 |
| pkd1L3.2 | 1.21 | 0.08 | 2.66 | -0.08 | 0.22 | 0.44 | -0.57 | -0.91 | 0.01 | -0.58 | 2.10 | 0.09 | 1.58 | -0.15 | 0.08 | -1.26 | -0.68 | -0.92 | -0.53 | -1.15 | -1.33 | -1.31 | -0.29 | -0.08 | -0.09 | -0.53 | -0.39 | 0.94 | 1.65 | 0.47 | -0.66 |
| pkd2 | -0.25 | -0.12 | 0.06 | 0.20 | 0.31 | 0.22 | 1.41 | -0.62 | -0.61 | 1.32 | -0.09 | -0.64 | -0.29 | 0.15 | -0.69 | 0.14 | 2.32 | -1.02 | 1.69 | -1.09 | -1.34 | -1.43 | 1.67 | 1.01 | 0.42 | -1.08 | -0.66 | -1.33 | -0.22 | -0.79 | 1.36 |
| pkd2L1 | -0.32 | -0.17 | 0.25 | -0.04 | 0.52 | 0.49 | -0.20 | 0.34 | -0.01 | -0.12 | 0.11 | -0.16 | -0.23 | 0.30 | 0.01 | -0.45 | 0.12 | -0.47 | -0.45 | -0.75 | -0.98 | -0.99 | -0.16 | -0.37 | 4.94 | -0.41 | 0.32 | -0.83 | 0.25 | -0.22 | -0.32 |
| pkd2L2 | -0.29 | -0.13 | 0.16 | -0.13 | -0.16 | -0.18 | -0.23 | -0.03 | -0.09 | -0.21 | -0.06 | -0.08 | -0.29 | 0.01 | -0.17 | -0.33 | 0.20 | -0.31 | -0.26 | -0.41 | -0.40 | -0.41 | -0.13 | -0.08 | -0.07 | -0.29 | -0.18 | -0.36 | 5.33 | -0.21 | -0.23 |
| mcoln1 | 0.94 | -0.63 | 0.63 | 0.10 | -0.96 | -0.06 | 0.05 | -0.28 | 0.64 | 0.24 | -0.34 | -0.32 | -1.28 | 0.72 | -0.08 | -0.08 | 2.50 | 1.51 | -1.07 | -0.99 | -1.43 | -1.59 | 1.27 | -0.71 | 2.16 | 0.61 | 0.71 | -0.54 | -1.03 | -0.38 | -0.30 |
| mcoln2 | -0.34 | -0.36 | -0.42 | -0.57 | -0.54 | -0.47 | 0.10 | -0.46 | 0.76 | -0.38 | -0.08 | 0.06 | 0.26 | -0.13 | 0.01 | -0.23 | 4.74 | 1.26 | -0.45 | -0.54 | -0.59 | -0.55 | -0.39 | -0.52 | -0.25 | 0.05 | 1.21 | -0.50 | -0.27 | 0.06 | -0.45 |
| trpml3.1 | -0.55 | 0.16 | 0.10 | -0.37 | 0.03 | 0.03 | 2.10 | -0.74 | 1.82 | 0.89 | -0.51 | 0.50 | -0.70 | 0.50 | -0.36 | -0.77 | 2.60 | -0.59 | -0.40 | -0.76 | -0.80 | -0.77 | -0.61 | -0.70 | 2.33 | -0.71 | -0.74 | -0.80 | 0.87 | -0.76 | -0.26 |
| trpml3.2 | 3.83 | 0.46 | 0.20 | -0.53 | -0.22 | 0.21 | 0.11 | -0.65 | 0.39 | 0.06 | 0.18 | 1.99 | -0.72 | -0.12 | -0.44 | -0.66 | 1.87 | -0.52 | -0.12 | -0.73 | -0.66 | -0.72 | -0.70 | -0.73 | 0.41 | -0.71 | -0.66 | -0.77 | 0.71 | -0.68 | -0.08 |
| trpa1 | -0.29 | -0.25 | -0.26 | -0.30 | -0.28 | -0.23 | -0.32 | -0.33 | -0.28 | 3.76 | -0.31 | -0.32 | -0.31 | -0.30 | -0.32 | -0.32 | -0.31 | -0.32 | -0.32 | -0.31 | 0.08 | -0.33 | -0.27 | -0.32 | -0.30 | 0.26 | -0.31 | 0.02 | -0.27 | -0.32 | 3.66 |
| trpw1 | -0.21 | -0.22 | -0.21 | -0.22 | -0.21 | -0.22 | 0.97 | -0.21 | 5.26 | -0.21 | -0.22 | -0.22 | -0.22 | -0.21 | -0.22 | -0.22 | -0.22 | -0.20 | -0.22 | -0.22 | -0.22 | -0.22 | -0.22 | -0.22 | -0.20 | -0.22 | -0.21 | -0.22 | -0.20 | -0.22 | -0.22 |
| trpw2.1 | -0.18 | -0.19 | 0.05 | -0.20 | -0.17 | -0.20 | -0.03 | -0.20 | 0.16 | -0.21 | -0.20 | -0.22 | -0.23 | -0.19 | -0.20 | -0.22 | -0.19 | -0.24 | -0.24 | -0.24 | -0.23 | -0.14 | -0.21 | -0.23 | -0.19 | -0.21 | -0.20 | -0.22 | 5.37 | -0.18 | -0.21 |
| trpw2.2 | -0.22 | -0.66 | 0.22 | -0.35 | -0.06 | -0.44 | 0.25 | -0.01 | 0.99 | -0.28 | 1.44 | 1.54 | -0.45 | 0.04 | 1.34 | -0.84 | -0.10 | -0.93 | -0.60 | -0.43 | -1.30 | -1.62 | 0.09 | -0.40 | -0.54 | -0.05 | 0.18 | -1.13 | 3.34 | 1.42 | -0.43 |
| trpw2.3 | 2.77 | 1.92 | 1.51 | 1.34 | 0.95 | 0.61 | -0.55 | -0.99 | 0.26 | -0.38 | -0.10 | 0.70 | -0.76 | -0.40 | 0.13 | -0.64 | -0.24 | -0.41 | -0.62 | -0.69 | -0.92 | -1.24 | -0.34 | -0.76 | -0.37 | -0.06 | -0.55 | -0.57 | 1.98 | -0.90 | -0.69 |

#### Supplementary Figure 27 Vertebrate Trpc gene tree inference by ML, rooted by an amphioxus sequence.

The alignment was 2097 positions, the best fit model was JTT+R8. Branch supports: SH-aLRT/ultrafast bootstraps (percentage); see main text for species names. The catshark sequences are highlighted in color, gnathostome orthology groups are identified by the position of bullets.

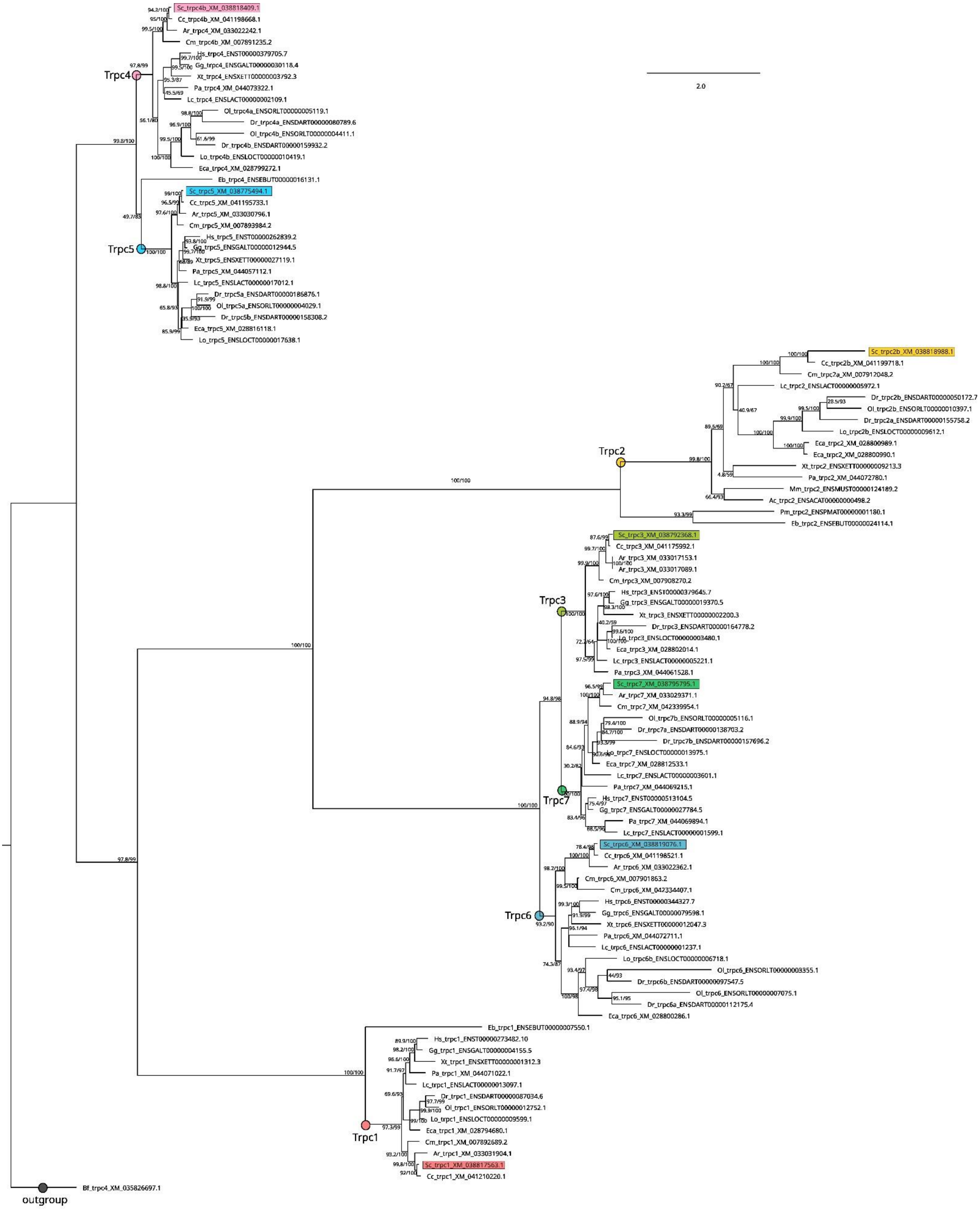

Supplementary Figure 28 Jawed vertebrate Trpn gene tree inference by ML, rooted by an amphioxus sequence. The alignment was 1723 positions, the best fit model was Q.plant+I+R3. Branch supports: SH-aLRT/ultrafast bootstraps (percentage); see main text for species names. The catshark sequences are highlighted in color, the gnathostome orthology group is identified by the position of a bullet.

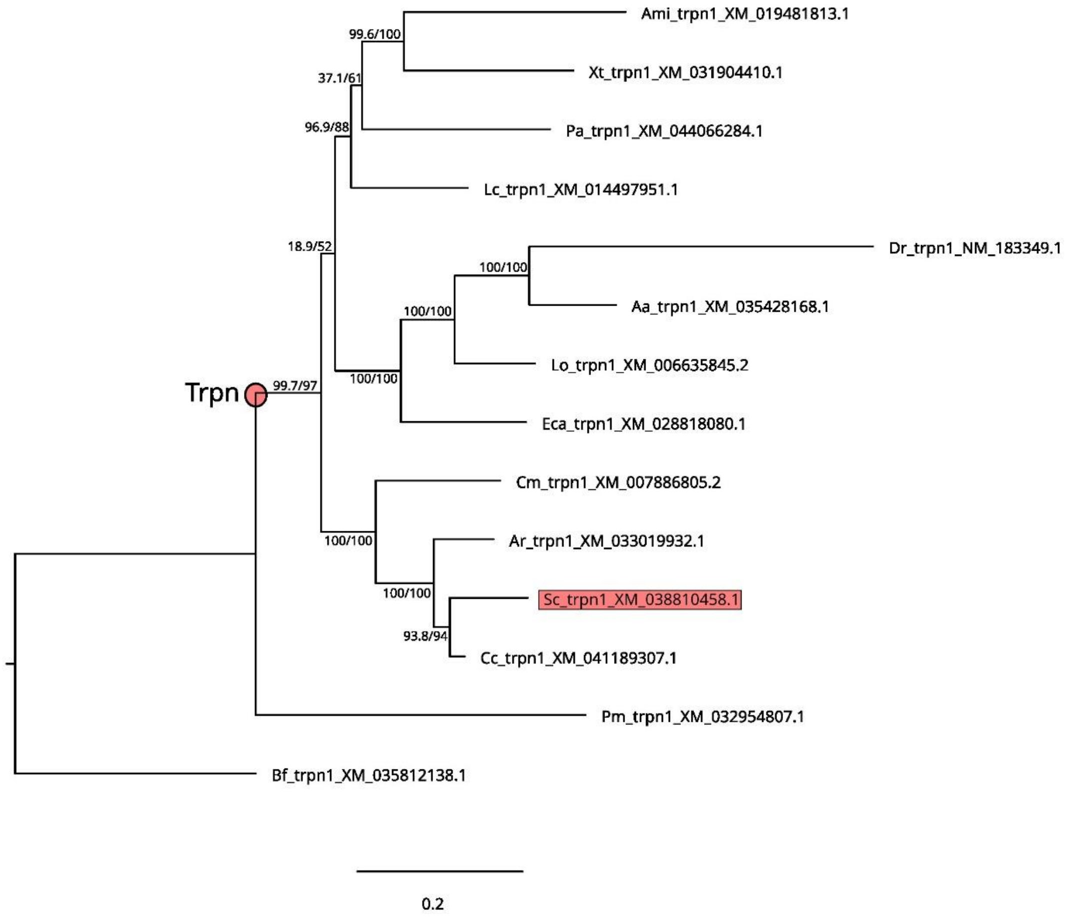

Supplementary Figure 29 Jawed vertebrate Trpa1-related gene tree inference by ML, rooted by an hemichordate sequence (*Saccoglossus kowalevskii*).
The alignment was 1344 positions, the best fit model was Q.plant+F+I+G4. Branch supports: SH-aLRT/ultrafast bootstraps (percentage); see main text for species names. The catshark sequences are highlighted in color, gnathostome orthology groups are identified by the position of bullets.

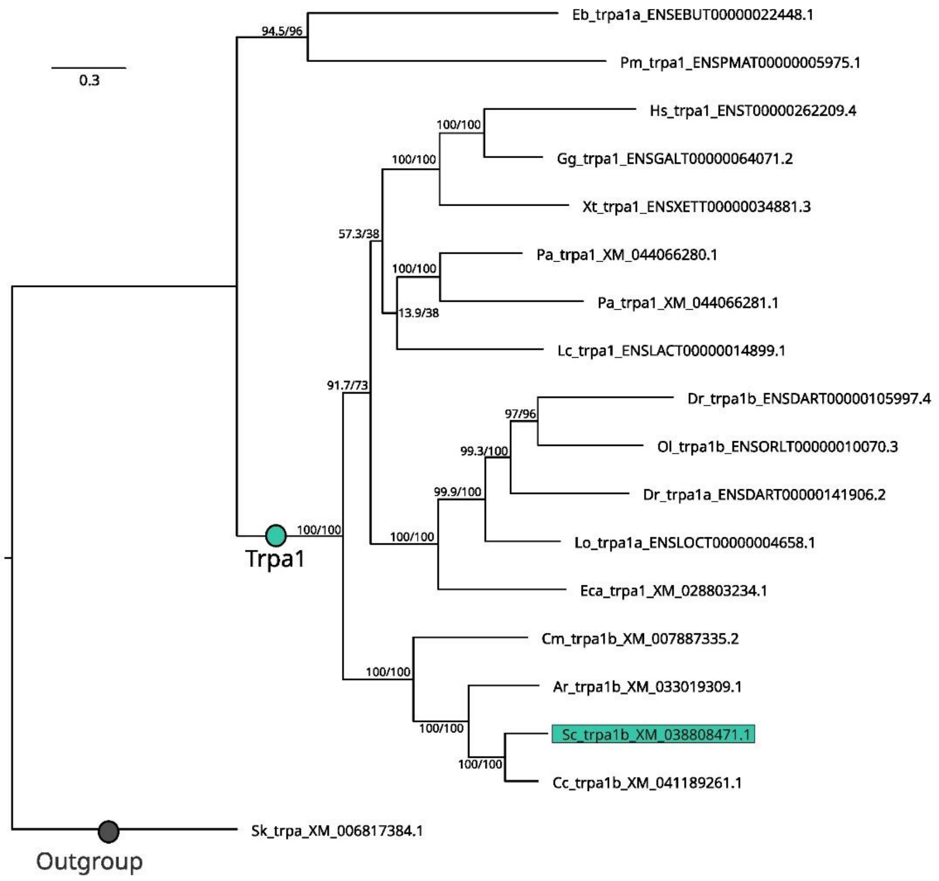

Supplementary Figure 30 Vertebrate Trpm gene tree inference by ML, rooted by an amphioxus sequence. The alignment was 3082 positions, the best fit model was JTT+F+R7. Branch supports: SH-aLRT/ultrafast bootstraps (percentage); see main text for species names. The catshark sequences are highlighted in color, gnathostome orthology groups are identified by the position of bullets.

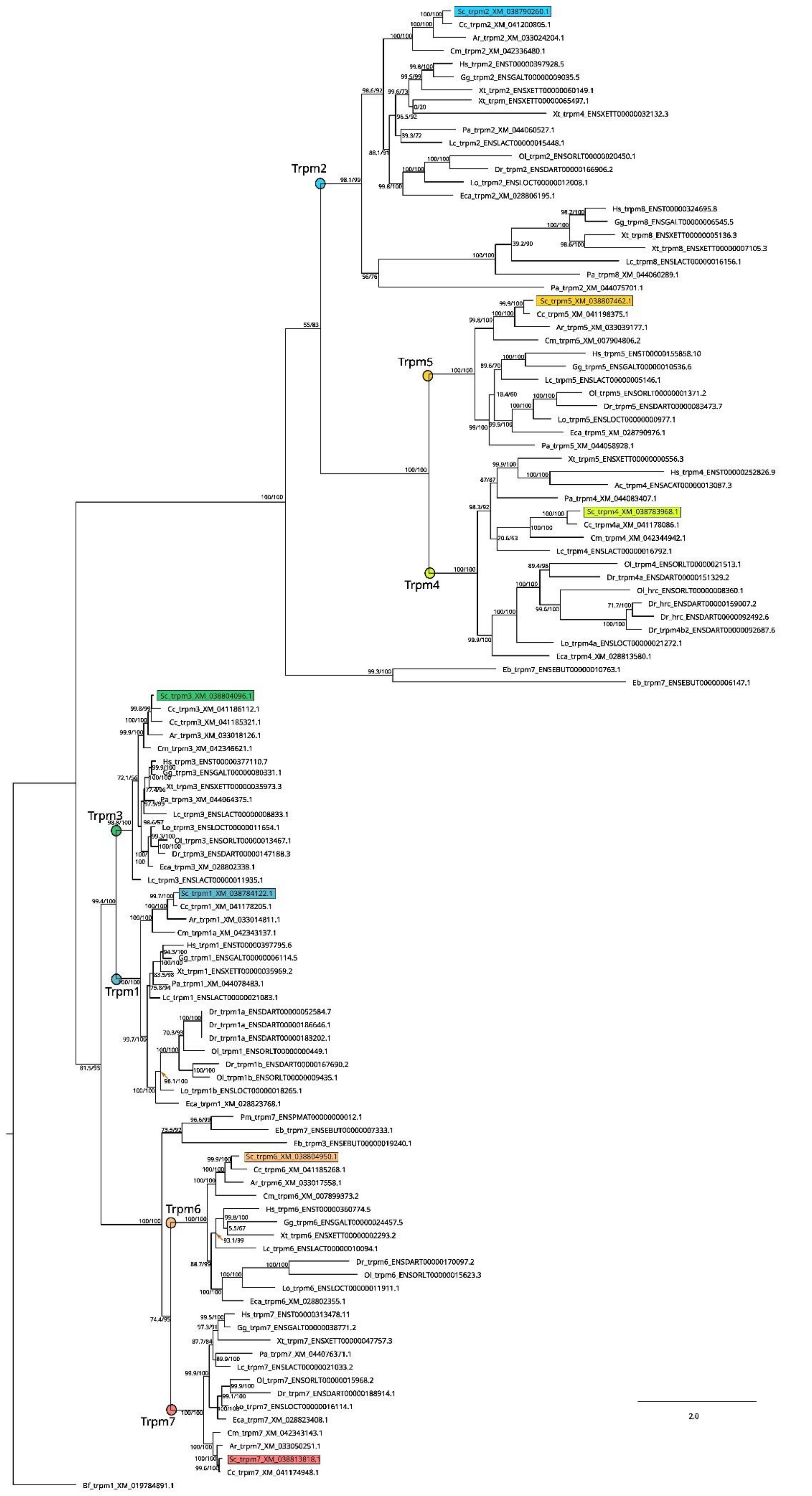

#### Supplementary Figure 31 Vertebrate Trpv gene tree inference by ML, rooted by an amphioxus sequence.

The alignment was 1205 positions, the best fit model was JTT+F+I+R7. Branch supports: SH-aLRT/ultrafast bootstraps (percentage); see main text for species names. The catshark sequences are highlighted in color, gnathostome orthology groups are identified by the position of bullets.

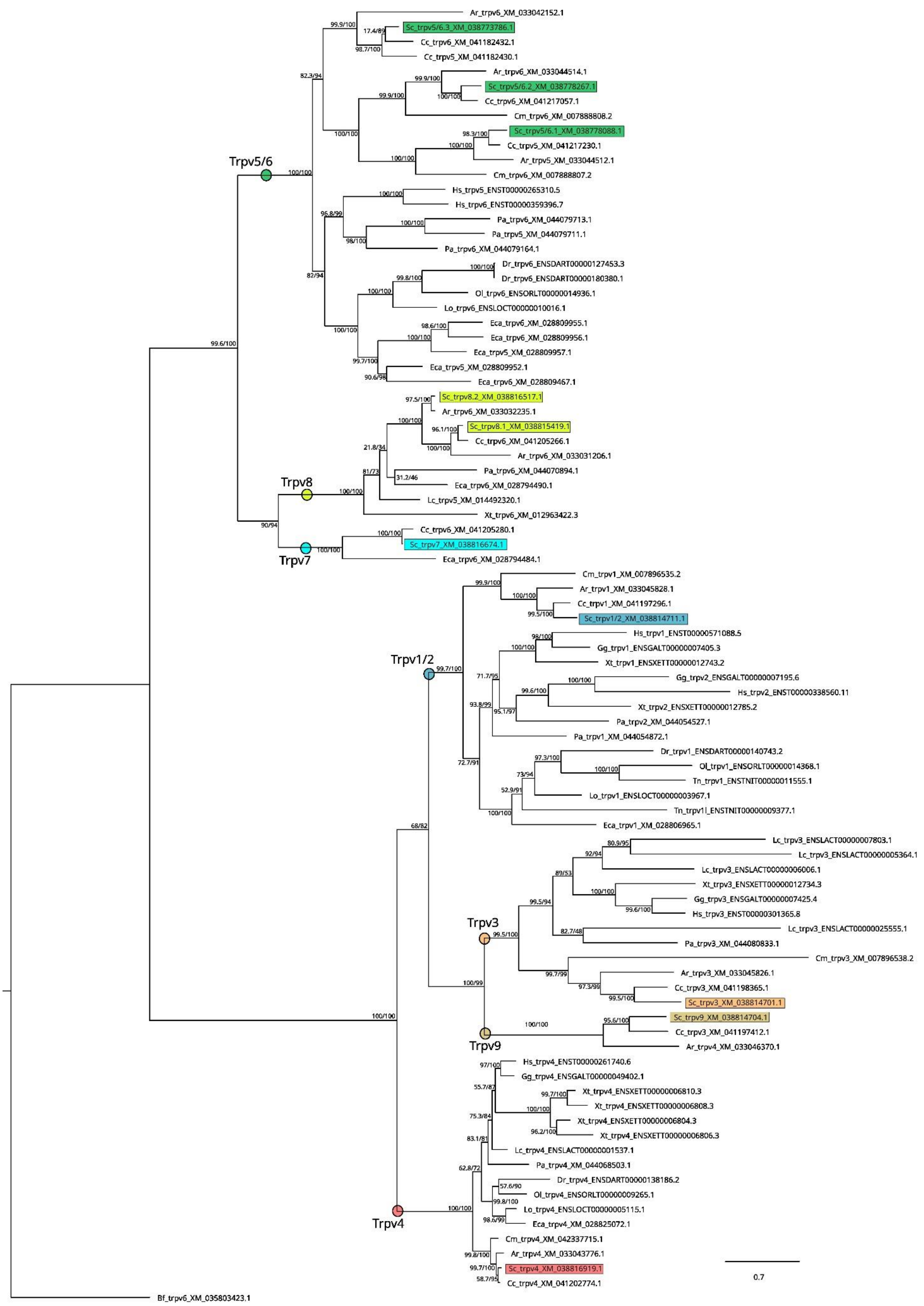

Supplementary Figure 32 Vertebrate Trpml (Mucolipin TRP; Mcoln) gene tree inference by ML, rooted by an amphioxus sequence.

The alignment was 748 positions, the best fit model was Q.plant+R5. Branch supports: SH-aLRT/ultrafast bootstraps (percentage); see main text for species names. The catshark sequences are highlighted in color, gnathostome orthology groups are identified by the position of bullets.

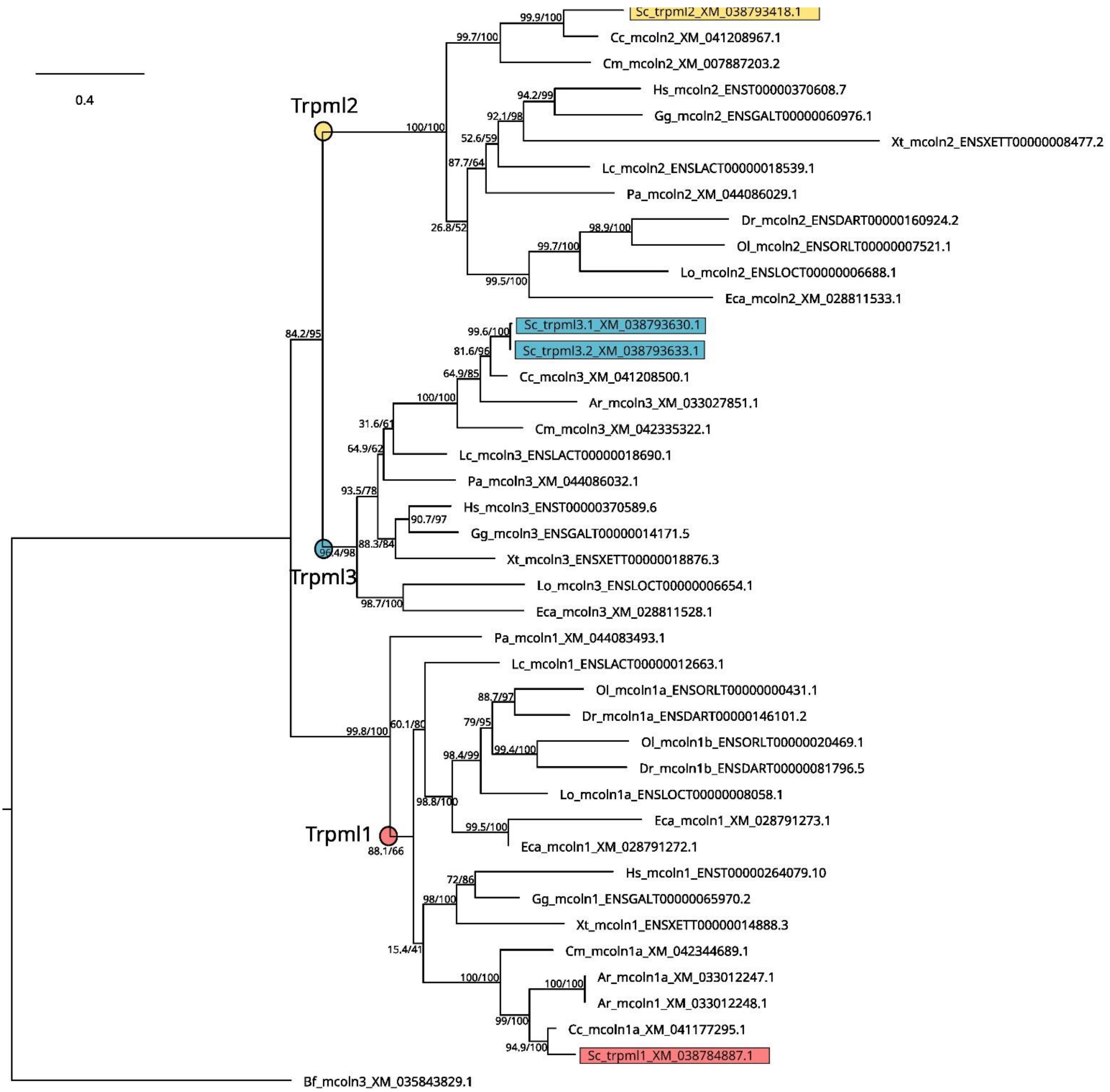

Supplementary Figure 33 Jawed vertebrate Pkd2-related gene tree inference by ML, rooted by cyclostome sequences. The alignment was 1083 positions, the best fit model was JTT+R6. Branch supports: SH-aLRT/ultrafast bootstraps (percentage); see main text for species names. The catshark sequences are highlighted in color, gnathostome orthology groups are identified by the position of bullets, gene nomenclature after England et al., 2017.

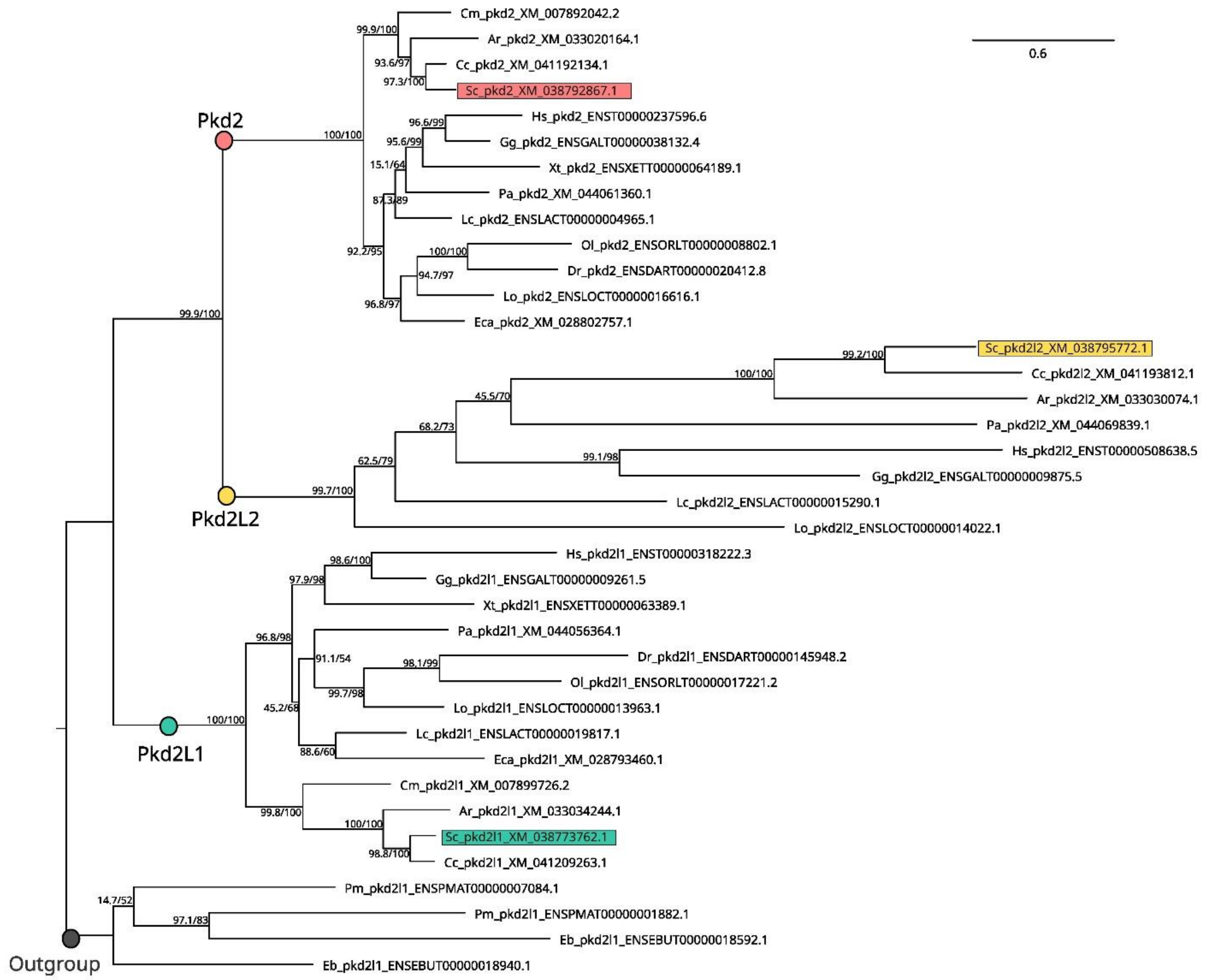

Supplementary Figure 34 Vertebrate Pkd1-related gene tree inference by ML, rooted by the vertebrate Pkdrej clade. The alignment was 6732 positions, the best fit model was JTT+F+R6. Branch supports: SH-aLRT/ultrafast bootstraps (percentage); see main text for species names. The catshark sequences are highlighted in color, gnathostome orthology groups are identified by the position of bullets. Very long branches have length divided by four in a clade colored brown.

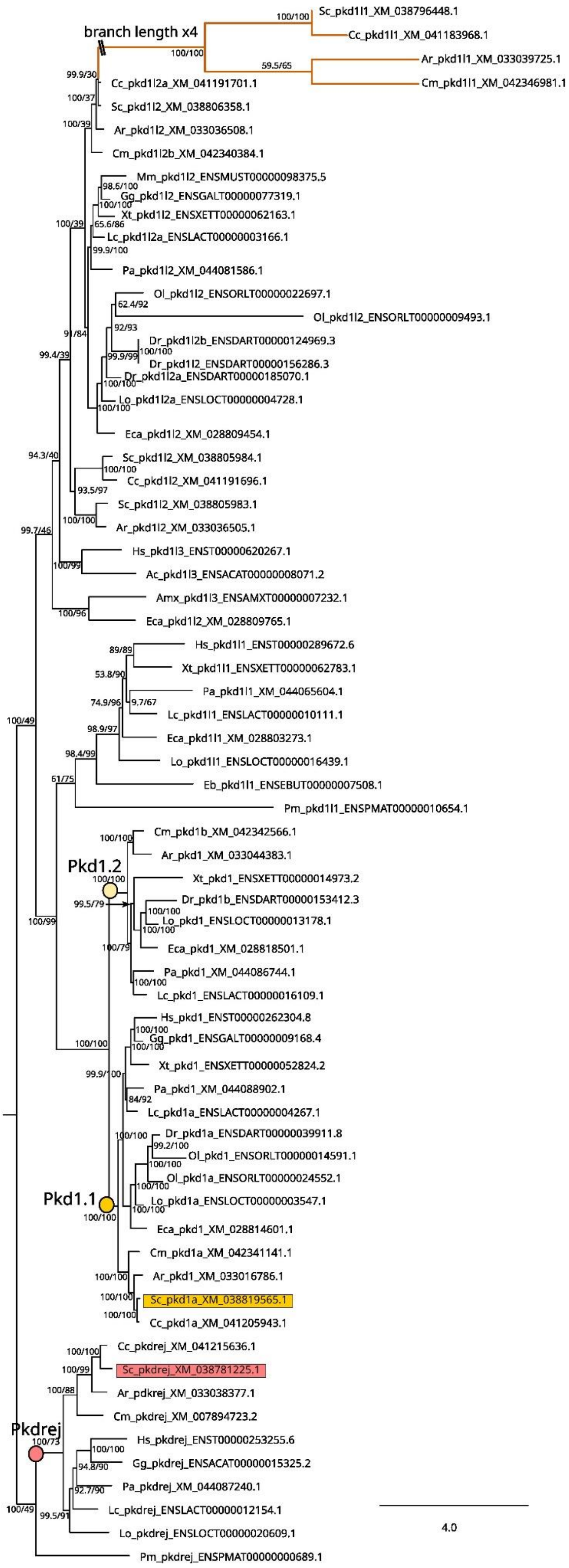

Supplementary Figure 35 Vertebrate Pkd1-related gene tree inference by ML, excluding the Pkd1 sequences, rooted by the Pkdrej clade.

The alignment was 6207 positions, the best fit model was JTT+F+R6. Branch supports: SH-aLRT/ultrafast bootstraps (percentage); see main text for species names. The catshark sequences are highlighted in color, gnathostome orthology groups are identified by the position of bullets.

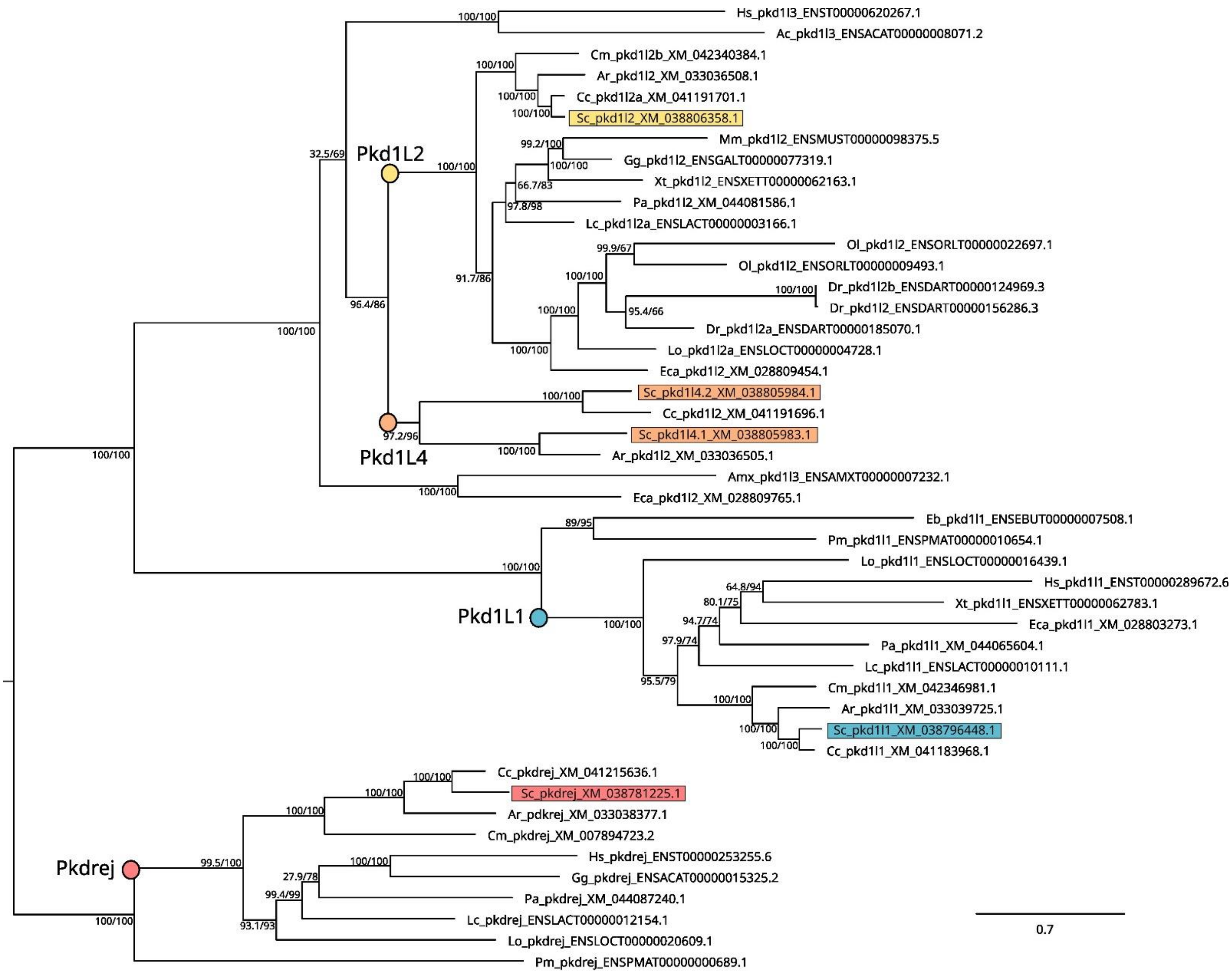

### SUPPLEMENTARY MATERIAL: Supplementary Text

Scripts for the successive steps conducted to construct the ncbi-utrs gene model reference are listed below:

**1. Initial construction of "isoform collapsed" version of gene models from NCBI data**

1.1. Selection of NCBI most supported isoform

1.2. Identification of the longest 3'UTR

BLAST command:

```
blastn -query transcrits_n.fa -db ./sc_gene_models_ncbi_isoform_collapsed_utrs_version_n_processed.fasta -outfmt '6 std qlen slen gaps' -max_target_seqs 1 -max_hsps 1 -evalue 1e-6 -num_threads 24 -out sortie_blastn_transcrits_n_vs_ncbi_collapsed.txt"
```

1.3. Appending of additional 3'UTR to NCBI most supported isoform

R code:

```
sortie = read.table("sortie_blastn_transcrits_n_vs_ncbi_collapsed.txt", header = F, fill = T, stringsAsFactors = F)

sortie = cbind(sortie, ifelse (sortie$V13 == "1/1", sortie$V14 - sortie$V8, sortie$V7 - 1))

sortie_ = sortie[((sortie$V10 >= sortie$V15 & sortie$V8 < sortie$V14 & sortie$V13 == "1/1") | (sortie$V9 >= sortie$V15 & sortie$V7 != 1 & sortie$V13 != "1/1")) & sortie$V3 == 100,]

group_sorted <- sortie_[order(sortie_$V2, -sortie_[,17]),]

group_sorted = group_sorted[!duplicated(group_sorted$V2),]

write.table(group_sorted, "sc_gene_models_ncbi_vs_version_n.txt", row.names = F, sep = "\t")
```

**2. Addition of divergent isoforms**

2.1 PERL code for the processing:

```
#!/usr/bin/perl

use strict;

use warnings;

sub main {

        my($contig, $line, $blast_line, $seq, $blast_contig, $n, $newseq, $utr, $frame);

        open(FHI,"./sc_gene_models_ncbi_isoform_collapsed_utrs_version_n_processed.fasta") or die ("$0 : cannot open file: $!");

        open(FHO, ">./sc_gene_models_ncbi_isoform_collapsed_utrs_version_n+1.fasta") or die ("$0 : cannot open file: $!");

        while ($line=<FHI>) {

                chomp($line);

                ($contig) = ($line =~ m/^>(\S*)/);

                if (defined($contig)) {

                        print "$contig\n";

                        $blast_line = `cat ./sc_gene_models_ncbi_vs_version_n.txt | grep $contig`;

                        chomp($blast_line);

                        if (length($blast_line) != 0) {

                                ($blast_contig) = ($blast_line =~ m/^"(\S*)"\s/);

                                ($n) = ($blast_line =~ m/\s(\S*)$/);

                                print "$blast_contig\n";
```

```
                              $seq = `module load bioinfo/samtools-1.8; samtools faidx
sc_gene_models_ncbi_isoform_collapsed_utrs_version_n_processed.fasta $contig`;

                              $newseq = `module load bioinfo/samtools-1.8; samtools faidx transcrits_n.fa
$blast_contig`;

                              $newseq =~ s/[\r\n]+//g;

                              $newseq =~ s/^(>.*\d){1}//;

                              ($frame) = ($blast_line =~ m/\s"(\d\/\S*)"\s/);

                              print "$frame\n";

                              ($utr) = ($frame eq "1/1") ? substr ($newseq, -$n) : substr ($newseq, 0, $n);

                              if ($frame ne "1/1") {

                                     $utr =~ tr/ACGTacgt/TGCAtgca/;

                                     $utr = reverse $utr;

                              }

                              print "$utr\n";

                              print FHO "$seq"."$utr"."\n";

                       }

                       else {

                              $seq = `module load bioinfo/samtools-1.8; samtools faidx
sc_gene_models_ncbi_isoform_collapsed_utrs_version_n_processed.fasta $contig`;

                              print FHO "$seq";

                       }

               }

       }

       close(FHO);

       close(FHI);

}

main();
```

2.2 File cleanup

```
#perl -pe '$. > 1 and /^>/ ? print "\n" : chomp' sc_gene_models_ncbi_isoform_collapsed_utrs_version_n+1.fasta
> sc_gene_models_ncbi_isoform_collapsed_utrs_version_n+1_processed.fasta
```

3 Inclusion of transcriptomic data

This step is analogous to steps 1 and 2, except for the following file cleanup at the end:

```
awk '/^>/ { print (NR==1 ? "" : RS) $0; next } { printf "%s", $0 } END { printf RS }'
sc_gene_models_ncbi_isoform_collapsed_utrs_version_n+1.fasta > sc_ncbi_utrs.fasta
```
